# Light-activated signaling in DNA-encoded sender-receiver architectures

**DOI:** 10.1101/2020.06.10.144162

**Authors:** Shuo Yang, Pascal A. Pieters, Alex Joesaar, Bas W.A. Bögels, Rens Brouwers, Iuliia Myrgorodska, Stephen Mann, Tom F.A. de Greef

## Abstract

Collective decision making by living cells is facilitated by exchange of diffusible signals where sender cells release a chemical signal that is interpreted by receiver cells. Biologists have started to unravel the underlying physicochemical determinants that control the effective communication distance using genetically modified cells. However, living systems are inherently challenging to manipulate and study systematically and quantitatively. Therefore, the development of generic and tunable abiotic mimics featuring compartmentalized signaling is highly desirable. Here, by adapting a previously reported artificial cell-cell communication system, we engineer DNA-encoded sender-receiver architectures, where protein-polymer microcapsules act as cell mimics and molecular communication occurs through diffusive DNA signals. We prepare spatial distributions of sender and receiver protocells using a microfluidic trapping array, and setup a signaling gradient from a single sender cell using light, which activates surrounding receivers through DNA strand displacement. Our systematic analysis reveals how the effective signal range of a single sender is determined by various factors including the density and permeability of receivers, extracellular signal degradation, signal consumption and catalytic regeneration. In addition, we construct a three-population configuration where two sender cells are embedded in a dense array of receivers that implement Boolean logic and investigate spatial integration of non-identical input cues. The results advance our understanding of diffusion-based sender-receiver topologies and present a strategy for constructing spatially controlled chemical communication systems that have the potential to reconstitute collective cellular behavior.

## Introduction

Collective behavior in cellular systems emerges from a tightly choreographed interplay between cellular communication and intracellular signaling programs^1^. Sender-receiver architectures, where sender cells secrete soluble signals which form a concentration gradient that is interpreted by receiver cells, are ubiquitous in biological systems^2^. Sender-receiver topologies allow cellular populations to collectively regulate key intracellular events such as cellular growth^3–5^, cell death^4^ and cell differentiation^6^, and orchestrate diverse multicellular functions such as tissue regeneration^7^, cell migration^8^, immune response amplification^9–12^ and robust positioning^13–17^ of cells within a tissue. The effective communication distance over which a single sender can propagate a soluble signal is determined by a number of physicochemical and biological determinants^18^. Previous work suggests that for many living sender-receiver systems this characteristic length-scale is in the order of 50-500 μm^7,19–21^ and can be modulated by diverse factors such as cell density^9^, signal diffusivity^18^, extracellular signal degradation^22^ and signal consumption^9^ by receiver cells. However, how each of these factors individually controls the signaling length scale remains unclear.

Synthetic sender-receiver architectures^2,23,24^ based on genetically engineered cells have emerged as excellent tools to establish population-level behaviors such as morphogen reconstitution^17,25^, artificial networks^26–29^, Boolean logic gates^30,31^ and pattern formation^32,33^, which all arise from the complex interplay of cell-cell communication and intracellular programs. However, quantitative analysis of sender-receiver systems in living cells is complicated by the large number of variables and context/pathway-specific responses of individual cells. Therefore, a generic platform that would allow quantitative spatiotemporal analysis of sender-receiver architectures remains to be developed. A promising approach to circumvent the use of real cells is to implement synthetic communication modules using protocells which serve as minimalistic model systems for living cells^34–37^. While current protocell-based systems do not display the rich information processing capabilities of living cells, their minimalistic design allows for rudimentary biological processes to be mimicked with high degrees of control over experimental conditions. Diffusion-mediated communication has been established in bead-based^38–41^ and more life-like protocell-based systems using range of soluble signaling factors such as small molecules^42,43^, proteins^44^ and DNA^45^. However precise control over signal release, downstream signal processing and protocell spatial positioning, which are prerequisites to engineer sender-receiver architectures, is currently lacking in these systems. Moreover, spatial patterns emergent from bead-based systems have been reported^40,46^ but macroscale reaction-diffusion (RD) patterns in protocell-based synthetic communication systems have not been achieved yet.

Previously, we have developed the general and scalable platform BIO-PC^45^ (Biomolecular Implementation of Protocellular Communication) capable of distributed inter-protocellular molecular communication through DNA strand displacement (DSD) reactions^47^. Proteinosomes^48^ are used as cell-like semi-permeable compartments, which contain localized DNA gate complexes and are permeable to short DNA strands that can initiate DSD reactions. Due to the excellent programmability and predictability of DSD, BIO-PC has great potential in mimicking key features of cell-cell communication in living systems. In the present work, we adapt the BIO-PC platform for implementation of diffusion-based sender-receiver architectures (Fig. 1a). We establish sender-receiver systems by sequential localization of a single sender and receiver protocells in a microfluidic trapping array. Spatially controlled signal release from the sender is initiated by light activation resulting in receiver cell activation by a diffusion-consumption mechanism^9^. Our work reveals how the corresponding signaling length scale depends on the density and permeability of receivers, signal consumption as a result of receiver activation, and extra-protocellular signal degradation. We then construct a sender-transceiver system where the diffusible signal responsible for activation of receivers is recycled by a fuel-driven catalytic DSD reaction and reveal an increase in signaling length scale. Finally, we establish a spatially encoded Boolean AND gate at the population level where the receiver population integrates non-identical signals released from two distant senders. Our spatial-controlled, DNA-encoded protocell system allows quantitative analysis of diffusion-based sender-receiver architectures and has the potential to uncover design principles of natural cell-cell communication modules.

**Figure 1.**
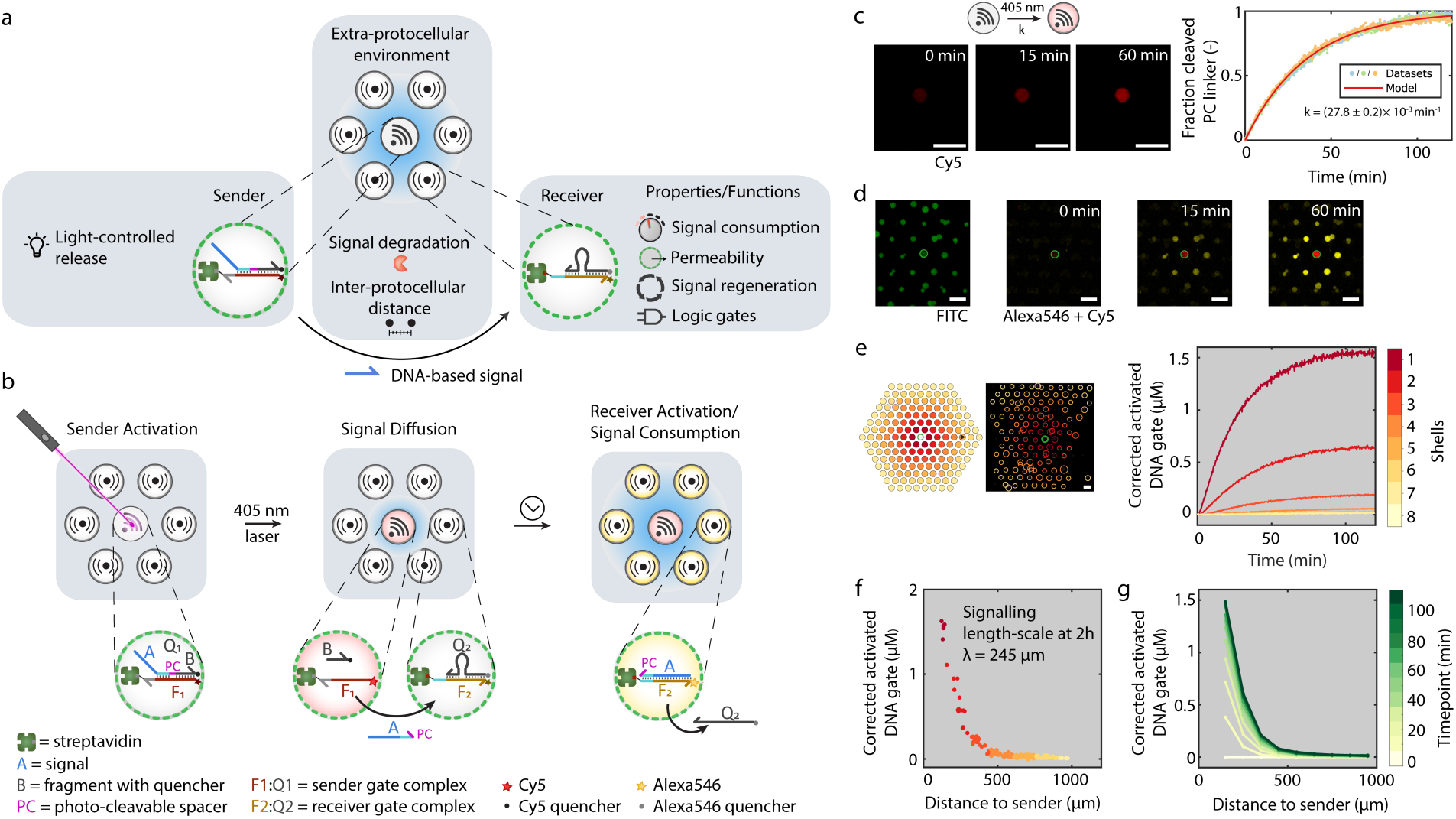
Light-activated DNA-encoded sender-receiver spatial system. **a** A single sender protocell and multiple receiver protocells are localized on a 2D spatial grid using a PDMS-based trapping array. Light-activated release of a ssDNA signal from the sender protocell sets up a signaling gradient which activates nearby receiver protocells. By controlling the characteristics of the protocells and environmental factors, this architecture enables quantitative analysis of diffusive signal propagation in space and time and programmable properties. **b** The sender protocell contains a fluorescently quenched internalized gate complex **F**_**1**_**Q**_**1**_ anchored to streptavidin using a biotinylated DNA gate strand (**F**_**1**_). Signal release from the sender protocell is triggered by laser irradiation resulting in photo-cleavage of strand **Q**_**1**_, concomitant dissociation of the two cleaved parts (**A** and **B**) and Cy5 fluorescence. Signaling strand **A** activates the surrounding receiver protocells by displacement of a quencher strand (**Q**_**2**_) from an internalized streptavidin-anchored gate complex **F**_**2**_**Q**_**2**_ to produce an Alexa546 fluorescence output and consumption of strand **A. c** Confocal micrographs of one sender protocell showing time-dependent increase in Cy5 fluorescence upon laser (405 nm) irradiation, indicating signal release. The plot shows the background-corrected fluorescence trace and exponential fit of the photo-cleavage reaction with a first-order rate constant of 0.0278 min^-1^. Experiments were performed in independent triplicates. **d** Confocal micrographs of one sender and multiple receivers (FITC-labelled proteinosome membrane, green) showing time-dependent increase in Cy5 fluorescence (red) and Alexa546 (yellow) fluorescence associated with signal release and activation, respectively. **e** Binning method employed to analyze spatial receiver activation. Protocells are binned in concentric shells based on their distance from the sender (left images). Plots (right) show time-dependent changes in the concentration of activated DNA gates associated with receiver protocells in different concentric shells arranged around the central sender. Shell **1** (dark red) is closest to the sender. **f** Plots of concentrations of activated DNA gates in individual receivers positioned at different distances from the sender protocell. Data collected after 2h of signal release. The color code corresponds to the different concentric shells as shown in (**e**). **g** Concentration of activated DNA gate complexes in receiver protocells plotted for different times as a function of distance to the sender. Sender protocells and receiver protocells were prepared using 10 μM and 4 μM streptavidin respectively. Experiments were performed at room temperature. All scale bars are 100 μm.

## Results and Discussion

### Construction of a light-activated spatial DNA-encoded sender-receiver system

We adapted the BIO-PC platform to function as a sender-receiver architecture by employing two types streptavidin-containing proteinosome-based semipemeable protocells which were loaded with biotinylated DNA gate complexes capable of sending or receiving short single-stranded DNA strands respectively (Fig. 1b). Our setup is based on preparation of multimodal protocell populations consisting of a single sender and multiple receiver protocells (average protocellular diameter 33.84 μm ± 5.99 μm, Supplementary Fig. 1) using a microfluidic trapping array. Diffusive molecular communication from the sender to surrounding receiver protocells is initiated by applying laser irradiation to the sender protocell, resulting in cleavage of an internalized photo-cleavable nitrobenzyl linker, and concomitant release of DNA strand **A**. Strand **A** functions as the diffusible signal that is secreted from the sender protocell and migrates through the medium thereby activating surrounding receiver protocells resulting in a fluorescent response which can be probed with high spatiotemporal resolution (*vide infra*). Specifically, the sender protocell contains an encapsulated DNA gate complex **F**_**1**_**Q**_**1**_ consisting of a fluorophore (Cy5)-labeled gate strand **F**_**1**_ and strand **Q**_**1**_ functionalized with a quencher and photo-cleavable nitrobenzyl moiety (Fig. 1B). Upon laser irradiation, strand **Q**_**1**_ is cleaved into two shorter ssDNA strands **A** and **B** which dissociate from the **F**_**1**_ strand at room temperature (Supplementary, Table. 1). We characterized photo-cleavage of the internalized **F**_**1**_**Q**_**1**_ gate complex by localizing a single sender protocell in the trapping array followed by irradiation with laser light, resulting in an increase in Cy5 fluorescence of the sender protocell over time due to the cleavage of **Q**_**1**_ and dissociation of the quencher-labeled fragment **B** (Fig. 1c and Supplementary Movie 1). The photo-cleavage process is observed to follow first order kinetics (Fig. 1c and Supplementary Fig. 2). Furthermore, photocleavage of the nitrobenzyl linker inside the sender protocell is localized to the illuminated area and a specific wavelength (Supplementary Figs. 3 and 4). Together, these results validate that the photo-cleavage of internalized DNA gate complexes inside a sender protocell can be achieved with a high spatial resolution.

Next, we assembled a multimodal sender-receiver population by sequential loading of a single sender and multiple (∼150) receiver protocells. The receiver protocells contain an encapsulated DNA gate complex **F**_**2**_**Q**_**2**_ consisting of a fluorophore (Alexa 546)-labeled gate strand **F**_**2**_ with an exposed toehold domain and a quencher-labeled strand **Q**_**2**_ (Fig. 1b) resulting in quenching of the Alexa fluorescence. Activation of the receiver protocells is initiated by toehold-mediated strand displacement of **Q**_**2**_ by signal strand **A** released from the sender protocell. Experiments were initiated by laser irradiation (405 nm laser, 2h) of the single sender protocell resulting in photo-cleavage of the **F**_**1**_**Q**_**1**_ gate complex which could be monitored by an increase in Cy5 fluorescence (Fig. 1d, red). We confirmed successful activation of the receiver protocells by signal strand **A** by monitoring an increase in Alexa546 fluorescence in individual receivers (Fig. 1d, yellow, and Supplementary Movie 2). To analyze receiver activation dynamics under the signaling gradient we binned receivers in concentric shells based on their radial distance from the sender and plotted the average fluorescence traces (Fig. 1e). Receivers in close proximity to the sender protocell are activated first and to a higher extent, confirming that the diffusible signal released from the central sender is consumed by the surrounding receivers, which therefore limits the activation of receivers at larger distances from the sender. To further quantify the spatiotemporal data, we defined a characteristic signaling length scale λ, as the average distance that signaling strands can diffuse from the sender and activate the surrounding receivers (Supplementary Methods and Supplementary Fig. 5). We calculated the length scale λ by plotting the receiver activation as a function of distance after 2h of illumination and found a λ value of approximately 245 μm (Fig. 1f). This value is well in the range of many natural systems that communicate via soluble factors, i.e. morphogens^49^ (40∼200 μm), cytokines^9^ (30∼150 μm) and retinoic acid^19^ (300∼500 μm). The spatial distribution of the receiver cells’ activation states at different timepoints is shown in Fig. 1g.

To validate the experimental observations, we performed 2D reaction-diffusion simulations of the sender-receiver population using the Visual DSD software^45,50,51^. The numerical model was parametrized using the average signal release rate constant, DSD rate constant and membrane permeability obtained in separate experiments (Supplementary Methods). The obtained activation dynamics and signaling length-scale (Supplementary Fig. 6) are in agreement with the experimental results. Collectively, these results establish that spatially controlled light-induced signal release from an individual sender protocell results in a distance-dependent activation of surrounding receivers in agreement with a diffusion-consumption mechanism.

### Modulation of signaling range

Intercellular communication distances established by soluble factors in multicellular populations are regulated by both internal and external physicochemical factors, such as signal secretion, diffusion and consumption rates^11,14^. How each of these determinants modulates the signaling length scale has remained difficult to analyze due to the intrinsic complexity of natural sender-receiver systems. Here we employ our synthetic sender-receiver architecture and quantitatively analyze the contribution of individual determinants to the signaling length scale. Specifically, we constructed multimodal sender-receiver populations through sequential loading of a single sender and multiple (100-150) receiver protocells. Using this setup, we determined the internal and external determinants leading to changes in effective signaling length scale associated with variations in the capacity and rate of signal consumption, and levels of signal degradation in the environment (Fig. 2a).

**Figure 2.**
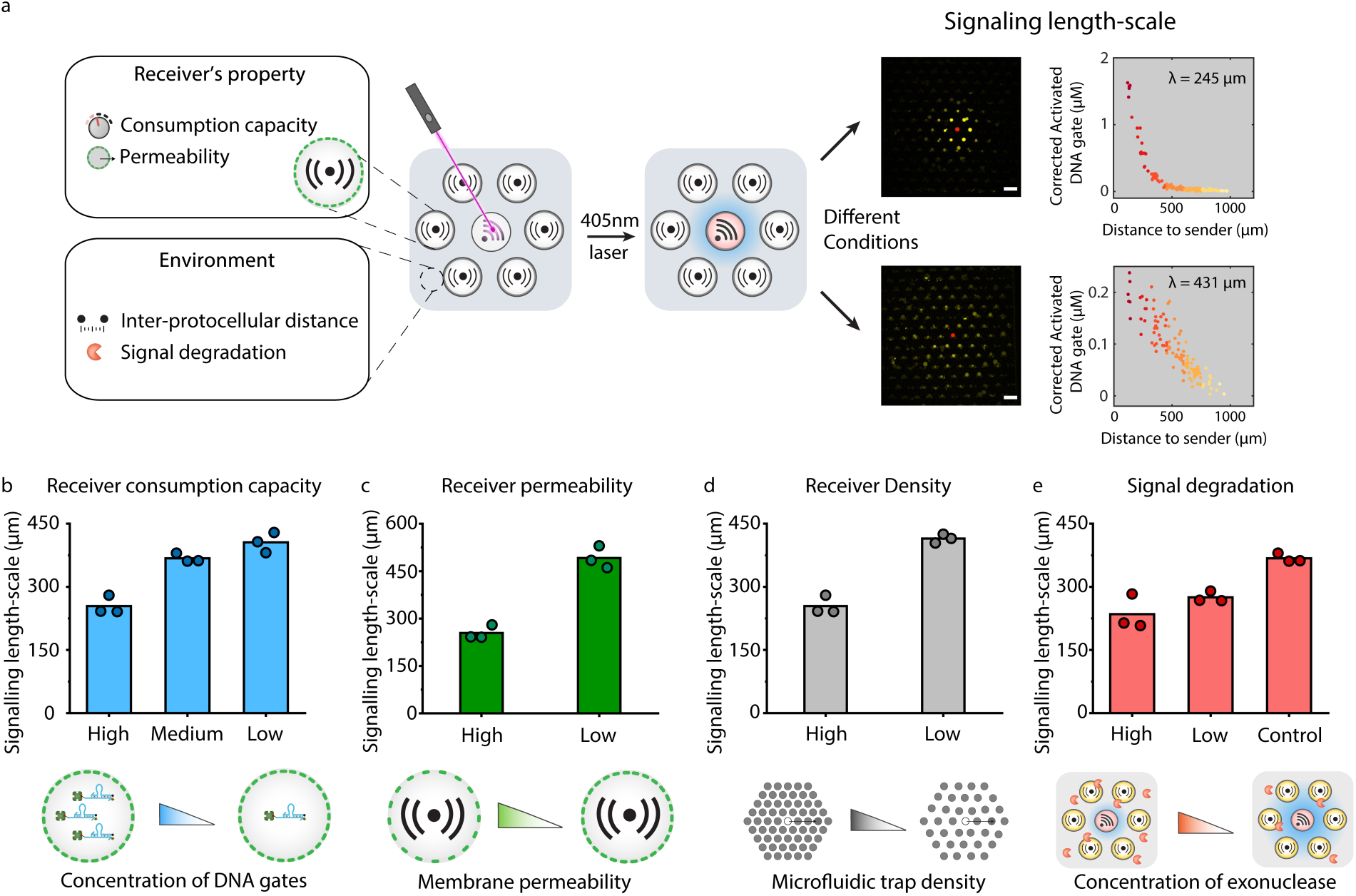
Tuning of the signalling length scale in light-activated DNA-encoded sender-receiver spatial systems. **a** Changes in internal factors such as signal consumption capacity (receiver DNA gate complex concentration) and consumption rate (receiver membrane permeability) and external factors such as inter-protocellular distance (protocell trap density) and signal degradation (exonuclease concentration) influence the signaling length scale (left and centre). Plots show typical experimental data used for the determination of the signaling length scale (right). Data collected after 2h of signal release from the sender. Scale bars 150 μm. **b-e** Modulation of signaling length scale at t = 2 h for changes in receiver consumption capacity (**b**), consumption rate (membrane permeability) (**c**), protocell trap density (**d**) and signal degradation (**e**). High, medium and low levels of the receiver protocell-entrapped DNA gate complex relate to changes in receiver-encapsulated streptavidin concentrations ([SA]) of 4, 1 and 0.4 μM, respectively (**b**). High (202.8 μm min^-1^) and low (2.16 μm min^-1^) receiver permeabilities relate to modifications in the protocell membrane crosslinking density; [SA] = 1 μM (**c**). High and low receiver number densities relate to the use of 90 or 70 microfluidic traps per mm^2^, respectively; [SA] = 4 μM (**d**). High and low levels of signal degradation arise from the presence of 0.1 unit/μL and 0.05 unit/μL of exonuclease I, respectively; [SA] = 1 μM. The control experiment is performed in the absence of exonuclease I (**e**). For all experiments, the concentration of encapsulated streptavidin in the sender protocells was 10 μM. All experiments were performed in independent triplicates at room temperature. All the experimental conditions are summarized in Supplementary Table 2.

In cellular populations, binding of soluble factors to receptors on neighboring cells results in consumption of the available signal and therefore influences the effective signaling range^11,14,18^. In the BIO-PC platform, the consumption capacity of individual receiver protocells can be varied by changing the concentration of encapsulated **F**_**2**_**Q**_**2**_ DNA gate complex, which depletes the diffusible signal by strand-displacement. We performed the sender-receiver experiments using receiver protocells with three different DNA gate complex concentrations (Supplementary Fig. 7) and calculated the corresponding signaling length-scales (Fig. 2b and Supplementary Figs. 5 and 8-15). In general, for the 20-40 μm sized receiver protocells used in this study, the effective signaling length scale ranges between 200-500 μm. In agreement with our expectations, increasing the consumption capacity of individual receiver protocells results in lower effective communication distances due to a higher local depletion of the soluble signal. Besides consumption capacity, the consumption rate of a soluble signal can also modulate the signaling length scale. In multicellular populations, the consumption rate of a morphogen or cytokine can be regulated by controlling the rate of endocytosis^11^. Here, we modulate the consumption rate by tuning the membrane permeability of receiver protocells. We have previously shown that the permeability (P) of proteinosomes can be tuned using protein-crosslinking reagents of different length and revealed how the permeability influences the compartmentalized DSD reaction kinetics^45^. We prepared high-P and low-P receiver protocells (Supplementary Methods), quantified their permeability and confirmed they have approximately similar binding capacity for biotin-labelled DNA (Supplementary Fig. 16). As expected, the experimentally derived signaling length scale is higher for low-P receiver protocells (Fig. 2c and Supplementary Figs. 10-12 and 17-19) as the soluble signal is consumed at a lower rate.

Because an individual sender is surrounded by multiple receivers, the effective signaling distance not only depends on the consumption rate and consumption capacity of individual receivers but also on the cumulative signal consumption which can be varied by modulating the protocell number density in the spatial array^9^. We fabricated microfluidic trapping arrays with two different densities of protocell traps and determined the effective signaling range from the experimental data (Supplementary Fig. 20). Our data shows that the signalling length scale increases when protocell density is decreased (Fig. 2d and Supplementary Figs. 4, 8-9 and 21-23). This increasing communication distance can be explained by lower total signal consumption capacity as the number density of receiver protocell decreases.

Biochemical degradation of diffusible factors is a key regulatory mechanism in morphogenesis and can control both the signaling range and sharpness of the diffusion front^14,16,52^. To mimic signal degradation, we added exonuclease I (3’ to 5’) to the trapped proteinosomes before initiating the photo-cleavage reaction. Exonuclease I selectively degrades the diffusible signal from its free 3’ end. Control experiments using proteinosomes containing an encapsulated 3’ fluorescently-labeled ssDNA show that exonuclease is capable of diffusing across the proteinosome membrane (Supplementary Fig. 24) resulting in the presence of exonuclease inside and outside the single sender and multiple receiver protocells. However, the encapsulated DNA gate complexes in the sender and receivers lack a free 3’ end preventing their degradation (Supplementary Fig. 25). Laser-irradiation of the sender protocell cleaves the internalized DNA gate complex and yields a diffusible signal strand with a free 3’ end that is amenable for exonuclease degradation. Because the 3’ end of the diffusible signal contains the toehold binding domain required for strand-displacement with the receiver gate complex, enzymatic degradation strongly inhibits receiver activation. We performed sender-receiver experiments for two different concentrations of exonuclease I added to the medium (Fig. 2e) and calculated the signaling length scale from the experimental data Supplementary Figs. 10-12 and 26-31. As expected, higher concentrations of exonuclease result in decreasing signaling length scales. Taken together, these findings reveal how a fully synthetic sender-receiver protocell platform can be used to systematically study the effect of isolated physicochemical factors on the diffusive communication range.

### Signal regeneration

In living cells, intercellular amplification of soluble signaling molecules is a ubiquitous mechanism employed to direct and control downstream cell fate decisions^53,54^. Here we implement non-enzymatic DNA-based catalytic reaction networks^55^ and realize inter-protocellular signal amplification by engineering the sender-receiver architecture into a sender-transceiver system, where transceiver protocells can be activated by the diffusible signal but are also capable of regenerating this signal in the presence of excess fuel strand (Fig. 3a). Similar to the sender-receiver experiments, sender protocells contain encapsulated DNA gate complex **F**_**1**_**Q**_**1**_. Transceiver protocells contain encapsulated DNA gate complex **F**_**3**_**Q**_**3**_ where the streptavidin-bound biotinylated strand **F**_**3**_ is labeled with Alexa546 while strand **Q**_**3**_ is modified with a quencher. Laser irradiation triggers the release of signal **A** from the sender protocell, which activates the transceiver gate complex by displacing **Q**_**3**_. The activated transceiver gate complex **F**_**3**_**A** exposes the toehold on the **F**_**3**_ strand, allowing an abundantly present fuel strand to bind **F**_**3**_ and regenerate signal strand **A**. We prepared a bimodal protocell population consisting of a single sender and multiple receivers (∼150) by sequential loading of protocells in the trapping device. Next, the trapping chamber was filled with fuel strand to a final concentration of 0.1 μM and the experiment was initiated by laser irradiation (405 nm, 2 h) of the sender protocell. Comparison of the fluorescent micrographs obtained in the presence and absence of fuel (Fig. 3b) clearly reveal transceiver activation at larger distances from the sender in the presence of the fuel, indicative of successful recycling of the diffusing signal. To further characterise sender-transceiver protocell communication, we plotted the transceiver activation dynamics for increasing distances from the sender (Fig. 3c) and the response time of each transceiver (Fig. 3d). We observed significant higher activation of transceivers at short distances from the sender in the presence of fuel while a significant higher fraction of transceivers is activated at larger distances from the sender (Supplementary Figs. 32-37). Importantly, a control experiment in which the individual sender protocell was not irradiated displays low leakage in the presence of fuel (Fig. 3c, black line), which is characteristic for catalytic DSD systems^55^. We also computed the characteristic length scale in the presence and absence of the fuel which reveals a higher signaling range as a result of regeneration of the soluble signal by fuel-driven DSD cycles (Fig. 3e). Simulations using a 2D reaction-diffusion model that incorporates the effect of the fuel-driven signal regeneration confirms this experimental observation (Supplementary Fig. 38). Collectively, these results show that signal regeneration can be integrated into spatial-controlled sender-receiver architectures and leads to an increase in the signaling length scale.

**Figure 3.**
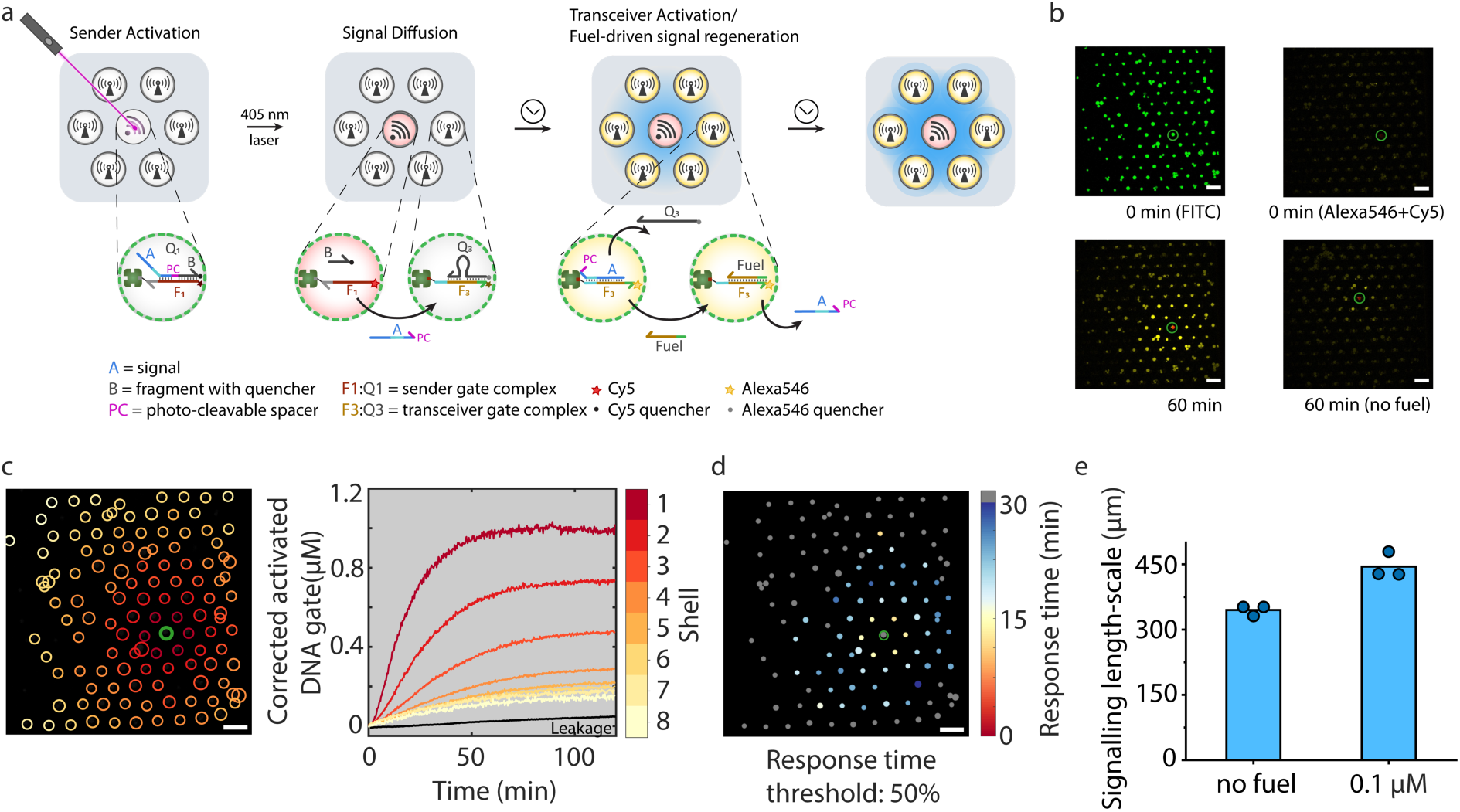
Signal regeneration in a light-activated spatial DNA-encoded sender-transceiver system. **a** The encapsulated sender gate complex is identical to that used for the sender-receiver system. Upon laser irradiation, signal strand **A** is released to generate Cy5 fluorescence and activates the surrounding transceiver protocells by displacement of a quencher strand (**Q**_**3**_) from encapsulated gate complex **F**_**3**_**Q**_**3**_ to produce Alexa546 fluorescence and consumption of strand **A.** After the initial response, a non-enzymatic DNA catalytic reaction recycles the signal strand **A** by consuming a fuel strand **b** Confocal micrographs of one sender and multiple transceivers (FITC-labelled proteinosome membrane, green) recorded at t = 0 (top left) showing minimal Cy5 and Alexa546 fluorescence before signal activation (top right) in the presence of a fuel strand; corresponding images 60 min after light-induced signal generation show increases in Cy5 fluorescence in the sender (red, release of signal **A**) and Alexa546 fluorescence (yellow, activation of **F**_**3**_**Q**_**3**_) in the surrounding transceivers (bottom left). The control experiment was performed without fuel and shows lower levels of Alexa546 fluorescence after 60 min due to the absence of signal regeneration (bottom right). **c** Binning of transceiver activation (left) and corresponding time-traces within different concentric shells for changes in the concentration of transceiver DNA gate activation (right). Shell **1** (dark red) is closest to the sender. To analyze spontaneous triggering of the transceiver gates by the fuel, the mean concentration of activated transceiver gate without signal release (i.e. no laser irradiation) is also plotted (black line). **d** Spatial barcode image of response time of transceiver protocells. The response time is defined as the time in which a transceiver reaches 50% of its final activated concentration; short ‘on’ time (red), long ‘on’ time (blue). To remove background noise, any protocells with an absolute increase less than 20 RFU are excluded and labelled with gray (n.d.) **e** Calculated signaling length-scales in a sender-transceiver system. Sender and transceiver protocells are prepared using 10 μM and 1 μM streptavidin respectively. The concentration of fuel was 0.1 μM. All experiments were performed in independent triplicates at room temperature. Scale bars, 150 μm.

### Spatial integration of diffusible signals by Boolean receivers

Spatial integration of chemical signals by Boolean operations is essential to generate collective behaviour in multicellular populations as exemplified by the immune and nervous systems^56,57^. Although Boolean RD systems have been implemented using the Belousov-Zhabotinsky reaction^58-62^, a versatile and tunable platform based on biomolecular reactions is currently lacking. Here, we reveal how the BIO-PC platform can be employed to integrate non-identical gradients by localized Boolean operations. Using a sequential loading procedure, we implemented a three-population configuration consisting of two non-identical sender protocells embedded in a high density of receivers that implement Boolean AND logic (Fig. 4a). The two senders contain gate complexes **F**_**4**_**Q**_**4**_ and **F**_**5**_**Q**_**5**_, respectively, which upon simultaneous laser illumination (405 nm, 1.8 h), secrete two distinct Cy3-labeled signal strands **A**_**2**_ and **A**_**3**_ as monitored by the decrease in Cy3 fluorescence of the sender protocells (Fig. 4b yellow). Receiver protocells contain an encapsulated DNA gate complex which is activated by a cooperative hybridization mechanism^63^ (Supplementary Fig. 39) where both **A**_**2**_ and **A**_**3**_ need to be simultaneously present to release quencher labelled strand **Q**_**6**_ resulting in an increase in Quasar 670 fluorescence (Fig. 4b, red). We analyzed spatiotemporal AND-gate receiver activation by binning receiver protocells into shells based on the maximum of the two distances to the senders and calculated the average fluorescence per bin over time (Fig. 4c and Supplementary Fig. 40). The experimental curves reveal that activation of the receivers is initiated at positions equidistant to the two senders, in agreement with AND-type logic. Our observations are also supported by 2D RD simulations using realistic parameters (Supplementary Fig. 41). Furthermore, we plotted the response time of each receiver as a function of the distance to both senders and find the lowest response time for receivers at equidistant position of both senders (Fig. 4d). Together, these results indicate that receiver protocells are activated by two non-identical signaling gradients of distributed spatial origins and demonstrate Boolean AND logic.

**Figure 4.**
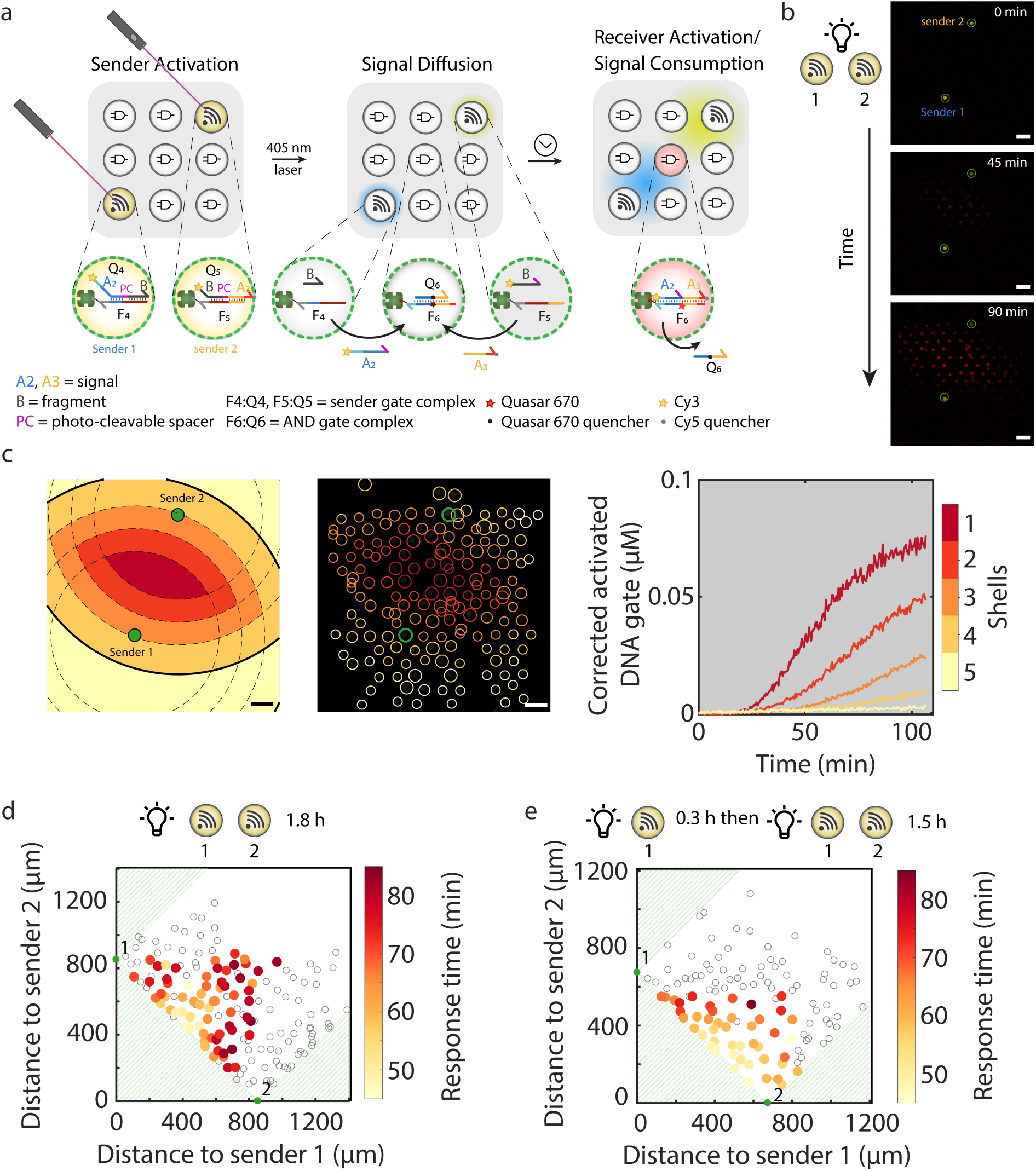
Spatial integration of non-identical signals by 2D-arrayed AND-gate protocells. **a** Two fluorescent sender protocells (**1** and **2**) containing internalized gate complexes **F**_**4**_**Q**_**4**_ or **F**_**5**_**Q**_**5**_, respectively, are embedded in a high number density of non-fluorescent AND-gate receivers. Signal release from sender protocells is triggered by laser irradiation resulting in photocleavage of **Q**_**4**_ and **Q**_**5**_, concomitant dissociation of the cleaved parts, **A**_**2**_+**B** and **A**_**3**_+**B**, and loss of Cy3 fluorescence. The Cy3-labelled dissociated strand **A**_**2**_ and non-fluorescent strand **A**_**3**_ activate Quasar670-quenched receiver protocells containing an encapsulated AND gate (**F**_**6**_**Q**_**6**_) through cooperative hybridization^63^ to produce a Cy3/Quasar670 fluorescence output. **b** Confocal micrographs of two sender protocells (**1** and **2**) and multiple AND gate receivers recorded at t = 0 (top) showing Cy3 fluorescence in the spatially separated transmitters. Light-induced activation leads to a reduction in Cy3 fluorescence (yellow) in the senders, and progressive increase in Quasar 670 fluorescence (red) associated with activation of receiver protocells. **c** Binning method used to analyze spatiotemporal activation of receiver protocells. Protocells are binned based on the maximum of the two distances to the senders, which yields bins with outer bounds that are the intersection of the equivalent bounds of single sender systems, as illustrated by the black lines (left). Corresponding time traces of AND gate receiver activation within different bins. Shell **1** (dark red) is closest to the two senders. **d** Response time of individual Boolean receivers upon simultaneous laser irradiation of two senders for 1.8 h plotted as a function of their distance to each of the two senders. The response time is defined as the time taken for an individual receiver to reach 50% of its final activated concentration. Two senders are marked as **1** and **2** in green. To remove background noise, any protocells with an absolute increase less than 1 RFU are excluded and labeled in hollow circles. (**e**) Response time of individual Boolean receivers upon sequential laser irradiation showing spatial bias in activation of receiver protocells Sender **1** is irradiated for 18 min, followed by irradiation of both senders for 1.5 h. Sender protocells and receiver protocells were prepared using 30 μM and 1 μM streptavidin respectively. Scale bars 150 μm. Experiments were performed at room temperature.

Because receiver protocells are activated by gradients from both senders, we wondered if we could spatially control initiation of receiver activation by sequential laser irradiation of the two senders. This would result in the development of a signal gradient from one of the two senders before the other gradient is established. Due to the reversible nature of the cooperative DSD mechanism^63^, signal strands from one sender that bind to the AND-gate are preferentially released in the absence of the other signal, preventing signal consumption by receivers. We first irradiated sender **1** for 18 minutes, followed by exposure of both senders for 1.5 h and calculated the response time for each receiver (Fig. 4e and Supplementary Fig. 42). We observe a skewed activation pattern where receiver activation is initiated in receivers in close proximity to sender **2**, in agreement with the presence of spatial bias in the established signal gradients. Together, these results show that a population of protocells could be programmed to integrate two non-identical cues and perform spatially-encoded Boolean operations using encapsulated DSD based reactions.

## Conclusion

Cellular communication by soluble factors allows populations of cells to coordinate their behavior. Secretion of diffusible messengers is often spatially localized resulting in local gradients near sender cells and the emergence of spatial niches characterized by a high concentration of a specific signal^9^. For non-migrating, micrometer-sized cells the characteristic signaling length scale of these gradients, i.e. the distance over which diffusive communication persists, appears to be around 50-500 μm^9,19,49^ and can be dynamically adjusted by competition between diffusion and signal consumption by receiver cells^9^ and active signal degradation^15^. In this work, we demonstrate a fully synthetic soft matter system based on semipermeable microcompartments that communicate via short DNA strands under a light-induced local signaling gradient arising from a single sender protocell. We prepared multimodal protocell arrays consisting of a single sender protocell and a polydisperse receiver population using a microfluidic trapping device and systematically quantified how individual parameters control the signaling length scale typically between 200 and 500 μm. The simplicity of the system allows variation of the consumption capacity and consumption rate of receiver protocells and introduction of active signal regeneration and degradation pathways. As a further showcase of our cytomimetic technology, we revealed how two local signaling gradients can be spatially integrated by employing receiver protocells containing Boolean AND gates.

For future research, we envision that the BIO-PC platform could form the basis for implementing a deterministic cellular automaton based on chemical reaction-diffusion networks which would be able to perform universal computation and permanently store the chemical state of each protocell^64^. In order to construct a DNA-based cellular automaton, additional DNA-based Boolean operations such as NOR, XOR and NAND gates, which have been shown to work in bulk^65^, need to be introduced into the protocell platform. In addition, the position of each protocell should be independently addressable, which could be achieved using either acoustic^66^ or magnetic^67^ driven manipulation. The development of minimally synthetic cellular communities with programmable communication protocols is a key goal in bottom-up synthetic biology^68^ and has the potential to inform the design rules of collective decision making in multicellular populations.

## Materials and Methods

Streptavidin-containing proteinosomes were prepared similarly to our previously reported procedures^45^. All DNA sequences are listed in Supplementary Table 3. DNA oligonucleotides were purchased from Integrated DNA Technologies and Biosearch Technologies. A two-layer PDMS microfluidic chip was produced using standard soft lithographic techniques^69^. To perform an experiment, protocells were first suspended in the buffer solution and delivered to the trapping chamber in the microfluidic device by a compressed air line. Photo-cleavage of gate complexes inside a sender protocell was triggered by 405 nm laser from a confocal laser scanning microscope (Leica SP8). The confocal microscope was also used to measure fluorescence of protocells in the trap array. Data analysis and 2D reaction-diffusion simulation were performed using Matlab and Visual DSD. Full details are given in Supplementary.

## Acknowledgments

We thank Pulin Li and Michael B. Elowitz for sharing Matlab codes used in our data analysis, Andrew Phillips from Microsoft Research for advice on Visual DSD. S.Y. and T.F.A.d.G. were supported by an NWO-VIDI grant from the Netherlands Organization for Scientific Research (NWO, 723.016.003); P.A.P., A.J. and T.F.A.d.G. were supported by an ERC starting grant by the European Research Council (project n. 677313 BioCircuit); B.W.A.B. was supported by Microsoft Research (PhD Scholarship Programme); I.M. and S.M. were supported by the European Commission (ERC Advanced Grant, 740235).

