## Supplementary Information for "Light-activated signaling in DNA-encoded sender-receiver architectures"

**Yang et al**

**This PDF file includes:**

Supplementary Materials and Methods

Supplementary Note

Figures. 1 to 44

Tables 1 to 6

Legends for Movies 1 to 2

Supplementary References

### Supplementary Materials and Methods

#### Materials

2-Ethyl-1-hexanol (Sigma, 98%), 1-(3-dimethylaminopropyl)-3-ethylcarbodiimide HCl (EDC, Carbosynth), 1,6-diaminohexane (Sigma, 98%), Fluorescein isothiocyanate (FITC, 90%, Sigma), PEG-bis(N-succinimidyl succinate) (Mw= 2000, Sigma), BS(PEG)5(Sigma), streptavidin from *Streptomyces avidinii* (Sigma), bovine serum albumin (heat shock fraction, pH 7, ≥98%, Sigma), AZ 40xt (MicroChemicals), SU8 3050 (MicroChemicals), polydimethylsiloxane (PDMS, Sylgard 184). All other reagents were purchased from Sigma Aldrich.

#### Preparation of BSA-$NH_2$/PNIPAAm nanoconjugates

Cationized bovine serum albumin (BSA-$NH_2$) is synthesized via a previously reported procedure[1]. Typically, 1.5 g diaminohexane is dissolved in 10 mL MilliQ water, the resulting solution is adjusted to pH 6.5 using 5 M HCl and added dropwise to a solution of BSA (200 mg in 10 mL MilliQ water). The coupling reaction is initiated by addition of 100 mg of 1-(3-Dimethylaminopropyl)-3-ethylcarbodiimide HCl (EDC) immediately and another 50 mg after 5 h. The pH value of the solution is readjusted to pH 6.5 after EDC is added. The reaction mixture is stirred at 300 rpm using a hotplate stirrer and held at ambient temperature. The total reaction time is 11 h and then the solution is centrifuged to remove precipitate. The corresponding supernatant is dialyzed (Medicell dialysis tubing, MWCO 12-14 kDa) in MilliQ water overnight and freeze-dried.

Mercaptothiazoline-activated PNIPAAm ($M_n$ = 9800 g mol$^{-1}$, 4 mg in 5 mL MilliQ water) is synthesized via a previously reported procedure[1] and added to a stirred solution of BSA-$NH_2$ (10 mg in 5 mL of PBS buffer, pH 8.0). The reaction mixture is stirred for 10h at ambient temperature. Then the solution is purified using a centrifugal filter (Millipore, Amicon Ultra, MWCO 50 kDa) and freeze-dried.

Fluorescent-labeled nanoconjugates BSA-$NH_2$/PNIPAAm are synthesized via the same method, except that Fluorescein isothiocyanate (FITC) labeled BSA[1] is used as starting material.

#### Preparation of streptavidin-containing proteinosomes

Typically, BSA-$NH_2$/PNIPAAm nanoconjugates (final concentration 15 mg mL$^{-1}$), FITC-labeled BSA-$NH_2$/PNIPAAm nanoconjugates (final concentration 1 mg mL$^{-1}$), streptavidin (final concentration: 0.4, 1, 4, 10 µM) and 1.5 mg PEG-bis(N-succinimidyl succinate) (Mw 2000) or 0.6 mg Bis(succinimidyl penta(ethylene glycol)) (BS(PEG)5, Mw 532.50) (to prepare proteinosomes with lower permeability) are mixed in 15 µL sodium carbonate buffer (50 mM, pH 9.0). The mixture is added to the oil 2-ethyl-1-hexanol (aqueous/oil volume fraction 0.06) and shaken by hand (15 s) to produce a Pickering emulsion. The upper oil layer is removed after 2 h of sedimentation. Then the emulsion is dispersed by addition of 400 µL mixture of ethanol/water (ethanol fraction 70%) and dialyzed sequentially in 70% ethanol (2 h), 50% ethanol (2 h) and water (overnight). The proteinosomes suspension is stored at 4°C.

#### DNA sequence design and synthesis

DNA sequences are designed via MATLAB script that generates random sequences with a G or C fraction between 0.3 and 0.7.

All gate complexes in protocells contain two strands, a gate strand (denoted as **F**) labeled with both a biotin group to enable binding to encapsulated streptavidin and a fluorophore to monitor signal release or receiver activation and an output strand (denoted as **Q**) functionalized with a quencher. Sender protocells used in the spatial Boolean setup (Fig. 4a) consist of gate complexes in which strand **F** is only biotinylated without any other modification while the fluorophore is attached to output strand **Q**.

Sender protocells

The design of DNA gate complex localized inside sender protocells used in the sender-receiver setup (Fig. 1b) and sender-transceiver setup (Fig. 3a) is based on a biotinylated gate strand $\mathbf{F_1}$ labeled with fluorophore Cy5 to monitor the photo-cleavage reaction. Strand $\mathbf{Q_1}$ consist of three sequences ($\mathbf{A_1}$**-PC-**$\mathbf{B_1}$): strand $\mathbf{A_1}$ functions as signal for activation of receiver protocell, strand $\mathbf{B_1}$ is labeled with quencher and a photo-cleavable nitrobenzyl spacer (**PC**) linking strand $\mathbf{A_1}$ and $\mathbf{B_1}$. Before activation $\mathbf{F_1Q_1}$ form a stable duplex where Cy5 fluorescence is quenched. Upon UV irradiation $\mathbf{Q_1}$ is cleaved into two parts which both dissociate from template **F**. Strand $\mathbf{A_1}$ diffuses out from the sender protocells and activate receivers while dissociation of strand $\mathbf{B_1}$ leads to unquenching of the Cy5 fluorescence providing a direct measure for the photo-cleavage reaction. We verified if increase in Cy5 fluorescence directly correlates to signal release by labeling strand $\mathbf{A_1}$ labeled with Cy3 providing a direct measure for signal release (Supplementary Fig. 2).

For the spatial Boolean setup a configuration of two non-identical sender protocells is used, where sender **1** is loaded with gate complex $\mathbf{F_4Q_4}$ and sender **2** is loaded with gate complex $\mathbf{F_5Q_5}$ (Fig. 4a). The corresponding biotinylated gate strands $\mathbf{F_4}$ and $\mathbf{F_5}$ are designed to bind Cy3-modified strands $\mathbf{Q_4}$ ($\mathbf{A_2}$**-PC-**$\mathbf{B_2}$) and $\mathbf{Q_5}$ ($\mathbf{A_3}$**-PC-**$\mathbf{B_3}$) respectively. Upon light activation, two non-identical signal strands $\mathbf{A_2}$ and $\mathbf{A_3}$ diffuse out and activate receiver protocells when the two strands are simultaneously present (AND logic).

The sequences of all gate complexes in sender protocells including their modification are summarized in Supplementary Table 3.

Receiver protocells

The design of DNA gate complex localized inside receiver and transceiver protocells is based on the "seesaw" gate motif[2]. Instead of using separate output and reporter complexes, the gate strands **F** are directly labelled with fluorophore Alexa546 at one end and a biotin group at the other end while the output strand **Q** is functionalized with a quencher. In the initial inactive state, **QF** forms a stable complex where the fluorophore is quenched. When signal strands enter receiver protocells and activate the gate complex by toehold-mediated DSD reaction, the fluorescence is turned on (Fig. 1b, 3a). Additionally, in the sender-transceiver setup a fuel strand is used to recycle the signal strand from the active gate complex, resulting in the regeneration of signal strand which can activate another gate complex (Fig. 3a).

For the spatial Boolean setup, cooperative DSD[3] is used to integrate two non-identical signals in receiver protocells (Fig. 4a). Biotinylated gate strand $\mathbf{F_6}$ is internally modified with a Quasar670 fluorophore whose fluorescence is quenched by binding to quencher-modified output strand $\mathbf{Q_6}$ in the initial inactive state. Upon light activation two non-identical strands $\mathbf{A_2}$ and $\mathbf{A_3}$ released form two distant senders are able to unquench the dye through cooperative DSD.

The sequences including chemical modification of all strands are listed in Supplementary Table 3. All sequences where screened using NUPACK[4] to detect any possible undesired interactions.

All DNA are synthesized by Integrated DNA Technologies (IDT) and BioSearch Technologies. The DNA strands are dissolved in TE buffer (10 mM Tris, 0.1 mM EDTA, pH 8.0, nuclease-free, IDT) as stock solutions (100 µM and 10 µM), which are stored at -20°C for later use.

#### Localization of DNA gate complexes in streptavidin-containing proteinosomes

The localization of DNA gate complexes in proteinosomes is performed in 10 mM Tris buffer with 12 mM $Mg^{2+}$ and 0.1% v/v Tween 20. Typically, a dispersion of streptavidin-containing proteinosomes (10 µL), 4X buffer (5 µL) and biotinylated DNA gate strand F (2 µL from a 10 µM stock) are gently mixed and incubated at room temperature for 1 h, then 3 µL of the corresponding output strand Q (10 µM stock) is added to the mixture which is incubated at 4°C overnight. After incubation, the supernatant (10 µL) is removed carefully from the top and 400 µL buffer is added. The proteinosomes suspension is allowed to sediment for 5 h and 400 µL of supernatant is removed. This process is repeated and the resulting DNA gate-containing proteinosomes are stored at 4°C.

#### Design and fabrication of microfluidic trapping devices

A two-layer microfluidic chip is used to physically trap the proteinosomes. The chip is prepared using a previously reported procedure[5], and contains a localization chamber with PDMS pillars, a filtering chamber, inlet channels with pneumatically actuated push-up valves and outlet channels. Master molds for the bottom and top layers are fabricated on separate silicon wafers (Silicon Materials) using photolithography techniques[6]. The bottom and top layer's mold is made by spin-coating SU8-3050 (50 µm in height) and spin-coating AZ 40xt (40 µm in height) respectively. After development the AZ 40xt mold is reflowed, resulting in rounded channels (60 µm in height) at the center. The two layers are assembled and then plasma bonded to circular #1.5 glass coverslips. Two chips with different density (270 and 210 in 1.5 mm X 2 mm) of traps are fabricated, with the use for density-dependent experiment.

#### Proteinosome trapping and activation

The microfluidic chip is installed on the stage of a confocal laser scanning microscope (Leica SP8). Water-filled control channels are actuated through pneumatic valve array (FESTO) which is connected to a programmable logic controlled (PLC, WAGO Kontakttechnik GmbH). The PLC is then linked to a PC and controlled by a custom Matlab GUI. The pressure in the control channels is 2 bar. The pressure of the inlet channels can be adjusted using pressure regulators (Flow-EZ, Fluigent). Typically, an experiment starts with removing air bubbles and filling all the chambers and channels with buffer solution by pressurizing the buffer channel (buffer solution connects to inlet port 1, Supplementary Fig. 43). Then sender and receiver protocells are loaded sequentially. One DNA gate-localized sender protocell is loaded into the trapping chamber from inlet port 2 (pressure 10 mbar, port 2 is washed with buffer every time before loading proteinosomes). The sender protocell is first trapped at the first row of the chamber, then is pushed through several trapping rows by applying a pulse of flow with high pressure (200 mbar) and eventually locate at the chamber's center approximately. The trapping chamber is washed through flowing in buffer solution (10 mbar) for 5 min to remove any unbound DNA. Next, receiver protocells are loaded into the trapping array from inlet port 2 and a flow of buffer (20 mbar) is applied to remove any unbound DNA and proteinosomes that are not properly trapped, resulting in spatial distribution of protocells contains one sender protocell with surrounding receiver protocells (6-8 shells). In the spatial Boolean setup (Fig. 4a), sender 1 and 2 are loaded sequentially with a distance of 6 to 8 rows, followed by loading of receiver protocells. Solution of fuel strand or exonuclease (if used) with desired concentration is flowing into the trap array (10 mbar) via inlet port 3. Confocal microscope is focused on the trapping chamber, in which the sender protocell is illuminated with light (405 um laser, 100 ms every 1s) and time-lapse imaging (488, 552 and 638 nm lasers, 100 ms every 18s) is started. All the experiments are measured for 2 h and performed at room temperature.

To quantify the permeability of proteinosomes, a solution of Cy3-labeled strand (0.5 μM) is flowed into the chamber loaded with proteinosomes. The diffusion of fluorescent strand is measured over time inside and outside the proteinosomes and the permeability constant is estimated via a previously reported method[5].

All of the experiments in sender-receiver and sender-transceiver systems were performed in independent triplicates.

**Data acquisition and analysis**

Fluorescence data are obtained using confocal microscope with solid state lasers (488 nm for FITC, 552 nm for Cy3 and Alexa546, 638 nm for Cy5 and Quasar670). The objective used is 10X/0.40NA (1.55X1.55mm field of view, 7 μm slice thickness) with a resolution of 512X512 pixels (or alternatively 20X/0.75NA objective with a resolution of 1024X1024 pixels for permeability measurement). A hybrid detector (photon counting mode) is used. Typically, one sender and approximately 150 receiver protocells are recorded for each experimental condition. For spatial Boolean logic setup, two senders are imaged. The relative fluorescence units (RFU) is measured and is used to calculate the concentration of activated DNA gate. For all experiments the baseline valve is subtracted and labeled with "corrected" in the data analysis. The RFU-to-concentration conversion factor is determined by measuring the average RFU (value across a horizontal line through the device) of specific activated DNA gate complex (0.1 μM, 0.5 μM, 1 μM) that is flown into the device. The conversion factor is then determined by plotting the RFU vs the concentration of activated DNA gate complex.

Data processing

All image analysis was performed using MATLAB programming environment and its imaging toolbox (Mathworks). All scripts are available upon request from the corresponding author.

*Protocell detection:* Microscopy images were imported, the FITC channel (label on the membrane of the protocells) was selected and the intensity was adjusted. Noise was removed using a median filter. Subsequently, circular objects (protocells) were detected using a two-stage circular Hough transform using a sensitivity of 0.9. Protocells intersecting with the edge of the image were ignored automatically and obvious misclassifications were discarded manually.

*Protocell fluorescence quantification:* Protocell fluorescence intensities for each fluorophore channel were determined as the average intensity of all pixels for each protocell, ignoring an outer band of pixels to minimize effects of slight discrepancies between the protocells and the Hough transform fitted circles. Fluorescence intensities were converted to fluorophore concentrations using a calibrated reference curve. All data was corrected by background subtraction. Sender and receiver protocells were automatically distinguished based on their fluorescence: Cy5 for the sender and Alexa546 for the receivers in sender-receiver (Fig. 1b) and sender-transceiver (Fig. 3a) systems, Cy3 for the senders and Qusaar670 for receivers in the spatial Boolean setup (Fig. 4a).

*Quantitative analysis of sender-receiver system*

To analyze receiver activation in space and time, receivers were binned into different concentric shells based on their distance toward the sender. The mean value of receiver fluorescence intensity per shell was used to represent time-dependent activation of the corresponding shell (Fig. 1e). Typically, 6~8 shells consisting of around 150 receiver protocells in total were tracked and used for data analysis.

The characteristic length-scale (Fig. 1f) is defined as the average distance that the signaling strands have traveled from the sender at a given time point, which can be described as

$$\text{Eq. 1}\quad \lambda = \frac{\sum L_n \times n_n}{\sum n_n}$$

Where λ is the length scale (μm), $L_n$ (μm) is the distance of individual receiver to the sender and $n_n$ is the amount of activated gate complex (μM) in the corresponding receiver. The amount of activated gate complex ($n_n$) per receiver is determined by the concentration of activated gate complex ($C_n$) and the volume of the corresponding receiver ($V_n$). Then we can write

$$\text{Eq. 2}\quad \lambda = \frac{\sum L_n \times C_n \times V_n}{\sum C_n \times V_n}$$

The volume ($V_n$) is calculated based on the radius ($r_n$) of the protocell, then Eq. 2 can be expressed as:

$$\text{Eq. 3}\quad \lambda = \frac{\sum L_n \times C_n \times 4/3 \times \pi \times r_n^3}{\sum C_n \times 4/3 \times \pi \times r_n^3}$$

$L_n$, $C_n$ and $r_n$ were determined using Matlab® based on the image taken at 2 h. Eq. 3 was used to calculate the length scale. To analyze the activation over time of the protocells with varying distance to the sender (Fig. 1g), the protocells were binned into bins of 50 μm. The fluorescence intensity of each protocell at intervals of exactly 10 minutes was obtained through linear interpolation and the average of each distance bin was plotted for these timepoints.

*Quantitative analysis of sender-transceiver system*

The same procedure used in sender-receiver activation analysis was applied to sender-transceiver systems. Additionally, the 'response time' (Fig. 3d) of each activated transceiver was defined as the timepoint at which the fluorescence reaches 50% of its final value. To remove the effect of protocell activation as a result of background reactions, any protocells with an absolute increase less than 20 RFU were excluded from the analysis.

*Quantitative analysis of Boolean receivers*

To analyze activating dynamics of the spatial Boolean setup where two senders are embedded in a high density of AND receivers, receivers were binned into shells based on their maximum distance to one of the senders (Fig. 4c). The method of quantifying the average activation of protocells per shell remains the same. In addition, response time (Fig. 4d, 4e) of each activated AND gate receiver was defined as the timepoint at which the fluorescence reaches 50% of its final value. To remove background noise a cutoff of minimum increase of 1 RFU was set.

**2D reaction-diffusion simulations**

*Sender-Receiver model*

2D reaction diffusion simulations were performed using a previously developed computational framework, consisting of Visual DSD, Chemical Reaction Network (CRN), Command Line Interface (CLI) and custom MATLAB scripts for the generation of configuration files and data analysis[5,7,8]. Protocells were placed on a 2D grid (1.5 mm X 1.5 mm) with the same protocell density as the experimental setup (90 protocells per $mm^2$). One single sender protocell was assigned to be located at the center of the area and surrounded by 212 receiver protocells. The concentrations of gate complex in protocells were obtained from a 2D slice. Next, the model parameters including signal release rate from the sender, consumption capacity of receivers, size and permeability of proteinosomes were derived from experimental measurements. The diffusion coefficient of the DNA strands in aqueous buffer was estimated from an earlier study[9]. The rate constant of signal release ($k_{rel}$) and DSD reactions in protocells ($k_{dsd}$) were determined experimentally. The reactions in Visual DSD code are listed below

1. SenderGate -> {$k_{rel}$} signal
2. SenderCel + Signal <-> {(3/rprot)*P}{(3/rprot)*P} SignalSol + ReceiverCel
3. ReceiverCel + SignalSol <-> {(3/rprot)*P}{(3/rprot)*P} Signal + ReceiverCel
4. Signal + ReceiverGate -> {$k_{dsd}$} Reporter + W1

Reaction 1 describes the release of signal from the sender protocell. Reaction 2 and 3 represent the processes of signal diffuses out of the sender and enters the receiver respectively (Cel: inside the protocell; Sol: outside the protocell; P: permeability; rprot: radius of protocell). Reaction 4 is the DSD reaction of signal with gate complex resulting in receiver activation (reporter: activated gate complex; W1: leaving strand as waste). The Matlab function “std” was use to calculate the standard deviation of receiver activation per shell, which originated from different distance of individual receiver protocells to sender protocell. The corresponding parameters are displayed in Supplementary Table 4. The complete description of the mathematical model and computer code is given in Supplementary Note 1.

*Sender-Transceiver model*

To simulate the sender-transceiver system, the sender-receiver system was supplemented with fuel strands capable of regenerating signal by DSD reactions ($k_{fuel}$). The $k_{fuel}$ was assumed to be the same as $k_{dsd}$. The fuel strands were assumed to have two distinct species: a ‘good’ fuel that reacts with activated transceiver gate and release signal as designed, and a ‘bad’ fuel that binds to activated transceiver gate but cannot release the signal strand[10]. Additionally, a leakage reaction was introduced to describe the activation caused by fuel strands in the absence of signal input ($k_{leak}$). The fraction of bad fuel and $k_{leak}$ were optimized manually to match the experimental data. The reactions in Visual DSD code are listed below

1. SenderGate -> {$k_{rel}$} Signal
2. SenderCel + Signal <-> {(3/rprot)*P}{(3/rprot)*P} SignalSol + ReceiverCel
3. TransceiverCel + SignalSol <-> {(3/rprot)*P}{(3/rprot)*P} Signal + TransceiverCel
4. TransceiverCel + FuelGSol <-> {(3/rprot)*P}{(3/rprot)*P} FuelG + TransceiverCel
5. TransceiverCel + FuelBSol <-> {(3/rprot)*P}{(3/rprot)*P} FuelB + TransceiverCel
6. Signal + TransceiverGate -> {$k_{dsd}$} Reporter + W1
7. Reporter + FuelG -> {$k_{fuel}$} Reporter + Signal
8. Reporter + FuelB -> {$k_{fuel}$} Reporter + Signal

9. FuelG + TransceiverGate -> {$k_{leak}$} Reporter

10. FuelB + TransceiverGate -> {$k_{leak}$} Reporter

Reaction 1 describes the release of signal from the sender protocell. Reactions 2 and 3 represent the processes of signal diffuses out if the sender and enters the receiver respectively (Cel: inside the protocell; Sol: outside the protocell; P: permeability; rprot: radius of protocell). Reactions 4 and 5 represent the processes of good and bad fuel strands[10] entering the transceiver respectively (FuelG: good fuel; FuelB: bad fuel). Reaction 6 is the DSD reaction of signal with gate complex resulting in transceiver activation (reporter: activated gate complex; W1: leaving strand as waste). Reactions 7 and 8 are the DSD reactions of fuel with activated gate complex where only good fuel leads to signal regeneration. Reactions 9 and 10 are the leak reactions caused by the fuel strands. The Matlab function "std" was use to calculate the standard deviation of transceiver activation per shell, which originated from different distance of individual transceiver protocells to sender protocell. The corresponding parameters are listed in Supplementary Table 5.

*Spatial Boolean AND gate model*

In the Boolean AND gate model, two distinct senders (sender **1** and sender **2**) were embedded into a population of receiver protocells. The three populations were assigned the same spatial distribution as the experimental setup. Sender **1** and sender **2** were assumed to have the same signal release rate ($k_{ref}$). Two different signals **$A_2$** (release from sender **1**) and **$A_3$** (release from sender **2**) individually bind to the gate complexes (**$F_6Q_6$**) in receiver protocells in a reversible manner and they have the same forward ($k_f$) and backward ($k_b$) rate constants. The forward rate constant was set to be the same as the $k_{dsd}$ used in sender-receiver system because the gate complexes in receiver protocells have comparable lengths of toehold. The backward rate constant was estimated using forward rate constant and the equilibrium constant of the complex derived from NUPACK. The formed intermediate species **$A_2$-$F_6Q_6$** (intermediate 1) or **$A_3$-$F_6Q_6$** (intermediate 2) can irreversibly react with **$A_3$** or **$A_2$** respectively resulting in the activation of AND gate receiver protocells. The irreversible reaction shares the same rate constant $k_f$ from the reversible reaction[3]. The reactions in Visual DSD format are listed below

1. SenderGate1 -> {$k_{rel}$} signal$A_2$
2. SenderCel1 + Signal$A_2$ <-> {(3/rprot)*P}{(3/rprot)*P} Signal$A_2$Sol + ReceiverCel
3. ReceiverCel + Signal$A_2$Sol <-> {(3/rprot)*P}{(3/rprot)*P} Signal$A_2$ + ReceiverCel
4. Signal$A_2$ + ReceiverGate <-> {$k_f$}{$k_b$} Intermediate1
5. Signal$A_3$ + Intermediate1 -> {$k_f$} Reporter + W1
6. SenderGate2 -> {$k_{rel}$} Signal$A_3$
7. SenderCel2 + Signal$A_3$ <-> {(3/rprot)*P}{(3/rprot)*P} Signal$A_3$Sol + ReceiverCel
8. ReceiverCel + Signal$A_3$Sol <-> {(3/rprot)*P}{(3/rprot)*P} Signal$A_3$ + ReceiverCel
9. Signal$A_3$ + ReceiverGate <-> {$k_f$}{$k_b$} Intermediate2
10. Signal$A_2$ + Intermediate2 -> {$k_f$} Reporter + W1

Reaction 1 describes the release of signal **$A_2$** from the sender **1**. Reactions 2 and 3 represent the processes of signal **$A_2$** diffuses out of the sender and enters the receiver respectively (Cel: inside the protocell; Sol: outside the protocell; P: permeability; rprot: radius of protocell). Reaction 4 is the reversible DSD reaction of signal with gate complex resulting in the formation of a triple-stranded intermediate **1** ($k_f$: forward reaction;

$k_b$: backward reaction). Reaction 5 is the DSD reaction of another signal $\mathbf{A_3}$ with intermediate **1** leading to activation of AND-gate receiver (reporter: activated gate complex; W1: leaving strand as waste). Similar to sender **1**, the activation pathway initiated by sender **2** is implemented by reactions 6-10. The Matlab function "std" was use to calculate the standard deviation of receiver activation per shell, which originated from different distance of individual receiver protocells to sender protocell. The corresponding parameters are displayed in Supplementary Table 6.

**Supplementary Note 1. Visual DSD code of the 2D simulation of the sender-receiver system.**

```
directive simulator pde
directive simulation {
  final= 120.000000;
  points= 120;
  plots=[ F1Act ; F2Q1 ;]
}

directive spatial {
  dimensions = 2;
  dt = 0.020000;
  xmax = 1500;
  nx = 201;
  boundary = ZeroFlux;
  diffusibles = [Q1sol = 2227.400000 ; Q2sol = 2227.400000];  //um^2 min^-1
}

directive parameters [
  rprot = 20;
  kdis = 0.002000;
  kdis2 = 0.027000;
  P = 200.000000;
]

| init psA {spatial = { points = [
  { width = 0.026667; value = 1; x = 0.500000; y = 0.500000 };
  ] } }

| init F1Q1 {spatial = { points = [
  { width = 0.026667; value = 26000.000000; x = 0.500000; y = 0.500000 };
  ] } }
```

```
| init psB {spatial = { points = [
  { width = 0.026667; value = 1; x = 0.020000; y = 0.020000 };
  { width = 0.026667; value = 1; x = 0.100000; y = 0.020000 };
  { width = 0.026667; value = 1; x = 0.180000; y = 0.020000 };
  { width = 0.026667; value = 1; x = 0.260000; y = 0.020000 };
  { width = 0.026667; value = 1; x = 0.340000; y = 0.020000 };
  { width = 0.026667; value = 1; x = 0.420000; y = 0.020000 };
  { width = 0.026667; value = 1; x = 0.500000; y = 0.020000 };
  { width = 0.026667; value = 1; x = 0.580000; y = 0.020000 };
  { width = 0.026667; value = 1; x = 0.660000; y = 0.020000 };
  { width = 0.026667; value = 1; x = 0.740000; y = 0.020000 };
  { width = 0.026667; value = 1; x = 0.820000; y = 0.020000 };
  { width = 0.026667; value = 1; x = 0.900000; y = 0.020000 };
  { width = 0.026667; value = 1; x = 0.980000; y = 0.020000 };
  { width = 0.026667; value = 1; x = 0.060000; y = 0.080000 };
  { width = 0.026667; value = 1; x = 0.140000; y = 0.080000 };
  { width = 0.026667; value = 1; x = 0.220000; y = 0.080000 };
  { width = 0.026667; value = 1; x = 0.300000; y = 0.080000 };
  { width = 0.026667; value = 1; x = 0.380000; y = 0.080000 };
  { width = 0.026667; value = 1; x = 0.460000; y = 0.080000 };
  { width = 0.026667; value = 1; x = 0.540000; y = 0.080000 };
  { width = 0.026667; value = 1; x = 0.620000; y = 0.080000 };
  { width = 0.026667; value = 1; x = 0.700000; y = 0.080000 };
  { width = 0.026667; value = 1; x = 0.780000; y = 0.080000 };
  { width = 0.026667; value = 1; x = 0.860000; y = 0.080000 };
  { width = 0.026667; value = 1; x = 0.940000; y = 0.080000 };
  { width = 0.026667; value = 1; x = 0.020000; y = 0.140000 };
  { width = 0.026667; value = 1; x = 0.100000; y = 0.140000 };
  { width = 0.026667; value = 1; x = 0.180000; y = 0.140000 };
  { width = 0.026667; value = 1; x = 0.260000; y = 0.140000 };
  { width = 0.026667; value = 1; x = 0.340000; y = 0.140000 };
```

{ width = 0.026667; value = 1; x = 0.420000; y = 0.140000 };

{ width = 0.026667; value = 1; x = 0.500000; y = 0.140000 };

{ width = 0.026667; value = 1; x = 0.580000; y = 0.140000 };

{ width = 0.026667; value = 1; x = 0.660000; y = 0.140000 };

{ width = 0.026667; value = 1; x = 0.740000; y = 0.140000 };

{ width = 0.026667; value = 1; x = 0.820000; y = 0.140000 };

{ width = 0.026667; value = 1; x = 0.900000; y = 0.140000 };

{ width = 0.026667; value = 1; x = 0.980000; y = 0.140000 };

{ width = 0.026667; value = 1; x = 0.060000; y = 0.200000 };

{ width = 0.026667; value = 1; x = 0.140000; y = 0.200000 };

{ width = 0.026667; value = 1; x = 0.220000; y = 0.200000 };

{ width = 0.026667; value = 1; x = 0.300000; y = 0.200000 };

{ width = 0.026667; value = 1; x = 0.380000; y = 0.200000 };

{ width = 0.026667; value = 1; x = 0.460000; y = 0.200000 };

{ width = 0.026667; value = 1; x = 0.540000; y = 0.200000 };

{ width = 0.026667; value = 1; x = 0.620000; y = 0.200000 };

{ width = 0.026667; value = 1; x = 0.700000; y = 0.200000 };

{ width = 0.026667; value = 1; x = 0.780000; y = 0.200000 };

{ width = 0.026667; value = 1; x = 0.860000; y = 0.200000 };

{ width = 0.026667; value = 1; x = 0.940000; y = 0.200000 };

{ width = 0.026667; value = 1; x = 0.020000; y = 0.260000 };

{ width = 0.026667; value = 1; x = 0.100000; y = 0.260000 };

{ width = 0.026667; value = 1; x = 0.180000; y = 0.260000 };

{ width = 0.026667; value = 1; x = 0.260000; y = 0.260000 };

{ width = 0.026667; value = 1; x = 0.340000; y = 0.260000 };

{ width = 0.026667; value = 1; x = 0.420000; y = 0.260000 };

{ width = 0.026667; value = 1; x = 0.500000; y = 0.260000 };

{ width = 0.026667; value = 1; x = 0.580000; y = 0.260000 };

{ width = 0.026667; value = 1; x = 0.660000; y = 0.260000 };

{ width = 0.026667; value = 1; x = 0.740000; y = 0.260000 };

{ width = 0.026667; value = 1; x = 0.820000; y = 0.260000 };

{ width = 0.026667; value = 1; x = 0.900000; y = 0.260000 };

{ width = 0.026667; value = 1; x = 0.980000; y = 0.260000 };

{ width = 0.026667; value = 1; x = 0.060000; y = 0.320000 };

{ width = 0.026667; value = 1; x = 0.140000; y = 0.320000 };

{ width = 0.026667; value = 1; x = 0.220000; y = 0.320000 };

{ width = 0.026667; value = 1; x = 0.300000; y = 0.320000 };

{ width = 0.026667; value = 1; x = 0.380000; y = 0.320000 };

{ width = 0.026667; value = 1; x = 0.460000; y = 0.320000 };

{ width = 0.026667; value = 1; x = 0.540000; y = 0.320000 };

{ width = 0.026667; value = 1; x = 0.620000; y = 0.320000 };

{ width = 0.026667; value = 1; x = 0.700000; y = 0.320000 };

{ width = 0.026667; value = 1; x = 0.780000; y = 0.320000 };

{ width = 0.026667; value = 1; x = 0.860000; y = 0.320000 };

{ width = 0.026667; value = 1; x = 0.940000; y = 0.320000 };

{ width = 0.026667; value = 1; x = 0.020000; y = 0.380000 };

{ width = 0.026667; value = 1; x = 0.100000; y = 0.380000 };

{ width = 0.026667; value = 1; x = 0.180000; y = 0.380000 };

{ width = 0.026667; value = 1; x = 0.260000; y = 0.380000 };

{ width = 0.026667; value = 1; x = 0.340000; y = 0.380000 };

{ width = 0.026667; value = 1; x = 0.420000; y = 0.380000 };

{ width = 0.026667; value = 1; x = 0.500000; y = 0.380000 };

{ width = 0.026667; value = 1; x = 0.580000; y = 0.380000 };

{ width = 0.026667; value = 1; x = 0.660000; y = 0.380000 };

{ width = 0.026667; value = 1; x = 0.740000; y = 0.380000 };

{ width = 0.026667; value = 1; x = 0.820000; y = 0.380000 };

{ width = 0.026667; value = 1; x = 0.900000; y = 0.380000 };

{ width = 0.026667; value = 1; x = 0.980000; y = 0.380000 };

{ width = 0.026667; value = 1; x = 0.060000; y = 0.440000 };

{ width = 0.026667; value = 1; x = 0.140000; y = 0.440000 };

{ width = 0.026667; value = 1; x = 0.220000; y = 0.440000 };

{ width = 0.026667; value = 1; x = 0.300000; y = 0.440000 };

{ width = 0.026667; value = 1; x = 0.380000; y = 0.440000 };

{ width = 0.026667; value = 1; x = 0.460000; y = 0.440000 };

{ width = 0.026667; value = 1; x = 0.540000; y = 0.440000 };
{ width = 0.026667; value = 1; x = 0.620000; y = 0.440000 };
{ width = 0.026667; value = 1; x = 0.700000; y = 0.440000 };
{ width = 0.026667; value = 1; x = 0.780000; y = 0.440000 };
{ width = 0.026667; value = 1; x = 0.860000; y = 0.440000 };
{ width = 0.026667; value = 1; x = 0.940000; y = 0.440000 };
{ width = 0.026667; value = 1; x = 0.020000; y = 0.500000 };
{ width = 0.026667; value = 1; x = 0.100000; y = 0.500000 };
{ width = 0.026667; value = 1; x = 0.180000; y = 0.500000 };
{ width = 0.026667; value = 1; x = 0.260000; y = 0.500000 };
{ width = 0.026667; value = 1; x = 0.340000; y = 0.500000 };
{ width = 0.026667; value = 1; x = 0.420000; y = 0.500000 };
{ width = 0.026667; value = 1; x = 0.580000; y = 0.500000 };
{ width = 0.026667; value = 1; x = 0.660000; y = 0.500000 };
{ width = 0.026667; value = 1; x = 0.740000; y = 0.500000 };
{ width = 0.026667; value = 1; x = 0.820000; y = 0.500000 };
{ width = 0.026667; value = 1; x = 0.900000; y = 0.500000 };
{ width = 0.026667; value = 1; x = 0.980000; y = 0.500000 };
{ width = 0.026667; value = 1; x = 0.060000; y = 0.560000 };
{ width = 0.026667; value = 1; x = 0.140000; y = 0.560000 };
{ width = 0.026667; value = 1; x = 0.220000; y = 0.560000 };
{ width = 0.026667; value = 1; x = 0.300000; y = 0.560000 };
{ width = 0.026667; value = 1; x = 0.380000; y = 0.560000 };
{ width = 0.026667; value = 1; x = 0.460000; y = 0.560000 };
{ width = 0.026667; value = 1; x = 0.540000; y = 0.560000 };
{ width = 0.026667; value = 1; x = 0.620000; y = 0.560000 };
{ width = 0.026667; value = 1; x = 0.700000; y = 0.560000 };
{ width = 0.026667; value = 1; x = 0.780000; y = 0.560000 };
{ width = 0.026667; value = 1; x = 0.860000; y = 0.560000 };
{ width = 0.026667; value = 1; x = 0.940000; y = 0.560000 };
{ width = 0.026667; value = 1; x = 0.020000; y = 0.620000 };
{ width = 0.026667; value = 1; x = 0.100000; y = 0.620000 };

{ width = 0.026667; value = 1; x = 0.180000; y = 0.620000 };
{ width = 0.026667; value = 1; x = 0.260000; y = 0.620000 };
{ width = 0.026667; value = 1; x = 0.340000; y = 0.620000 };
{ width = 0.026667; value = 1; x = 0.420000; y = 0.620000 };
{ width = 0.026667; value = 1; x = 0.500000; y = 0.620000 };
{ width = 0.026667; value = 1; x = 0.580000; y = 0.620000 };
{ width = 0.026667; value = 1; x = 0.660000; y = 0.620000 };
{ width = 0.026667; value = 1; x = 0.740000; y = 0.620000 };
{ width = 0.026667; value = 1; x = 0.820000; y = 0.620000 };
{ width = 0.026667; value = 1; x = 0.900000; y = 0.620000 };
{ width = 0.026667; value = 1; x = 0.980000; y = 0.620000 };
{ width = 0.026667; value = 1; x = 0.060000; y = 0.680000 };
{ width = 0.026667; value = 1; x = 0.140000; y = 0.680000 };
{ width = 0.026667; value = 1; x = 0.220000; y = 0.680000 };
{ width = 0.026667; value = 1; x = 0.300000; y = 0.680000 };
{ width = 0.026667; value = 1; x = 0.380000; y = 0.680000 };
{ width = 0.026667; value = 1; x = 0.460000; y = 0.680000 };
{ width = 0.026667; value = 1; x = 0.540000; y = 0.680000 };
{ width = 0.026667; value = 1; x = 0.620000; y = 0.680000 };
{ width = 0.026667; value = 1; x = 0.700000; y = 0.680000 };
{ width = 0.026667; value = 1; x = 0.780000; y = 0.680000 };
{ width = 0.026667; value = 1; x = 0.860000; y = 0.680000 };
{ width = 0.026667; value = 1; x = 0.940000; y = 0.680000 };
{ width = 0.026667; value = 1; x = 0.020000; y = 0.740000 };
{ width = 0.026667; value = 1; x = 0.100000; y = 0.740000 };
{ width = 0.026667; value = 1; x = 0.180000; y = 0.740000 };
{ width = 0.026667; value = 1; x = 0.260000; y = 0.740000 };
{ width = 0.026667; value = 1; x = 0.340000; y = 0.740000 };
{ width = 0.026667; value = 1; x = 0.420000; y = 0.740000 };
{ width = 0.026667; value = 1; x = 0.500000; y = 0.740000 };
{ width = 0.026667; value = 1; x = 0.580000; y = 0.740000 };
{ width = 0.026667; value = 1; x = 0.660000; y = 0.740000 };

{ width = 0.026667; value = 1; x = 0.740000; y = 0.740000 };

{ width = 0.026667; value = 1; x = 0.820000; y = 0.740000 };

{ width = 0.026667; value = 1; x = 0.900000; y = 0.740000 };

{ width = 0.026667; value = 1; x = 0.980000; y = 0.740000 };

{ width = 0.026667; value = 1; x = 0.060000; y = 0.800000 };

{ width = 0.026667; value = 1; x = 0.140000; y = 0.800000 };

{ width = 0.026667; value = 1; x = 0.220000; y = 0.800000 };

{ width = 0.026667; value = 1; x = 0.300000; y = 0.800000 };

{ width = 0.026667; value = 1; x = 0.380000; y = 0.800000 };

{ width = 0.026667; value = 1; x = 0.460000; y = 0.800000 };

{ width = 0.026667; value = 1; x = 0.540000; y = 0.800000 };

{ width = 0.026667; value = 1; x = 0.620000; y = 0.800000 };

{ width = 0.026667; value = 1; x = 0.700000; y = 0.800000 };

{ width = 0.026667; value = 1; x = 0.780000; y = 0.800000 };

{ width = 0.026667; value = 1; x = 0.860000; y = 0.800000 };

{ width = 0.026667; value = 1; x = 0.940000; y = 0.800000 };

{ width = 0.026667; value = 1; x = 0.020000; y = 0.860000 };

{ width = 0.026667; value = 1; x = 0.100000; y = 0.860000 };

{ width = 0.026667; value = 1; x = 0.180000; y = 0.860000 };

{ width = 0.026667; value = 1; x = 0.260000; y = 0.860000 };

{ width = 0.026667; value = 1; x = 0.340000; y = 0.860000 };

{ width = 0.026667; value = 1; x = 0.420000; y = 0.860000 };

{ width = 0.026667; value = 1; x = 0.500000; y = 0.860000 };

{ width = 0.026667; value = 1; x = 0.580000; y = 0.860000 };

{ width = 0.026667; value = 1; x = 0.660000; y = 0.860000 };

{ width = 0.026667; value = 1; x = 0.740000; y = 0.860000 };

{ width = 0.026667; value = 1; x = 0.820000; y = 0.860000 };

{ width = 0.026667; value = 1; x = 0.900000; y = 0.860000 };

{ width = 0.026667; value = 1; x = 0.980000; y = 0.860000 };

{ width = 0.026667; value = 1; x = 0.060000; y = 0.920000 };

{ width = 0.026667; value = 1; x = 0.140000; y = 0.920000 };

{ width = 0.026667; value = 1; x = 0.220000; y = 0.920000 };

```
 { width = 0.026667; value = 1; x = 0.300000; y = 0.920000 };
 { width = 0.026667; value = 1; x = 0.380000; y = 0.920000 };
 { width = 0.026667; value = 1; x = 0.460000; y = 0.920000 };
 { width = 0.026667; value = 1; x = 0.540000; y = 0.920000 };
 { width = 0.026667; value = 1; x = 0.620000; y = 0.920000 };
 { width = 0.026667; value = 1; x = 0.700000; y = 0.920000 };
 { width = 0.026667; value = 1; x = 0.780000; y = 0.920000 };
 { width = 0.026667; value = 1; x = 0.860000; y = 0.920000 };
 { width = 0.026667; value = 1; x = 0.940000; y = 0.920000 };
 { width = 0.026667; value = 1; x = 0.020000; y = 0.980000 };
 { width = 0.026667; value = 1; x = 0.100000; y = 0.980000 };
 { width = 0.026667; value = 1; x = 0.180000; y = 0.980000 };
 { width = 0.026667; value = 1; x = 0.260000; y = 0.980000 };
 { width = 0.026667; value = 1; x = 0.340000; y = 0.980000 };
 { width = 0.026667; value = 1; x = 0.420000; y = 0.980000 };
 { width = 0.026667; value = 1; x = 0.500000; y = 0.980000 };
 { width = 0.026667; value = 1; x = 0.580000; y = 0.980000 };
 { width = 0.026667; value = 1; x = 0.660000; y = 0.980000 };
 { width = 0.026667; value = 1; x = 0.740000; y = 0.980000 };
 { width = 0.026667; value = 1; x = 0.820000; y = 0.980000 };
 { width = 0.026667; value = 1; x = 0.900000; y = 0.980000 };
 { width = 0.026667; value = 1; x = 0.980000; y = 0.980000 };
 ] } }

| init F2Q2 {spatial = { points = [
 { width = 0.026667; value = 2000.000000; x = 0.020000; y = 0.020000 };
 { width = 0.026667; value = 2000.000000; x = 0.100000; y = 0.020000 };
 { width = 0.026667; value = 2000.000000; x = 0.180000; y = 0.020000 };
 { width = 0.026667; value = 2000.000000; x = 0.260000; y = 0.020000 };
 { width = 0.026667; value = 2000.000000; x = 0.340000; y = 0.020000 };
 { width = 0.026667; value = 2000.000000; x = 0.420000; y = 0.020000 };
 { width = 0.026667; value = 2000.000000; x = 0.500000; y = 0.020000 };
```

{ width = 0.026667; value = 2000.000000; x = 0.580000; y = 0.020000 };
{ width = 0.026667; value = 2000.000000; x = 0.660000; y = 0.020000 };
{ width = 0.026667; value = 2000.000000; x = 0.740000; y = 0.020000 };
{ width = 0.026667; value = 2000.000000; x = 0.820000; y = 0.020000 };
{ width = 0.026667; value = 2000.000000; x = 0.900000; y = 0.020000 };
{ width = 0.026667; value = 2000.000000; x = 0.980000; y = 0.020000 };
{ width = 0.026667; value = 2000.000000; x = 0.060000; y = 0.080000 };
{ width = 0.026667; value = 2000.000000; x = 0.140000; y = 0.080000 };
{ width = 0.026667; value = 2000.000000; x = 0.220000; y = 0.080000 };
{ width = 0.026667; value = 2000.000000; x = 0.300000; y = 0.080000 };
{ width = 0.026667; value = 2000.000000; x = 0.380000; y = 0.080000 };
{ width = 0.026667; value = 2000.000000; x = 0.460000; y = 0.080000 };
{ width = 0.026667; value = 2000.000000; x = 0.540000; y = 0.080000 };
{ width = 0.026667; value = 2000.000000; x = 0.620000; y = 0.080000 };
{ width = 0.026667; value = 2000.000000; x = 0.700000; y = 0.080000 };
{ width = 0.026667; value = 2000.000000; x = 0.780000; y = 0.080000 };
{ width = 0.026667; value = 2000.000000; x = 0.860000; y = 0.080000 };
{ width = 0.026667; value = 2000.000000; x = 0.940000; y = 0.080000 };
{ width = 0.026667; value = 2000.000000; x = 0.020000; y = 0.140000 };
{ width = 0.026667; value = 2000.000000; x = 0.100000; y = 0.140000 };
{ width = 0.026667; value = 2000.000000; x = 0.180000; y = 0.140000 };
{ width = 0.026667; value = 2000.000000; x = 0.260000; y = 0.140000 };
{ width = 0.026667; value = 2000.000000; x = 0.340000; y = 0.140000 };
{ width = 0.026667; value = 2000.000000; x = 0.420000; y = 0.140000 };
{ width = 0.026667; value = 2000.000000; x = 0.500000; y = 0.140000 };
{ width = 0.026667; value = 2000.000000; x = 0.580000; y = 0.140000 };
{ width = 0.026667; value = 2000.000000; x = 0.660000; y = 0.140000 };
{ width = 0.026667; value = 2000.000000; x = 0.740000; y = 0.140000 };
{ width = 0.026667; value = 2000.000000; x = 0.820000; y = 0.140000 };
{ width = 0.026667; value = 2000.000000; x = 0.900000; y = 0.140000 };
{ width = 0.026667; value = 2000.000000; x = 0.980000; y = 0.140000 };
{ width = 0.026667; value = 2000.000000; x = 0.060000; y = 0.200000 };

{ width = 0.026667; value = 2000.000000; x = 0.140000; y = 0.200000 };

{ width = 0.026667; value = 2000.000000; x = 0.220000; y = 0.200000 };

{ width = 0.026667; value = 2000.000000; x = 0.300000; y = 0.200000 };

{ width = 0.026667; value = 2000.000000; x = 0.380000; y = 0.200000 };

{ width = 0.026667; value = 2000.000000; x = 0.460000; y = 0.200000 };

{ width = 0.026667; value = 2000.000000; x = 0.540000; y = 0.200000 };

{ width = 0.026667; value = 2000.000000; x = 0.620000; y = 0.200000 };

{ width = 0.026667; value = 2000.000000; x = 0.700000; y = 0.200000 };

{ width = 0.026667; value = 2000.000000; x = 0.780000; y = 0.200000 };

{ width = 0.026667; value = 2000.000000; x = 0.860000; y = 0.200000 };

{ width = 0.026667; value = 2000.000000; x = 0.940000; y = 0.200000 };

{ width = 0.026667; value = 2000.000000; x = 0.020000; y = 0.260000 };

{ width = 0.026667; value = 2000.000000; x = 0.100000; y = 0.260000 };

{ width = 0.026667; value = 2000.000000; x = 0.180000; y = 0.260000 };

{ width = 0.026667; value = 2000.000000; x = 0.260000; y = 0.260000 };

{ width = 0.026667; value = 2000.000000; x = 0.340000; y = 0.260000 };

{ width = 0.026667; value = 2000.000000; x = 0.420000; y = 0.260000 };

{ width = 0.026667; value = 2000.000000; x = 0.500000; y = 0.260000 };

{ width = 0.026667; value = 2000.000000; x = 0.580000; y = 0.260000 };

{ width = 0.026667; value = 2000.000000; x = 0.660000; y = 0.260000 };

{ width = 0.026667; value = 2000.000000; x = 0.740000; y = 0.260000 };

{ width = 0.026667; value = 2000.000000; x = 0.820000; y = 0.260000 };

{ width = 0.026667; value = 2000.000000; x = 0.900000; y = 0.260000 };

{ width = 0.026667; value = 2000.000000; x = 0.980000; y = 0.260000 };

{ width = 0.026667; value = 2000.000000; x = 0.060000; y = 0.320000 };

{ width = 0.026667; value = 2000.000000; x = 0.140000; y = 0.320000 };

{ width = 0.026667; value = 2000.000000; x = 0.220000; y = 0.320000 };

{ width = 0.026667; value = 2000.000000; x = 0.300000; y = 0.320000 };

{ width = 0.026667; value = 2000.000000; x = 0.380000; y = 0.320000 };

{ width = 0.026667; value = 2000.000000; x = 0.460000; y = 0.320000 };

{ width = 0.026667; value = 2000.000000; x = 0.540000; y = 0.320000 };

{ width = 0.026667; value = 2000.000000; x = 0.620000; y = 0.320000 };

{ width = 0.026667; value = 2000.000000; x = 0.700000; y = 0.320000 };

{ width = 0.026667; value = 2000.000000; x = 0.780000; y = 0.320000 };

{ width = 0.026667; value = 2000.000000; x = 0.860000; y = 0.320000 };

{ width = 0.026667; value = 2000.000000; x = 0.940000; y = 0.320000 };

{ width = 0.026667; value = 2000.000000; x = 0.020000; y = 0.380000 };

{ width = 0.026667; value = 2000.000000; x = 0.100000; y = 0.380000 };

{ width = 0.026667; value = 2000.000000; x = 0.180000; y = 0.380000 };

{ width = 0.026667; value = 2000.000000; x = 0.260000; y = 0.380000 };

{ width = 0.026667; value = 2000.000000; x = 0.340000; y = 0.380000 };

{ width = 0.026667; value = 2000.000000; x = 0.420000; y = 0.380000 };

{ width = 0.026667; value = 2000.000000; x = 0.500000; y = 0.380000 };

{ width = 0.026667; value = 2000.000000; x = 0.580000; y = 0.380000 };

{ width = 0.026667; value = 2000.000000; x = 0.660000; y = 0.380000 };

{ width = 0.026667; value = 2000.000000; x = 0.740000; y = 0.380000 };

{ width = 0.026667; value = 2000.000000; x = 0.820000; y = 0.380000 };

{ width = 0.026667; value = 2000.000000; x = 0.900000; y = 0.380000 };

{ width = 0.026667; value = 2000.000000; x = 0.980000; y = 0.380000 };

{ width = 0.026667; value = 2000.000000; x = 0.060000; y = 0.440000 };

{ width = 0.026667; value = 2000.000000; x = 0.140000; y = 0.440000 };

{ width = 0.026667; value = 2000.000000; x = 0.220000; y = 0.440000 };

{ width = 0.026667; value = 2000.000000; x = 0.300000; y = 0.440000 };

{ width = 0.026667; value = 2000.000000; x = 0.380000; y = 0.440000 };

{ width = 0.026667; value = 2000.000000; x = 0.460000; y = 0.440000 };

{ width = 0.026667; value = 2000.000000; x = 0.540000; y = 0.440000 };

{ width = 0.026667; value = 2000.000000; x = 0.620000; y = 0.440000 };

{ width = 0.026667; value = 2000.000000; x = 0.700000; y = 0.440000 };

{ width = 0.026667; value = 2000.000000; x = 0.780000; y = 0.440000 };

{ width = 0.026667; value = 2000.000000; x = 0.860000; y = 0.440000 };

{ width = 0.026667; value = 2000.000000; x = 0.940000; y = 0.440000 };

{ width = 0.026667; value = 2000.000000; x = 0.020000; y = 0.500000 };

{ width = 0.026667; value = 2000.000000; x = 0.100000; y = 0.500000 };

{ width = 0.026667; value = 2000.000000; x = 0.180000; y = 0.500000 };

{ width = 0.026667; value = 2000.000000; x = 0.260000; y = 0.500000 };

{ width = 0.026667; value = 2000.000000; x = 0.340000; y = 0.500000 };

{ width = 0.026667; value = 2000.000000; x = 0.420000; y = 0.500000 };

{ width = 0.026667; value = 2000.000000; x = 0.580000; y = 0.500000 };

{ width = 0.026667; value = 2000.000000; x = 0.660000; y = 0.500000 };

{ width = 0.026667; value = 2000.000000; x = 0.740000; y = 0.500000 };

{ width = 0.026667; value = 2000.000000; x = 0.820000; y = 0.500000 };

{ width = 0.026667; value = 2000.000000; x = 0.900000; y = 0.500000 };

{ width = 0.026667; value = 2000.000000; x = 0.980000; y = 0.500000 };

{ width = 0.026667; value = 2000.000000; x = 0.060000; y = 0.560000 };

{ width = 0.026667; value = 2000.000000; x = 0.140000; y = 0.560000 };

{ width = 0.026667; value = 2000.000000; x = 0.220000; y = 0.560000 };

{ width = 0.026667; value = 2000.000000; x = 0.300000; y = 0.560000 };

{ width = 0.026667; value = 2000.000000; x = 0.380000; y = 0.560000 };

{ width = 0.026667; value = 2000.000000; x = 0.460000; y = 0.560000 };

{ width = 0.026667; value = 2000.000000; x = 0.540000; y = 0.560000 };

{ width = 0.026667; value = 2000.000000; x = 0.620000; y = 0.560000 };

{ width = 0.026667; value = 2000.000000; x = 0.700000; y = 0.560000 };

{ width = 0.026667; value = 2000.000000; x = 0.780000; y = 0.560000 };

{ width = 0.026667; value = 2000.000000; x = 0.860000; y = 0.560000 };

{ width = 0.026667; value = 2000.000000; x = 0.940000; y = 0.560000 };

{ width = 0.026667; value = 2000.000000; x = 0.020000; y = 0.620000 };

{ width = 0.026667; value = 2000.000000; x = 0.100000; y = 0.620000 };

{ width = 0.026667; value = 2000.000000; x = 0.180000; y = 0.620000 };

{ width = 0.026667; value = 2000.000000; x = 0.260000; y = 0.620000 };

{ width = 0.026667; value = 2000.000000; x = 0.340000; y = 0.620000 };

{ width = 0.026667; value = 2000.000000; x = 0.420000; y = 0.620000 };

{ width = 0.026667; value = 2000.000000; x = 0.500000; y = 0.620000 };

{ width = 0.026667; value = 2000.000000; x = 0.580000; y = 0.620000 };

{ width = 0.026667; value = 2000.000000; x = 0.660000; y = 0.620000 };

{ width = 0.026667; value = 2000.000000; x = 0.740000; y = 0.620000 };

{ width = 0.026667; value = 2000.000000; x = 0.820000; y = 0.620000 };

{ width = 0.026667; value = 2000.000000; x = 0.900000; y = 0.620000 };
{ width = 0.026667; value = 2000.000000; x = 0.980000; y = 0.620000 };
{ width = 0.026667; value = 2000.000000; x = 0.060000; y = 0.680000 };
{ width = 0.026667; value = 2000.000000; x = 0.140000; y = 0.680000 };
{ width = 0.026667; value = 2000.000000; x = 0.220000; y = 0.680000 };
{ width = 0.026667; value = 2000.000000; x = 0.300000; y = 0.680000 };
{ width = 0.026667; value = 2000.000000; x = 0.380000; y = 0.680000 };
{ width = 0.026667; value = 2000.000000; x = 0.460000; y = 0.680000 };
{ width = 0.026667; value = 2000.000000; x = 0.540000; y = 0.680000 };
{ width = 0.026667; value = 2000.000000; x = 0.620000; y = 0.680000 };
{ width = 0.026667; value = 2000.000000; x = 0.700000; y = 0.680000 };
{ width = 0.026667; value = 2000.000000; x = 0.780000; y = 0.680000 };
{ width = 0.026667; value = 2000.000000; x = 0.860000; y = 0.680000 };
{ width = 0.026667; value = 2000.000000; x = 0.940000; y = 0.680000 };
{ width = 0.026667; value = 2000.000000; x = 0.020000; y = 0.740000 };
{ width = 0.026667; value = 2000.000000; x = 0.100000; y = 0.740000 };
{ width = 0.026667; value = 2000.000000; x = 0.180000; y = 0.740000 };
{ width = 0.026667; value = 2000.000000; x = 0.260000; y = 0.740000 };
{ width = 0.026667; value = 2000.000000; x = 0.340000; y = 0.740000 };
{ width = 0.026667; value = 2000.000000; x = 0.420000; y = 0.740000 };
{ width = 0.026667; value = 2000.000000; x = 0.500000; y = 0.740000 };
{ width = 0.026667; value = 2000.000000; x = 0.580000; y = 0.740000 };
{ width = 0.026667; value = 2000.000000; x = 0.660000; y = 0.740000 };
{ width = 0.026667; value = 2000.000000; x = 0.740000; y = 0.740000 };
{ width = 0.026667; value = 2000.000000; x = 0.820000; y = 0.740000 };
{ width = 0.026667; value = 2000.000000; x = 0.900000; y = 0.740000 };
{ width = 0.026667; value = 2000.000000; x = 0.980000; y = 0.740000 };
{ width = 0.026667; value = 2000.000000; x = 0.060000; y = 0.800000 };
{ width = 0.026667; value = 2000.000000; x = 0.140000; y = 0.800000 };
{ width = 0.026667; value = 2000.000000; x = 0.220000; y = 0.800000 };
{ width = 0.026667; value = 2000.000000; x = 0.300000; y = 0.800000 };
{ width = 0.026667; value = 2000.000000; x = 0.380000; y = 0.800000 };

{ width = 0.026667; value = 2000.000000; x = 0.460000; y = 0.800000 };
{ width = 0.026667; value = 2000.000000; x = 0.540000; y = 0.800000 };
{ width = 0.026667; value = 2000.000000; x = 0.620000; y = 0.800000 };
{ width = 0.026667; value = 2000.000000; x = 0.700000; y = 0.800000 };
{ width = 0.026667; value = 2000.000000; x = 0.780000; y = 0.800000 };
{ width = 0.026667; value = 2000.000000; x = 0.860000; y = 0.800000 };
{ width = 0.026667; value = 2000.000000; x = 0.940000; y = 0.800000 };
{ width = 0.026667; value = 2000.000000; x = 0.020000; y = 0.860000 };
{ width = 0.026667; value = 2000.000000; x = 0.100000; y = 0.860000 };
{ width = 0.026667; value = 2000.000000; x = 0.180000; y = 0.860000 };
{ width = 0.026667; value = 2000.000000; x = 0.260000; y = 0.860000 };
{ width = 0.026667; value = 2000.000000; x = 0.340000; y = 0.860000 };
{ width = 0.026667; value = 2000.000000; x = 0.420000; y = 0.860000 };
{ width = 0.026667; value = 2000.000000; x = 0.500000; y = 0.860000 };
{ width = 0.026667; value = 2000.000000; x = 0.580000; y = 0.860000 };
{ width = 0.026667; value = 2000.000000; x = 0.660000; y = 0.860000 };
{ width = 0.026667; value = 2000.000000; x = 0.740000; y = 0.860000 };
{ width = 0.026667; value = 2000.000000; x = 0.820000; y = 0.860000 };
{ width = 0.026667; value = 2000.000000; x = 0.900000; y = 0.860000 };
{ width = 0.026667; value = 2000.000000; x = 0.980000; y = 0.860000 };
{ width = 0.026667; value = 2000.000000; x = 0.060000; y = 0.920000 };
{ width = 0.026667; value = 2000.000000; x = 0.140000; y = 0.920000 };
{ width = 0.026667; value = 2000.000000; x = 0.220000; y = 0.920000 };
{ width = 0.026667; value = 2000.000000; x = 0.300000; y = 0.920000 };
{ width = 0.026667; value = 2000.000000; x = 0.380000; y = 0.920000 };
{ width = 0.026667; value = 2000.000000; x = 0.460000; y = 0.920000 };
{ width = 0.026667; value = 2000.000000; x = 0.540000; y = 0.920000 };
{ width = 0.026667; value = 2000.000000; x = 0.620000; y = 0.920000 };
{ width = 0.026667; value = 2000.000000; x = 0.700000; y = 0.920000 };
{ width = 0.026667; value = 2000.000000; x = 0.780000; y = 0.920000 };
{ width = 0.026667; value = 2000.000000; x = 0.860000; y = 0.920000 };
{ width = 0.026667; value = 2000.000000; x = 0.940000; y = 0.920000 };

```
{ width = 0.026667; value = 2000.000000; x = 0.020000; y = 0.980000 };
{ width = 0.026667; value = 2000.000000; x = 0.100000; y = 0.980000 };
{ width = 0.026667; value = 2000.000000; x = 0.180000; y = 0.980000 };
{ width = 0.026667; value = 2000.000000; x = 0.260000; y = 0.980000 };
{ width = 0.026667; value = 2000.000000; x = 0.340000; y = 0.980000 };
{ width = 0.026667; value = 2000.000000; x = 0.420000; y = 0.980000 };
{ width = 0.026667; value = 2000.000000; x = 0.500000; y = 0.980000 };
{ width = 0.026667; value = 2000.000000; x = 0.580000; y = 0.980000 };
{ width = 0.026667; value = 2000.000000; x = 0.660000; y = 0.980000 };
{ width = 0.026667; value = 2000.000000; x = 0.740000; y = 0.980000 };
{ width = 0.026667; value = 2000.000000; x = 0.820000; y = 0.980000 };
{ width = 0.026667; value = 2000.000000; x = 0.900000; y = 0.980000 };
{ width = 0.026667; value = 2000.000000; x = 0.980000; y = 0.980000 };
] } }
```

```
| F1Q1 ->{kdis2} F1Act + Q1A
| psA + Q1A <->{(3/rprot)*P}{(3/rprot)*P} Q1sol + psA

| psB + Q1sol <->{(3/rprot)*P}{(3/rprot)*P} Q1B + psB
| Q1B + F2Q2 ->{kdis} F2Q1
```

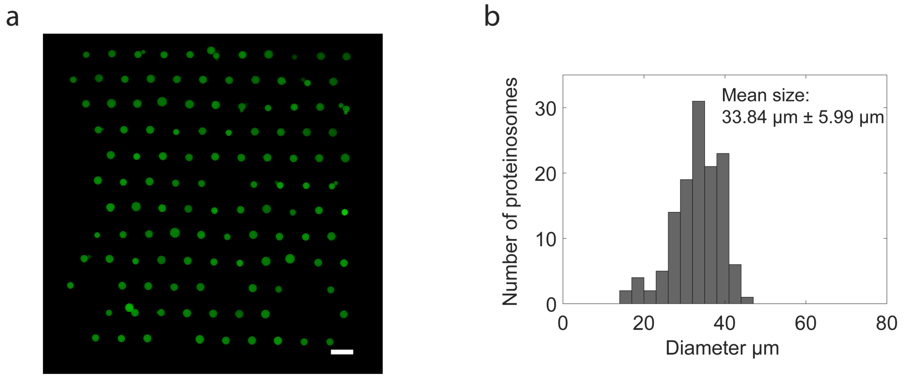

**Supplementary Figure 1.** Protocellular size distribution. (a) Confocal micrograph of FITC-labeled proteinosomes localized in the microfluidic trapping device. Scale bar 100 μm. (b) Typical size distribution of the trapped proteinosomes.

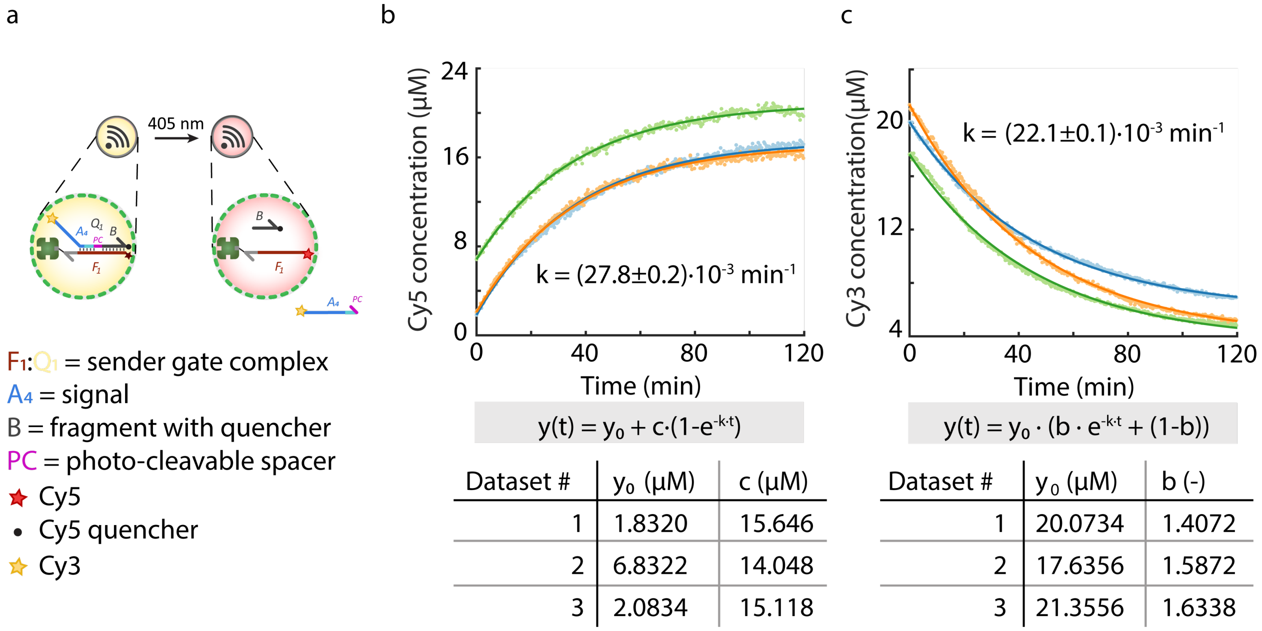

$$y(t) = y_0 + c\cdot(1-e^{-k\cdot t})$$

| Dataset # | $y_0$ (µM) | c (µM) |
|---|---|---|
| 1 | 1.8320 | 15.646 |
| 2 | 6.8322 | 14.048 |
| 3 | 2.0834 | 15.118 |

$$y(t) = y_0 \cdot (b \cdot e^{-k\cdot t} + (1-b))$$

| Dataset # | $y_0$ (µM) | b (-) |
|---|---|---|
| 1 | 20.0734 | 1.4072 |
| 2 | 17.6356 | 1.5872 |
| 3 | 21.3556 | 1.6338 |

**Supplementary Figure 2.** (a) Photo-cleavage reaction in proteinosomes. Upon cleavage of the photo-cleavable spacer (**PC**) by laser light irradiation (405 nm), the signal (strand $\mathbf{A_4}$) and Cy5 quencher (strand **B**) are released. This reaction was monitored through unquenching of Cy5 and release of Cy3 from the proteinosomes. (b) Cy5 fluorescence in three distinct proteinosomes over time upon irradiation by laser (405 nm, 2 h). The photo-cleavage was modeled as a first-order reaction, yielding the exponential equation shown in the figure. Since proteinosomes are heterogeneous in size, background fluorescence and DNA loading capacity, the background ($y_0$) was determined from the first timepoint for each proteinosome individually. In addition, a unique parameter *c* (concentration of reactive sender gate complex) was fitted for each proteinosome, whilst the rate of release (*k*), which should be independent of proteinosome characteristics, was fitted as a single parameter for all datasets. (c) Similarly, the release of the Cy3 labeled signal strand was modeled using first-order kinetics. Now the fluorescence at the initial timepoint ($y_0$) indicates the total concentration of complex plus background fluorescence. The parameter *b*, which is fitted as unique parameter for each dataset, signifies the fraction of the total fluorescence that is accounted for by the sender gate complex, making 1-*b* the fraction originating from background fluorescence. Experiments were performed in independent triplicates.

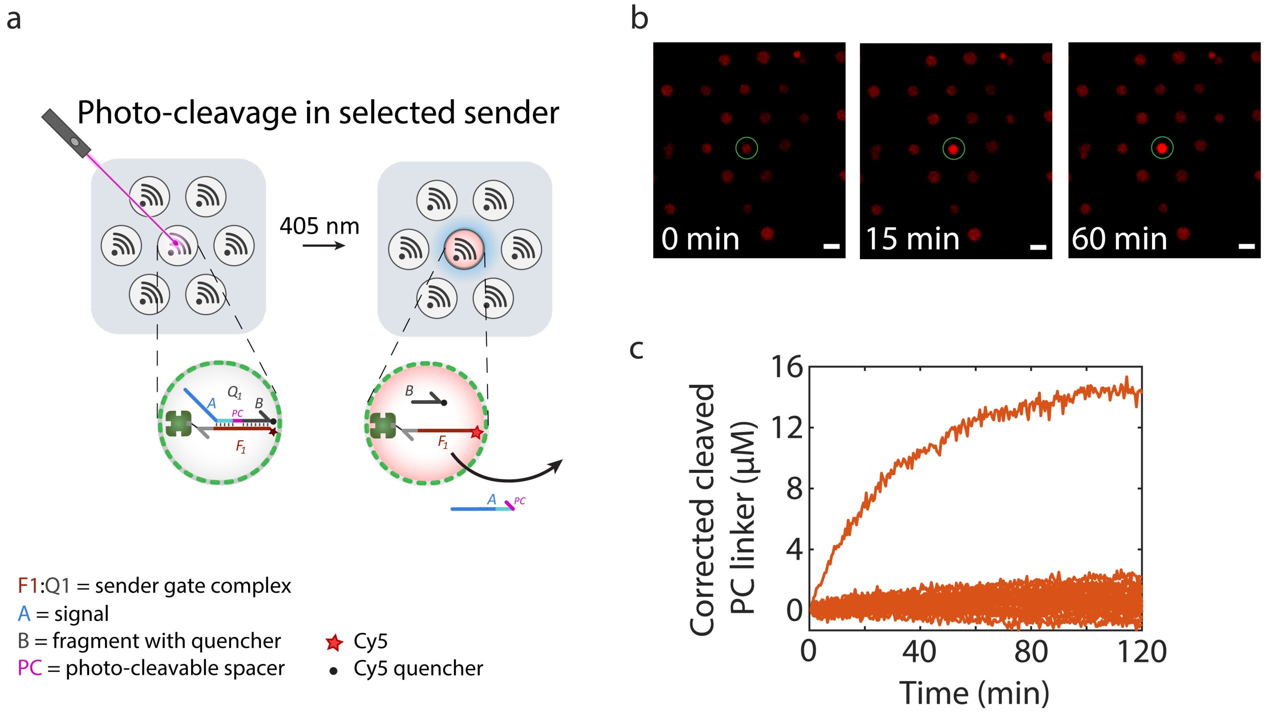

**Supplementary Figure 3.** Photo-cleavage of **PC** linker in selected sender protocell. (a) A population of sender protocells were loaded into the trapping device and a single sender protocell was illuminated with laser light (405 nm, 100 ms every 1 s). (b) Confocal micrographs of sender protocells showing time-dependent increase in Cy5 fluorescence of the irradiated sender due to the cleavage of $\mathbf{Q_7}$ and dissociation of quencher-modified strand **B**. Scale bar 50 μm. (c) Time-traces of cleaved **PC** linker in sender protocells through selective light irradiation. The concentration of cleaved **PC** linker was obtained through conversion of measured Cy5 fluorescence using calibration values of $\mathbf{F_1}$ strand. The data shows that only the irradiated sender protocells shows a time-dependent increase in concentration of cleaved duplex, indicating selective photo-cleavage. Sender protocells were prepared using 10 μM streptavidin.

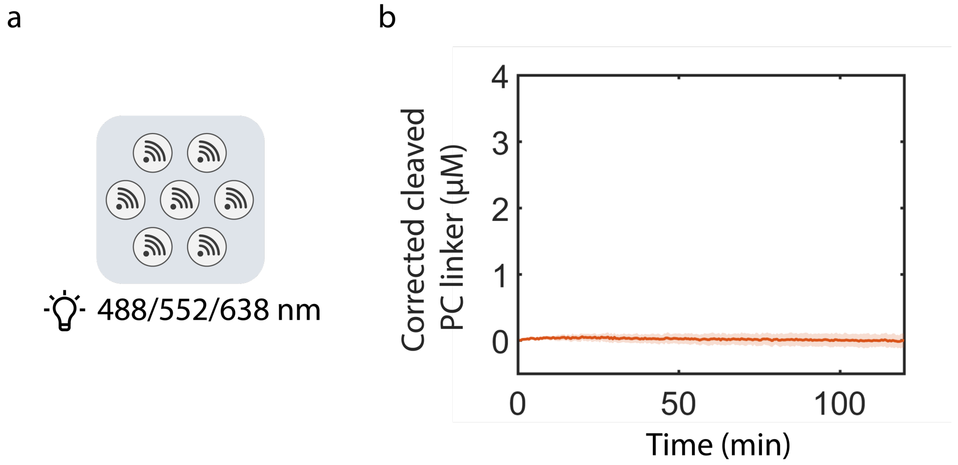

**Supplementary Figure 4.** Effect of light with wavelengths other than 405 nm on cleavage of the photocleavable linker. (a) A population of sender protocells was illuminated with light (488 nm, 552 nm, 638 nm). The respective lasers were pulsed on alternatively for 100 ms every 18 s. (b) Mean and standard deviation of time-traces of cleaved **PC** linker in sender protocells showing that cleavage of the photocleavable linker does not occur upon irradiation with the tested alternative wavelengths. Sender protocells were prepared using 10 µM streptavidin.

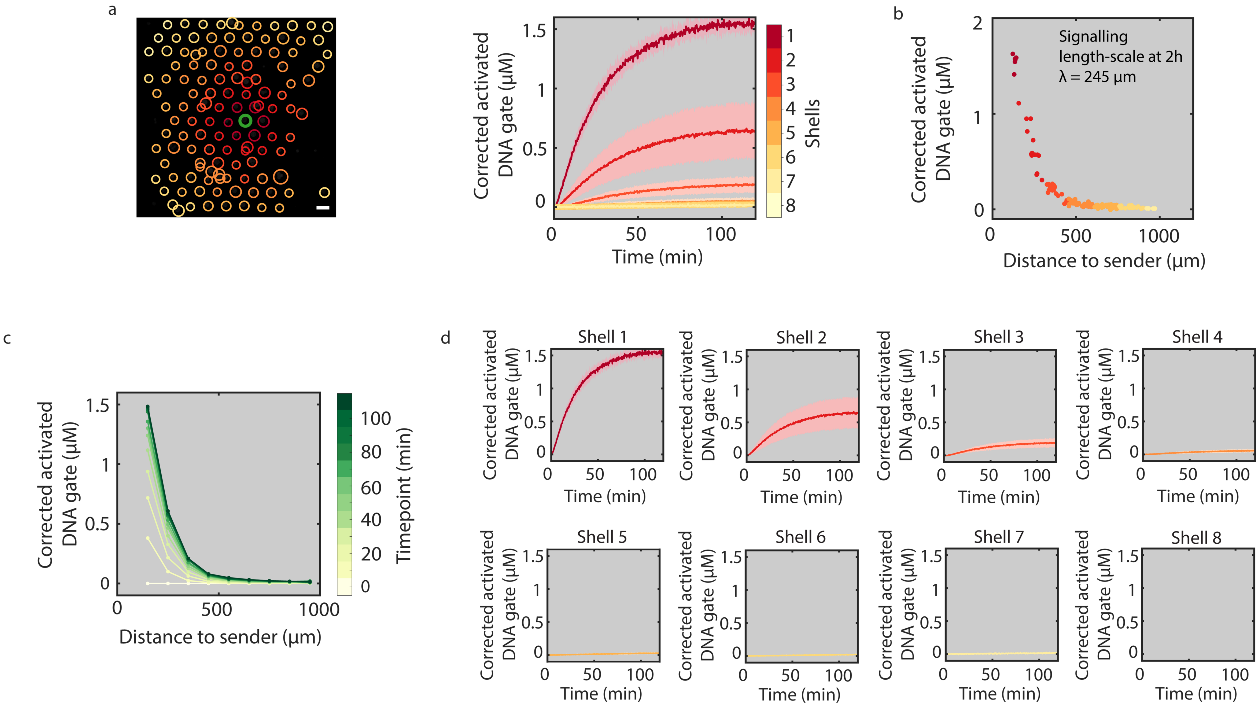

**Supplementary Figure 5.** Quantification of receiver activation in full detail. (a) Time-traces of receiver activation including the standard deviation for each shell (shaded regions). (b) Concentrations of activated DNA gates in individual receivers after 2 h of signal release as function of the distance to the sender protocell and the calculated signaling length-scale. (c) The concentration of activated gate complexes in receiver protocells is plotted for different times of the reaction as a function of distance to the sender. (d) Time-traces of mean receiver activity per shell including the standard deviation (shaded region). This data corresponds to the high consumption capacity (Fig. 2b, data set 1/3) and high density (Fig. 2d, data set 1/3) experiments. All conditions are summarized in Supplementary Table 2, entry 1. Scale bar 100 µm.

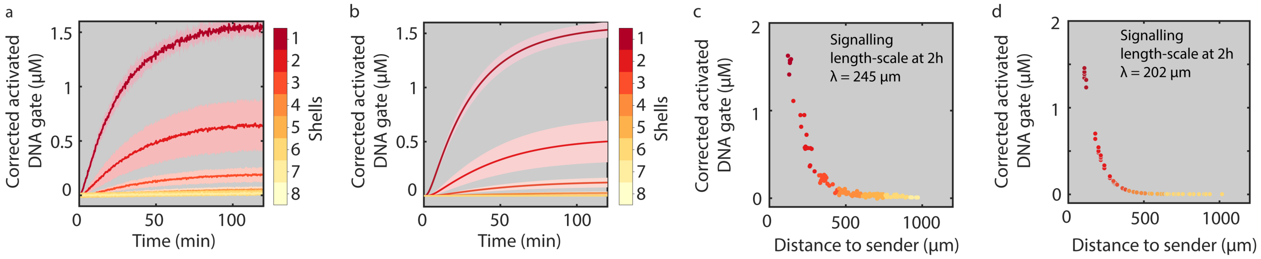

**Supplementary Figure 6.** Experimental and simulated data of the sender-receiver system obtained by 2D reaction-diffusion simulations. Experimental (a) and simulated (b) time traces of activated receiver protocells in different shells including standard deviation. The standard deviation per shell in the model originates from the different distance of individual receiver protocells from the sender protocell. Experimental (c) and simulated (d) concentrations of activated DNA gates in individual receivers after 2 h of signal release as function of the distance to the sender protocell and the calculated signaling length-scales. The reactions and parameters used in the model are shown in Supplementary Methods and Table 4.

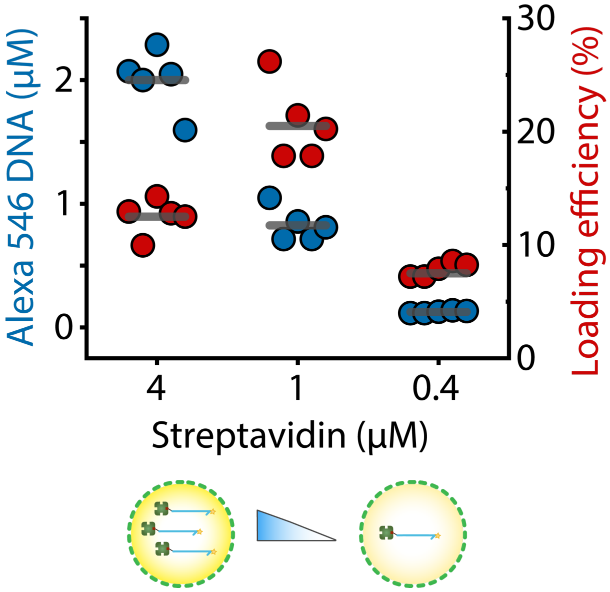

**Supplementary Figure 7.** DNA localization capacity of proteinosomes (Alexa 546-DNA concentration, blue dots) prepared using a range of streptavidin concentrations. The streptavidin concentration corresponds to the concentration of streptavidin present in the aqueous phase during proteinosomes assembly. In addition, loading efficiencies (red dots) were obtained under assumption that each streptavidin can bind four biotinylated ssDNA strands at maximum. The three protocell populations were used to investigate receiver activation with different receiver consumption capacities (Fig. 2b).

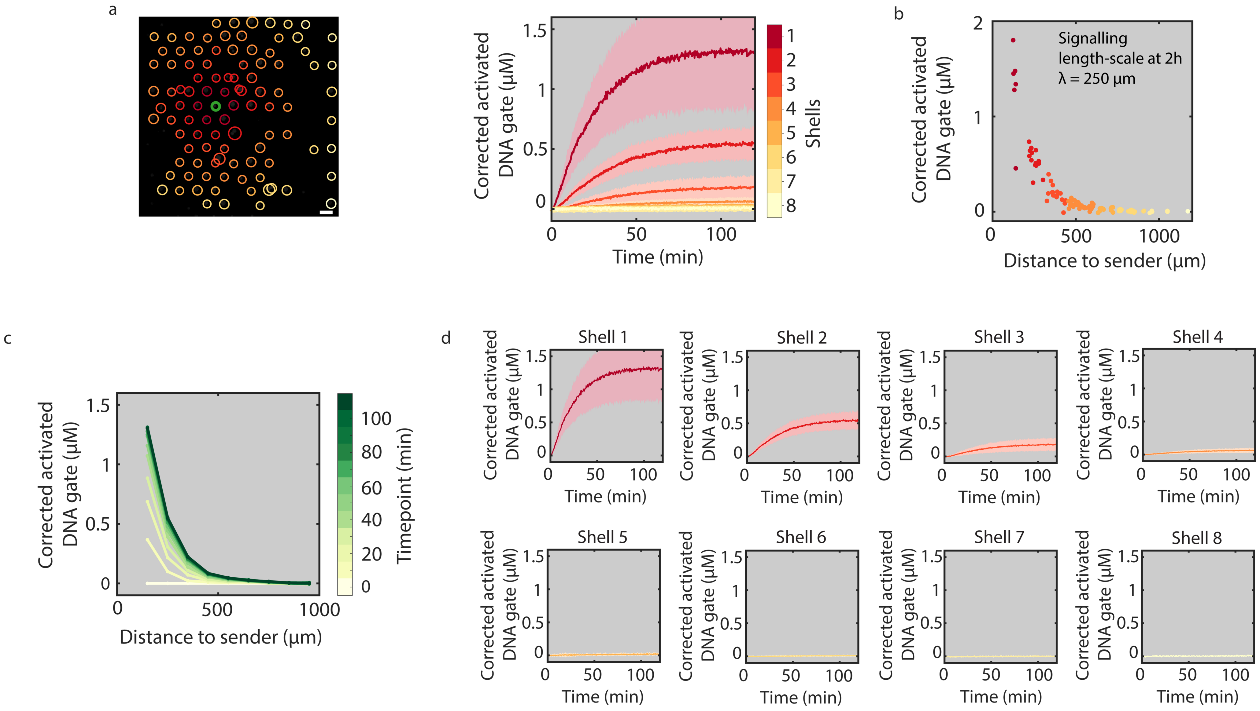

**Supplementary Figure 8.** Receiver activation corresponding to high consumption capacity (data set 2/3). (a) Time-traces of receiver activation including the standard deviation for each shell (shaded regions). (b) Concentrations of activated DNA gates in individual receivers after 2 h of signal release as function of the distance to the sender protocell and the calculated signaling length-scale. (c) The concentration of activated gate complexes in receiver protocells is plotted for different times of the reaction as a function of distance to the sender. (d) Time-traces of mean receiver activity per shell including the standard deviation (shaded region). This data corresponds to the high-density experiments (Fig. 2d, data set 2/3). All conditions are summarized in Supplementary Table 2, entry 1. Scale bar 100 µm.

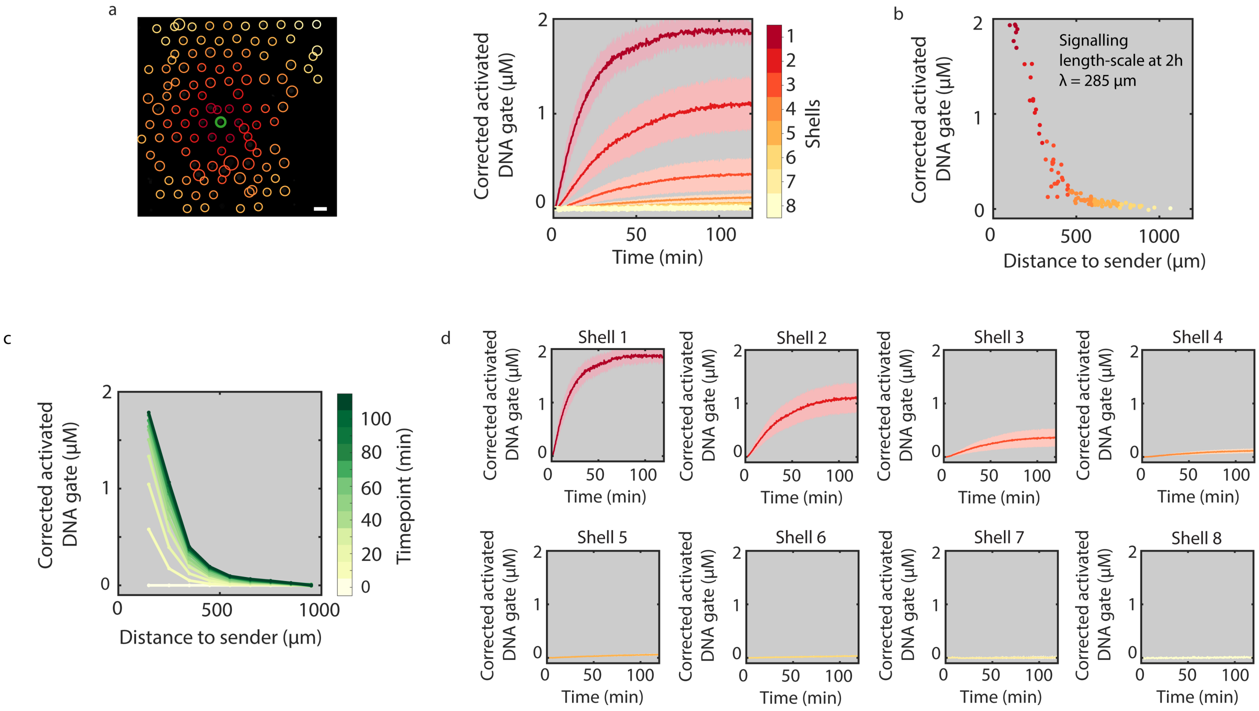

**Supplementary Figure 9.** Receiver activation corresponding to high consumption capacity (data set 3/3). (a) Time-traces of receiver activation including the standard deviation for each shell (shaded regions). (b) Concentrations of activated DNA gates in individual receivers after 2 h of signal release as function of the distance to the sender protocell and the calculated signaling length-scale. (c) The concentration of activated gate complexes in receiver protocells is plotted for different times of the reaction as a function of distance to the sender. (d) Time-traces of mean receiver activity per shell including the standard deviation (shaded region). This data corresponds to the high-density experiments (Fig. 2d, data set 3/3). All conditions are summarized in Supplementary Table 2, entry 1. Scale bar 100 μm.

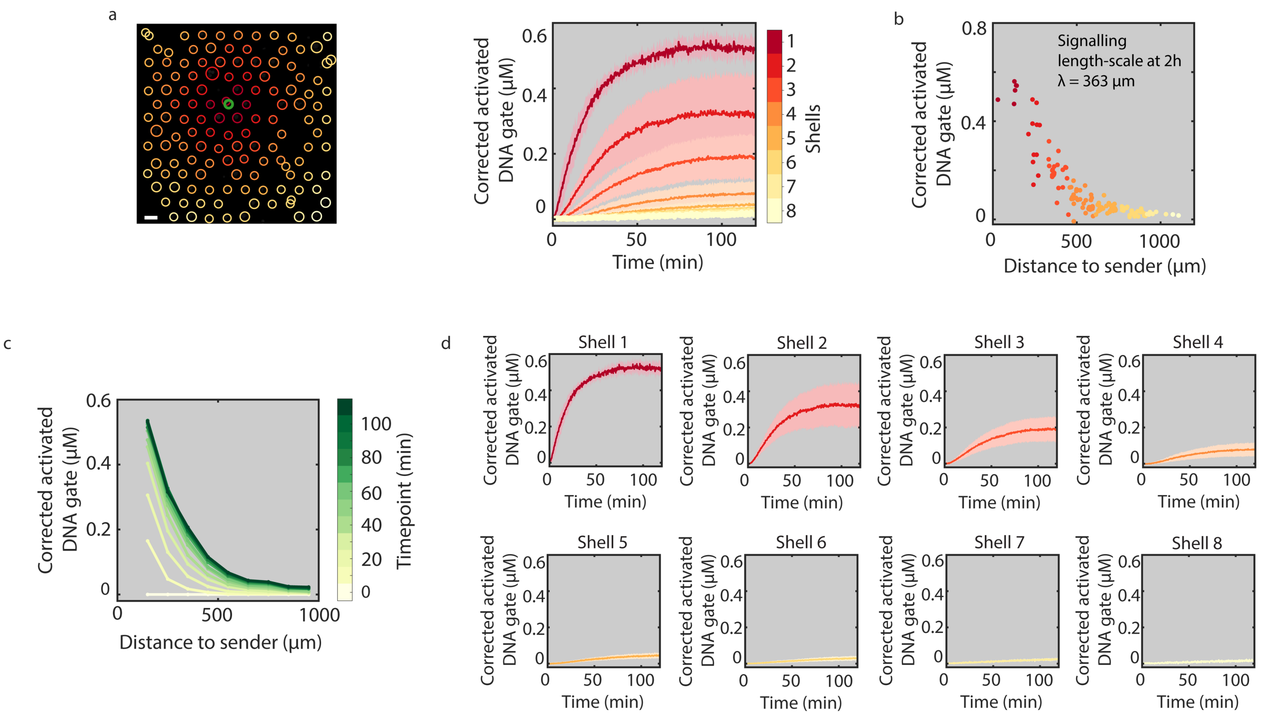

**Supplementary Figure 10.** Receiver activation corresponding to medium consumption capacity (data set 1/3). (a) Time-traces of receiver activation including the standard deviation for each shell (shaded regions). (b) Concentrations of activated DNA gates in individual receivers after 2 h of signal release as function of the distance to the sender protocell and the calculated signaling length-scale. (c) The concentration of activated gate complexes in receiver protocells is plotted for different times of the reaction as a function of distance to the sender. (d) Time-traces of mean receiver activity per shell including the standard deviation (shaded region). This data corresponds to the high permeability experiments (Fig. 2c, data set 1/3) and is used as a control for the exonuclease experiments (Fig. 2e, data set 1/3). All conditions are summarized in Supplementary Table 2, entry 2. Scale bar 100 µm.

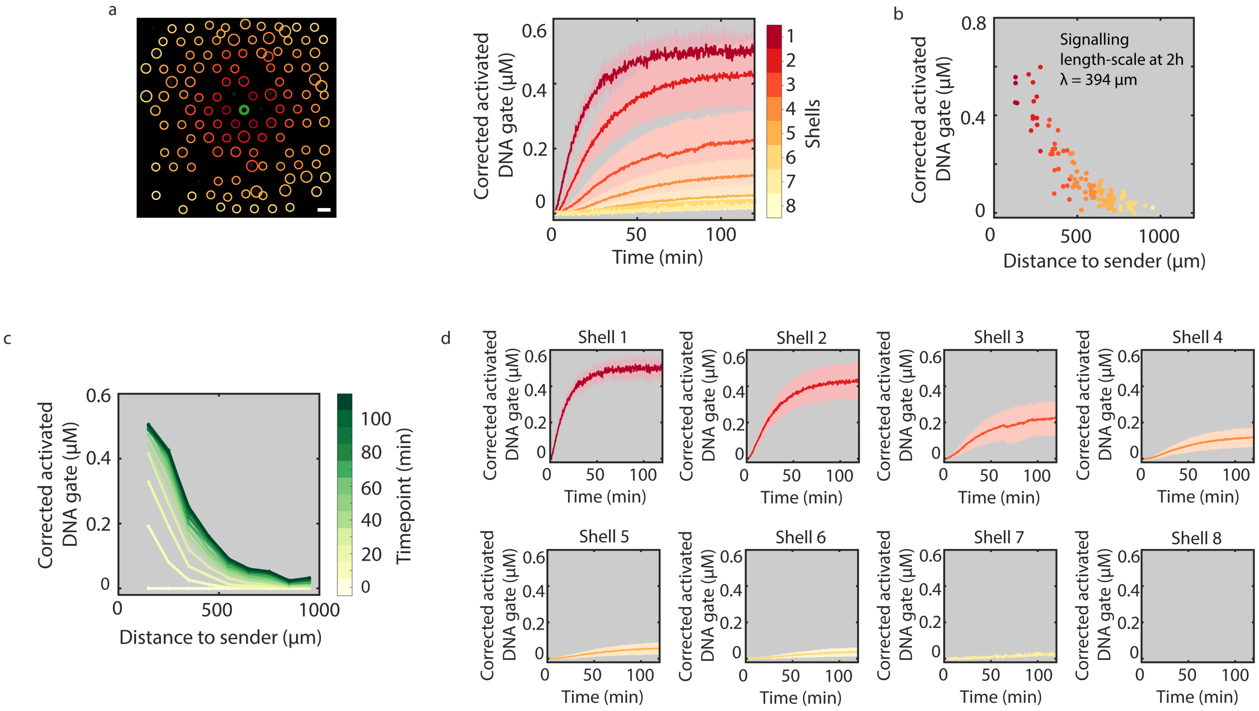

**Supplementary Figure 11.** Receiver activation corresponding to medium consumption capacity (data set 2/3). (a) Time-traces of receiver activation including the standard deviation for each shell (shaded regions). (b) Concentrations of activated DNA gates in individual receivers after 2 h of signal release as function of the distance to the sender protocell and the calculated signaling length-scale. (c) The concentration of activated gate complexes in receiver protocells is plotted for different times of the reaction as a function of distance to the sender. (d) Time-traces of mean receiver activity per shell including the standard deviation (highlighted region). This data corresponds to the high permeability experiments (Fig. 2c, data set 2/3) and is used as a control for the exonuclease experiments (Fig. 2e, data set 2/3). All conditions are summarized in Supplementary Table 2, entry 2. Scale bar 100 μm.

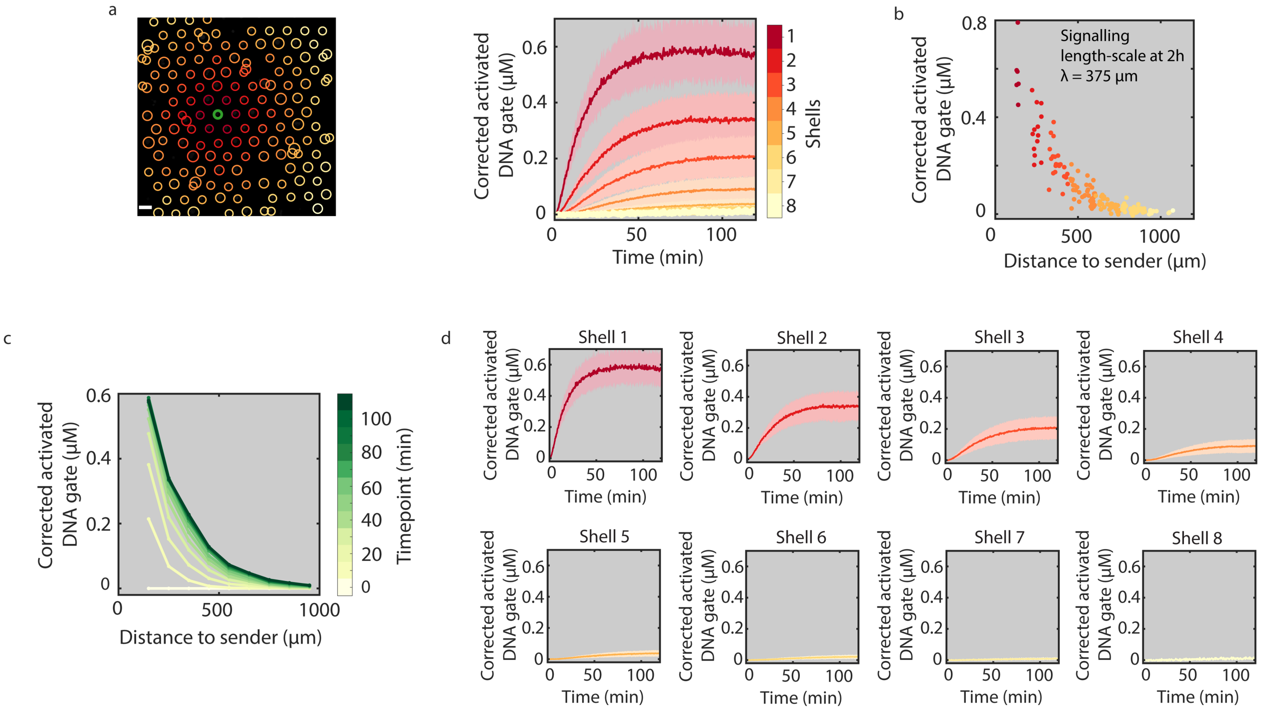

**Supplementary Figure 12.** Receiver activation corresponding to medium consumption capacity (data set 3/3). (a) Time-traces of receiver activation including the standard deviation for each shell (shaded regions). (b) Concentrations of activated DNA gates in individual receivers after 2 h of signal release as function of the distance to the sender protocell and the calculated signaling length-scale. (c) The concentration of activated gate complexes in receiver protocells is plotted for different times of the reaction as a function of distance to the sender. (d) Time-traces of mean receiver activity per shell including the standard deviation (shaded region). This data corresponds to the high permeability experiments (Fig. 2c, data set 3/3) and is used as a control for the exonuclease experiments (Fig. 2e, data set 3/3). All conditions are summarized in Supplementary Table 2, entry 2. Scale bar 100 µm.

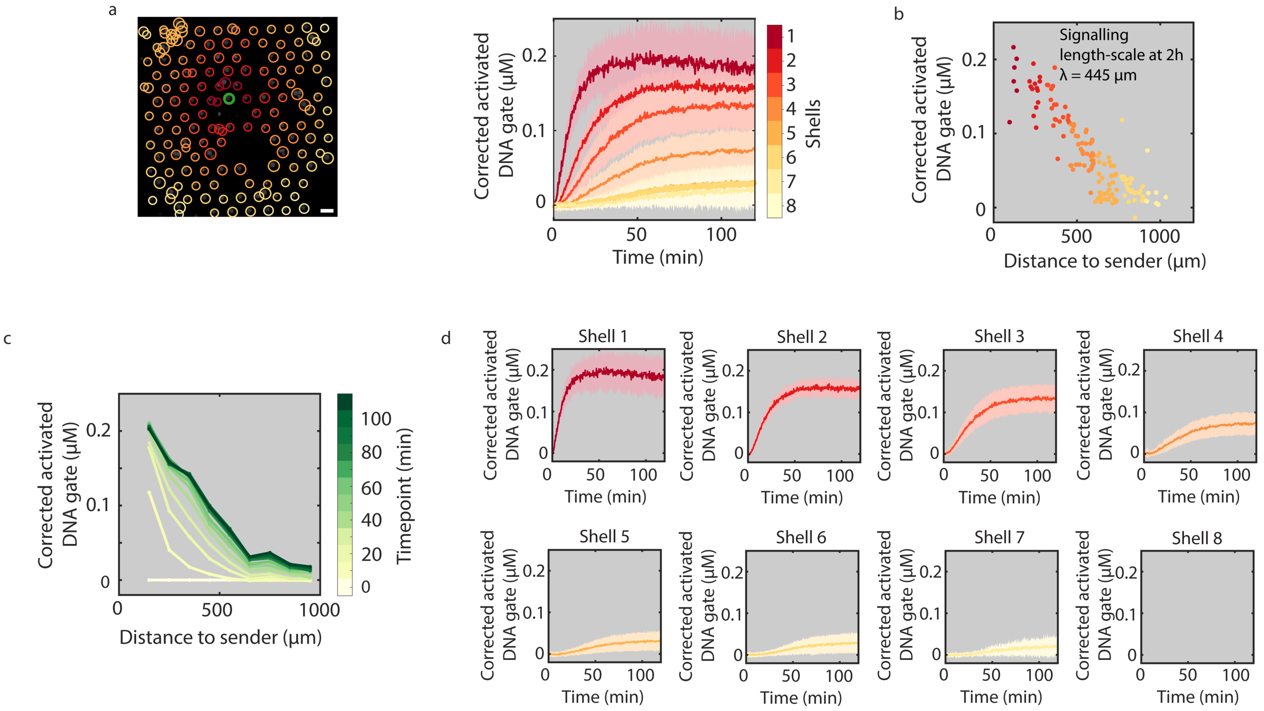

**Supplementary Figure 13.** Receiver activation corresponding to low consumption capacity (data set 1/3). (a) Time-traces of receiver activation including the standard deviation for each shell (shaded regions). (b) Concentrations of activated DNA gates in individual receivers after 2 h of signal release as function of the distance to the sender protocell and the calculated signaling length-scale. (c) The concentration of activated gate complexes in receiver protocells is plotted for different times of the reaction as a function of distance to the sender. (d) Time-traces of mean receiver activity per shell including the standard deviation (shaded region). All conditions are summarized in Supplementary Table 2, entry 3. Scale bar 100 µm.

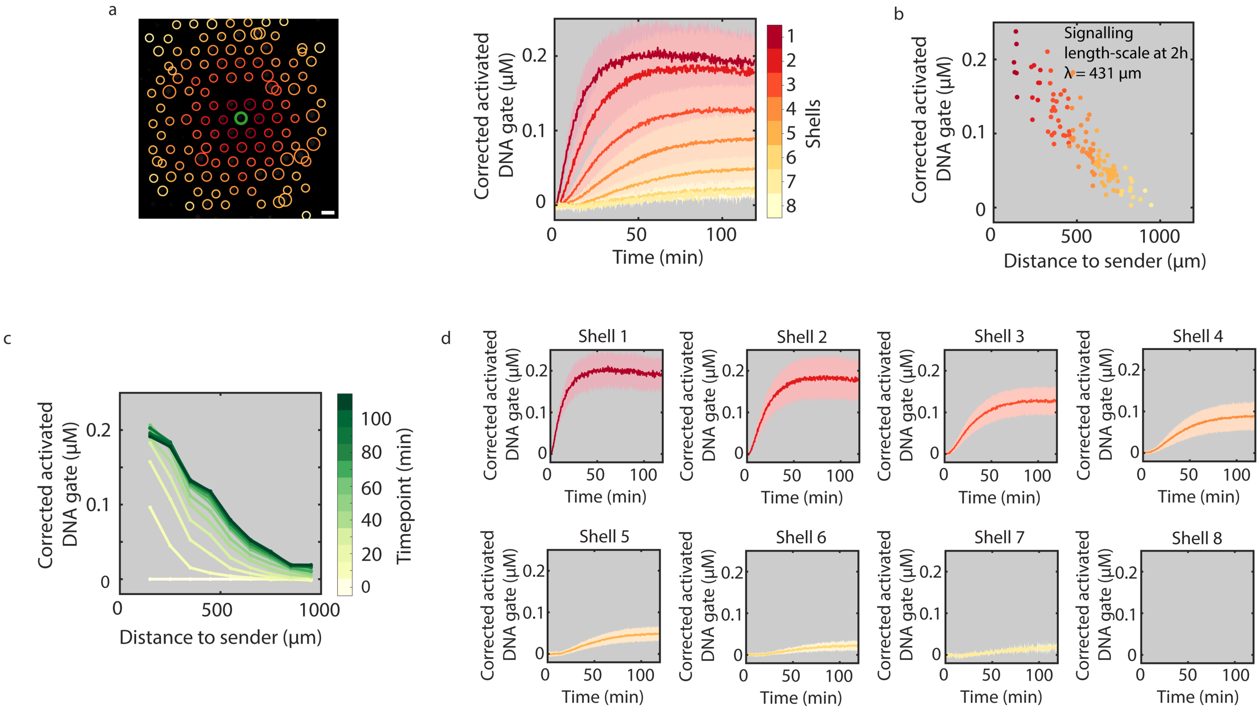

**Supplementary Figure 14.** Receiver activation corresponding to low consumption capacity (data set 2/3). (a) Time-traces of receiver activation including the standard deviation for each shell (shaded regions). (b) Concentrations of activated DNA gates in individual receivers after 2 h of signal release as function of the distance to the sender protocell and the calculated signaling length-scale. (c) The concentration of activated gate complexes in receiver protocells is plotted for different times of the reaction as a function of distance to the sender. (d) Time-traces of mean receiver activity per shell including the standard deviation (shaded region). All conditions are summarized in Supplementary Table 2, entry 3. Scale bar 100 µm.

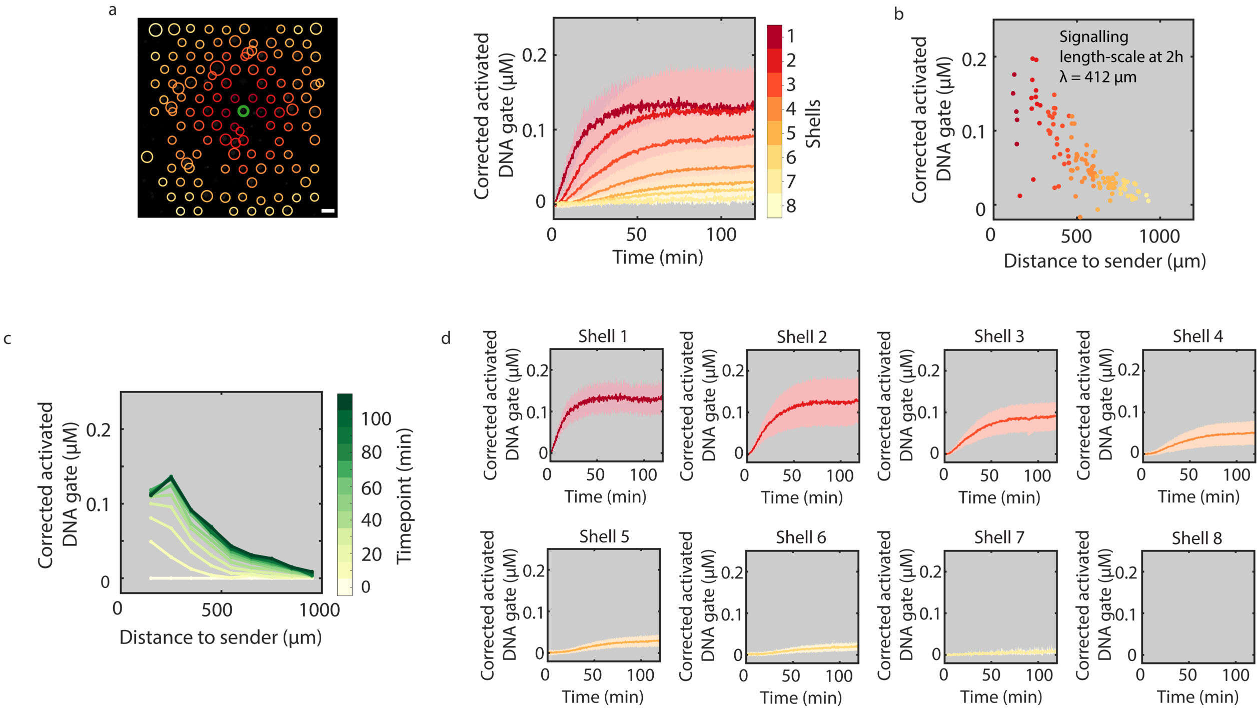

**Supplementary Figure 15.** Receiver activation corresponding to low consumption capacity (data set 3/3). (a) Time-traces of receiver activation including the standard deviation for each shell (shaded regions). (b) Concentrations of activated DNA gates in individual receivers after 2 h of signal release as function of the distance to the sender protocell and the calculated signaling length-scale. (c) The concentration of activated gate complexes in receiver protocells is plotted for different times of the reaction as a function of distance to the sender. (d) Time-traces of mean receiver activity per shell including the standard deviation (shaded region). All conditions are summarized in Supplementary Table 2, entry 3. Scale bar 100 µm.

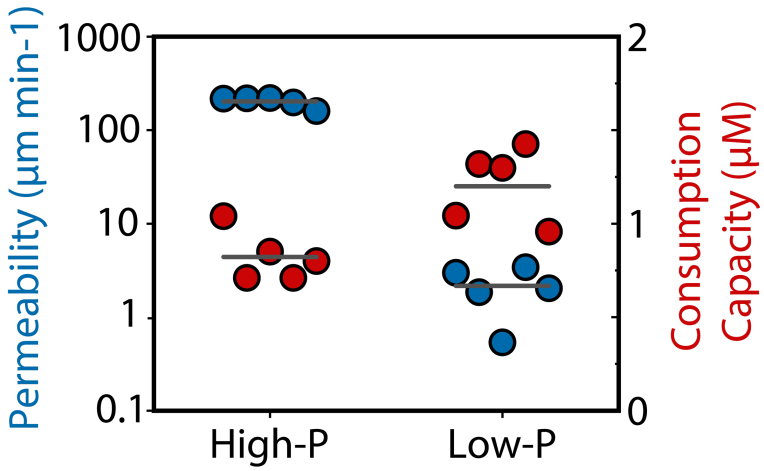

**Supplementary Figure 16.** Permeability (blue dots) and consumption capacity (red dots) of high-P and low-P proteinosomes. Permeability constants were estimated as reported previously[5]. The consumption capacity was determined through fluorescence measurements of proteinosomes that were loaded with Alexa 546-modified DNA. The two protocell populations were used to investigate receiver activation with different permeabilities (Fig. 2c).

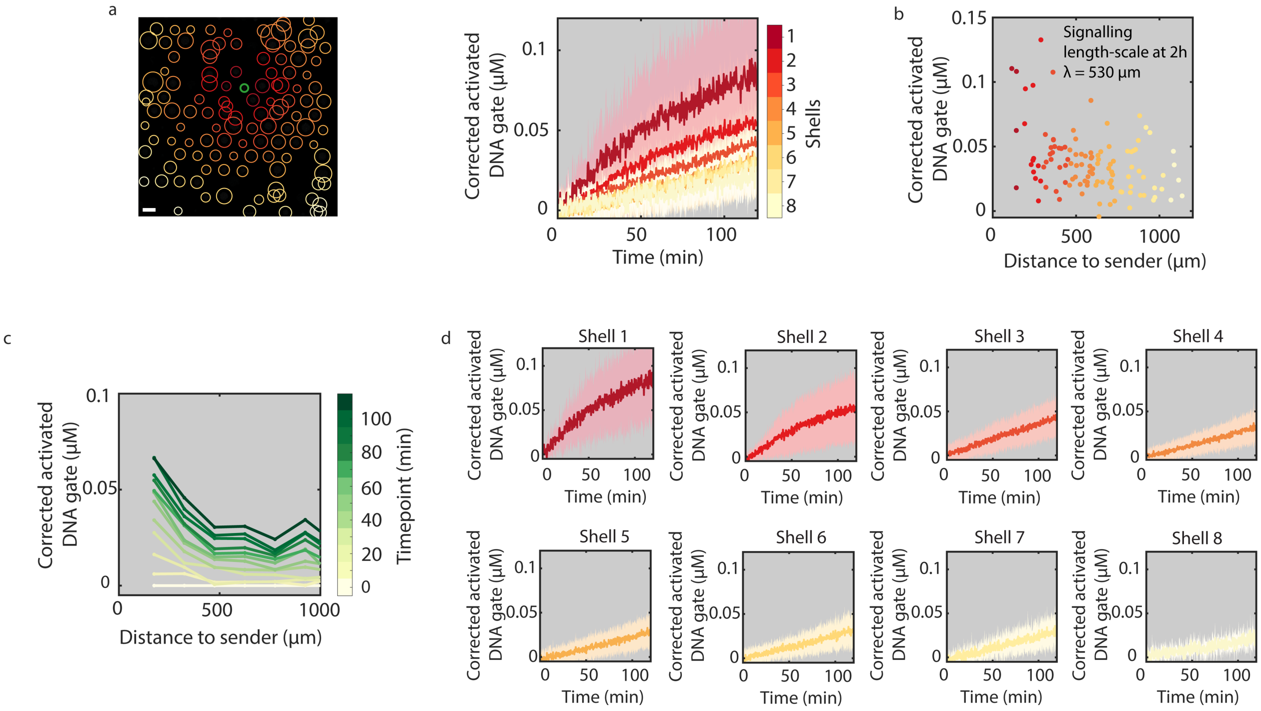

**Supplementary Figure 17.** Activation of receiver protocells corresponding to low permeability (data set 1/3). (a) Time-traces of receiver activation including the standard deviation for each shell (shaded regions). (b) Concentrations of activated DNA gates in individual receivers after 2 h of signal release as function of the distance to the sender protocell and the calculated signaling length-scale. (c) The concentration of activated gate complexes in receiver protocells is plotted for different times of the reaction as a function of distance to the sender. (d) Time-traces of mean receiver activity per shell including the standard deviation (shaded region). All conditions are summarized in Supplementary Table 2, entry 4. Scale bar 100 μm.

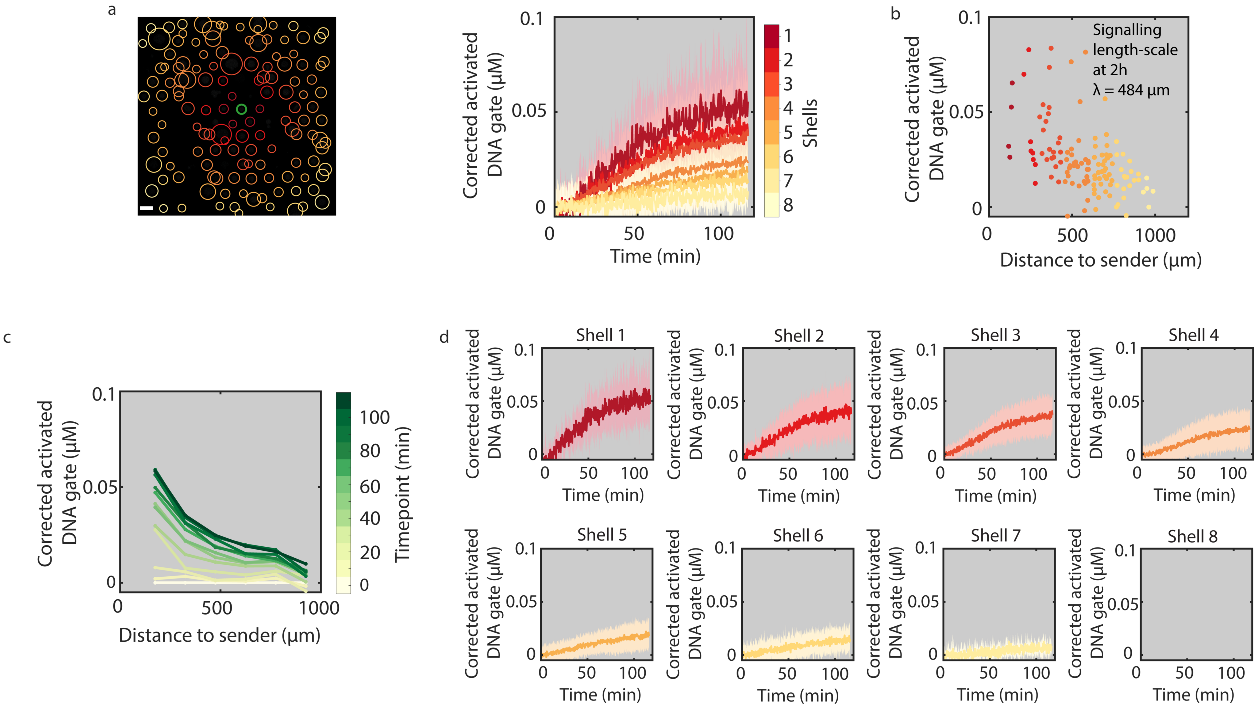

**Supplementary Figure 18.** Activation of receiver protocells corresponding to low permeability (data set 2/3). (a) Time-traces of receiver activation including the standard deviation for each shell (shaded regions). (b) Concentrations of activated DNA gates in individual receivers after 2 h of signal release as function of the distance to the sender protocell and the calculated signaling length-scale. (c) The concentration of activated gate complexes in receiver protocells is plotted for different times of the reaction as a function of distance to the sender. (d) Time-traces of mean receiver activity per shell including the standard deviation (shaded region). All conditions are summarized in Supplementary Table 2, entry 4. Scale bar 100 µm.

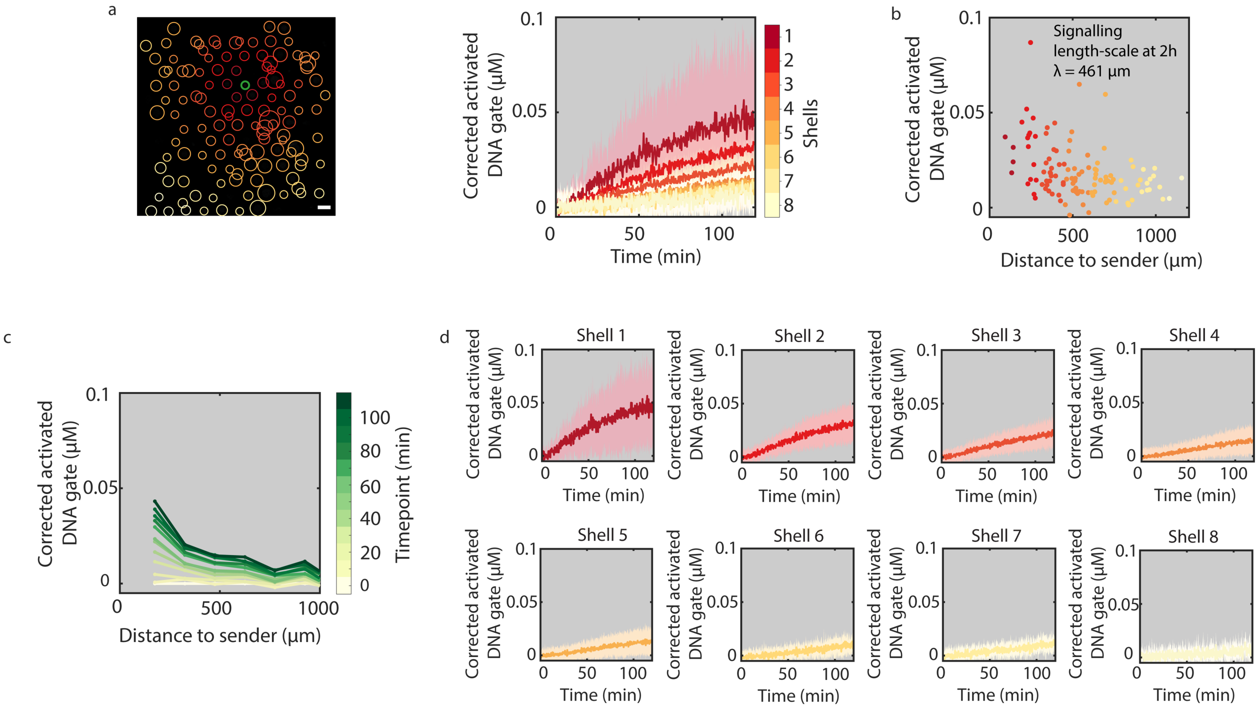

**Supplementary Figure 19.** Activation of receiver protocells corresponding to low permeability (data set 3/3).(a) Time-traces of receiver activation including the standard deviation for each shell (shaded regions). (b) Concentrations of activated DNA gates in individual receivers after 2 h of signal release as function of the distance to the sender protocell and the calculated signaling length-scale. (c) The concentration of activated gate complexes in receiver protocells is plotted for different times of the reaction as a function of distance to the sender. (d) Time-traces of mean receiver activity per shell including the standard deviation (shaded region). All conditions are summarized in Supplementary Table 2, entry 4. Scale bar 100 µm.

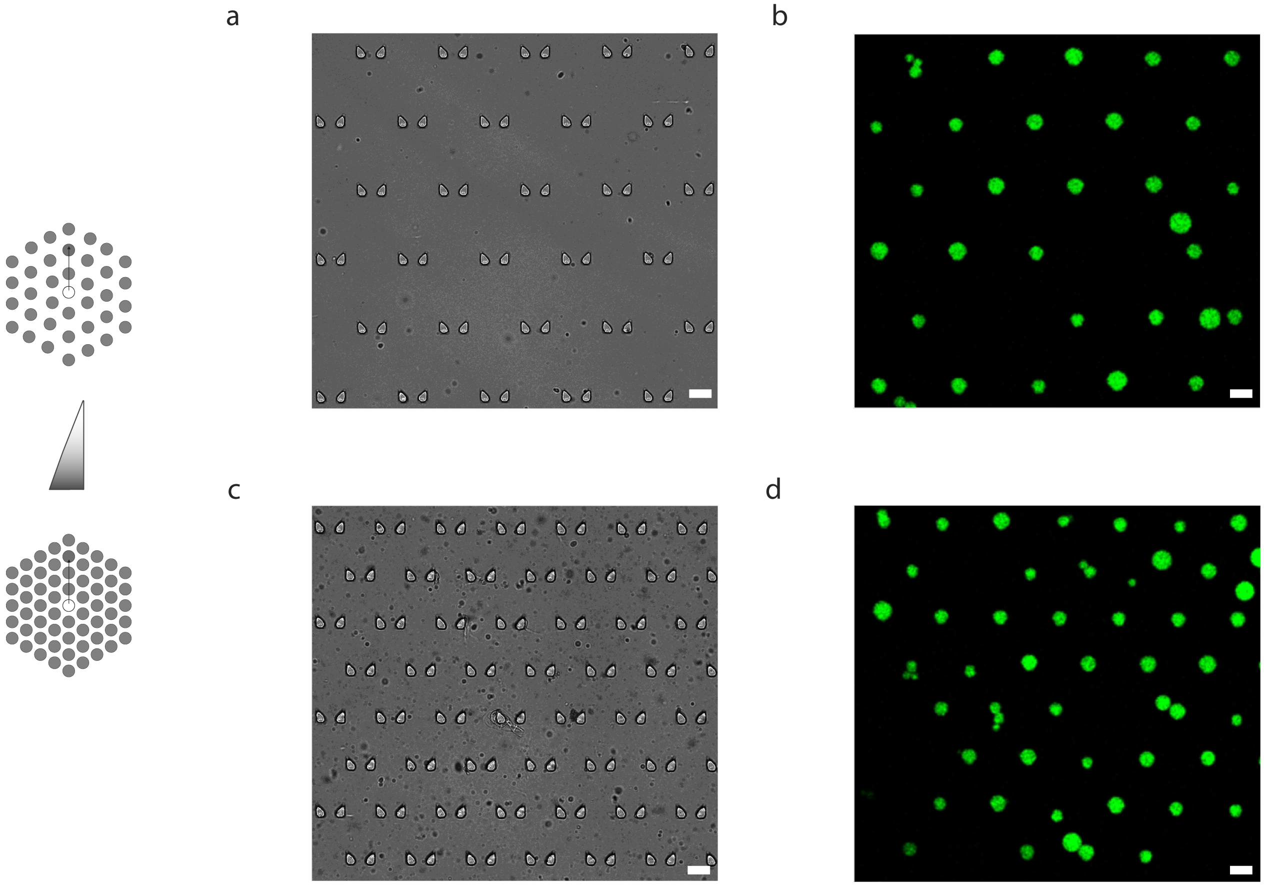

**Supplementary Figure 20.** Bright-field micrograph of low-density (a) and high-density (c) trapping array. Fluorescence micrograph of FITC-labeled proteinosomes in a low-density (b) and high-density (d) trapping array. Scale bar 50 µm. The high and low density device contain 270 and 210 traps, respectively, in a 2 mm X 1.5 mm localization chamber. The two trap arrays were used to study receiver activation for different protocell densities (Fig. 2d).

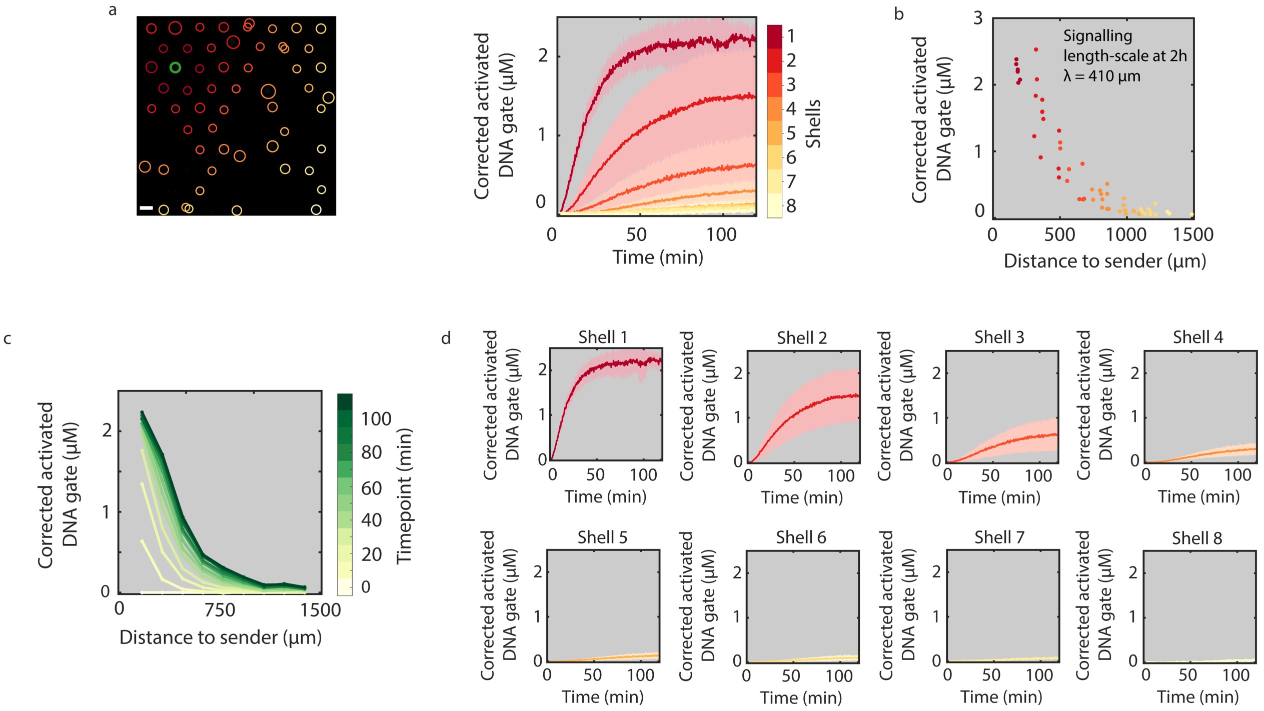

**Supplementary Figure 21.** Activation of receiver protocells corresponding to low protocell density (data set 1/3). (a) Time-traces of receiver activation including the standard deviation (shaded regions). (b) Concentrations of activated DNA gates in individual receivers after 2 h of signal release as function of the distance to the sender protocell and the calculated signaling length-scale. (c) The concentration of activated gate complexes in receiver protocells is plotted for different times of the reaction as a function of distance to the sender. (d) Time-traces of mean receiver activity per shell including the standard deviation (shaded region). All conditions are summarized in Supplementary Table 2, entry 5. Scale bar 100 µm.

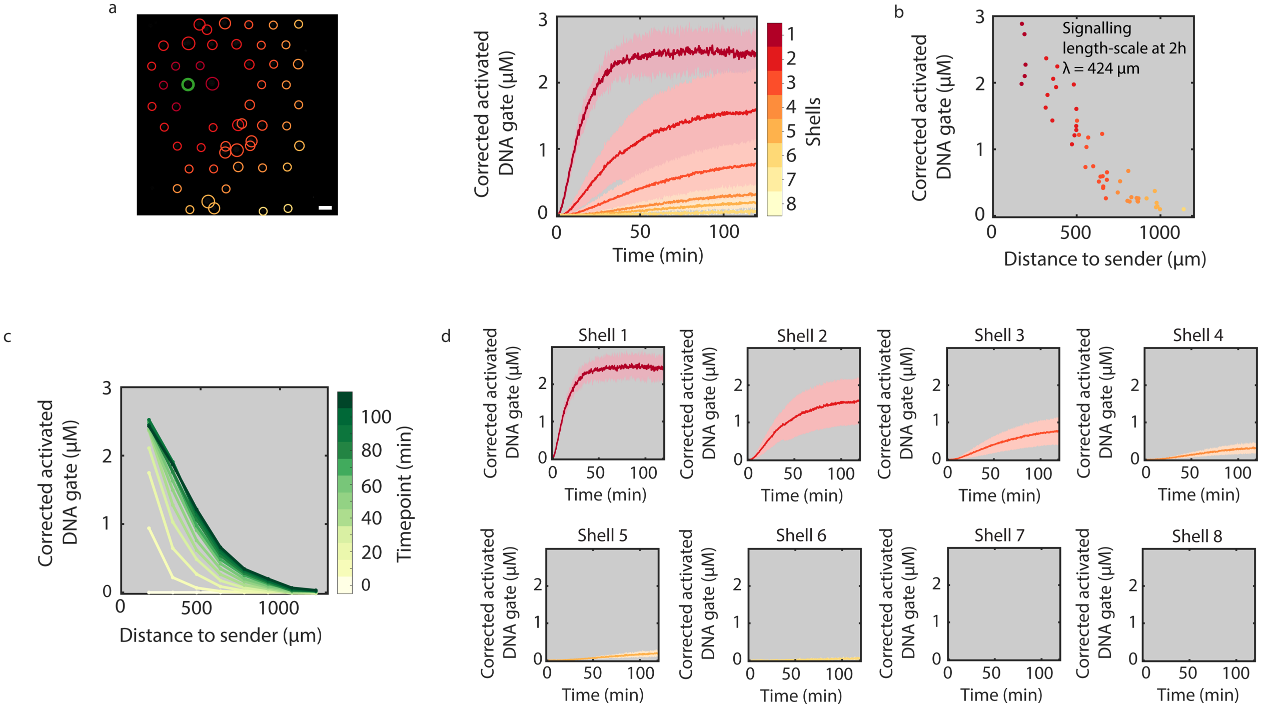

**Supplementary Figure 22.** Activation of receiver protocells corresponding to low protocell density (data set 2/3).(a) Time-traces of receiver activation including the standard deviation (shaded regions). (b) Concentrations of activated DNA gates in individual receivers after 2 h of signal release as function of the distance to the sender protocell and the calculated signaling length-scale. (c) The concentration of activated gate complexes in receiver protocells is plotted for different times of the reaction as a function of distance to the sender. (d) Time-traces of mean receiver activity per shell including the standard deviation (shaded region). All conditions are summarized in Supplementary Table 2, entry 5. Scale bar 100 µm.

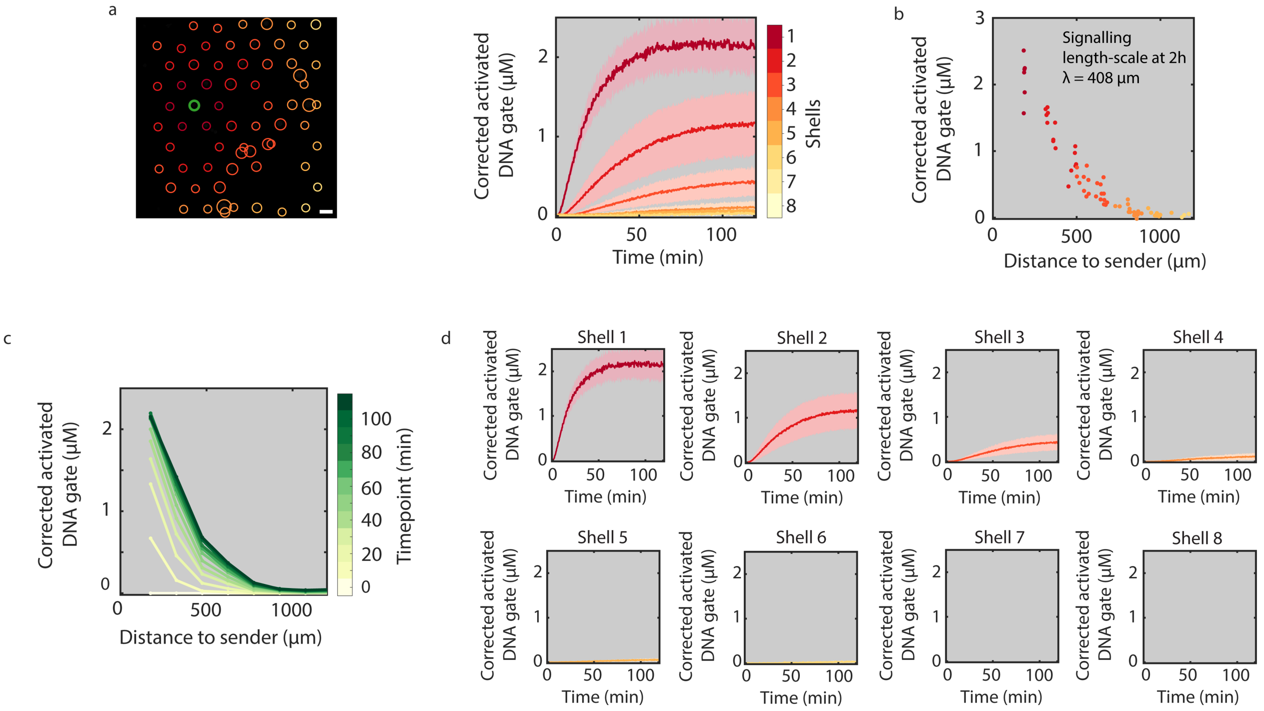

**Supplementary Figure 23.** Activation of receiver protocells with low protocellular density (data set 3/3). (a) Time-traces of receiver activation including the standard deviation (shaded regions). (b) Concentrations of activated DNA gates in individual receivers after 2 h of signal release as function of the distance to the sender protocell and the calculated signaling length-scale. (c) The concentration of activated gate complexes in receiver protocells is plotted for different times of the reaction as a function of distance to the sender. (d) Time-traces of mean receiver activity per shell including the standard deviation (shaded region). All conditions are summarized in Supplementary Table 2, entry 5. Scale bar 100 μm.

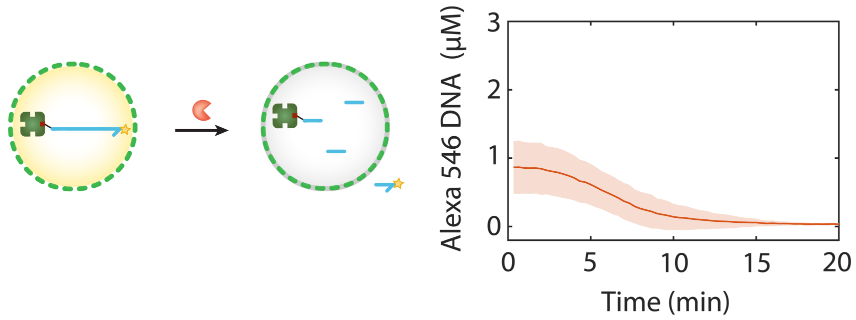

**Supplementary Figure 24.** Diffusion of exonuclease into protocells. Exonuclease I (0.1 Unit/μL) was added to the buffer outside protocells containing Alexa 546-labeled ssDNA (3’ to 5’). Subsequently, the concentration of the fluorescently-labeled DNA was monitored over time (mean and standard deviation are shown as line and shaded area respectively. The graph shows that exonuclease can enter the protocells and degrade the DNA.

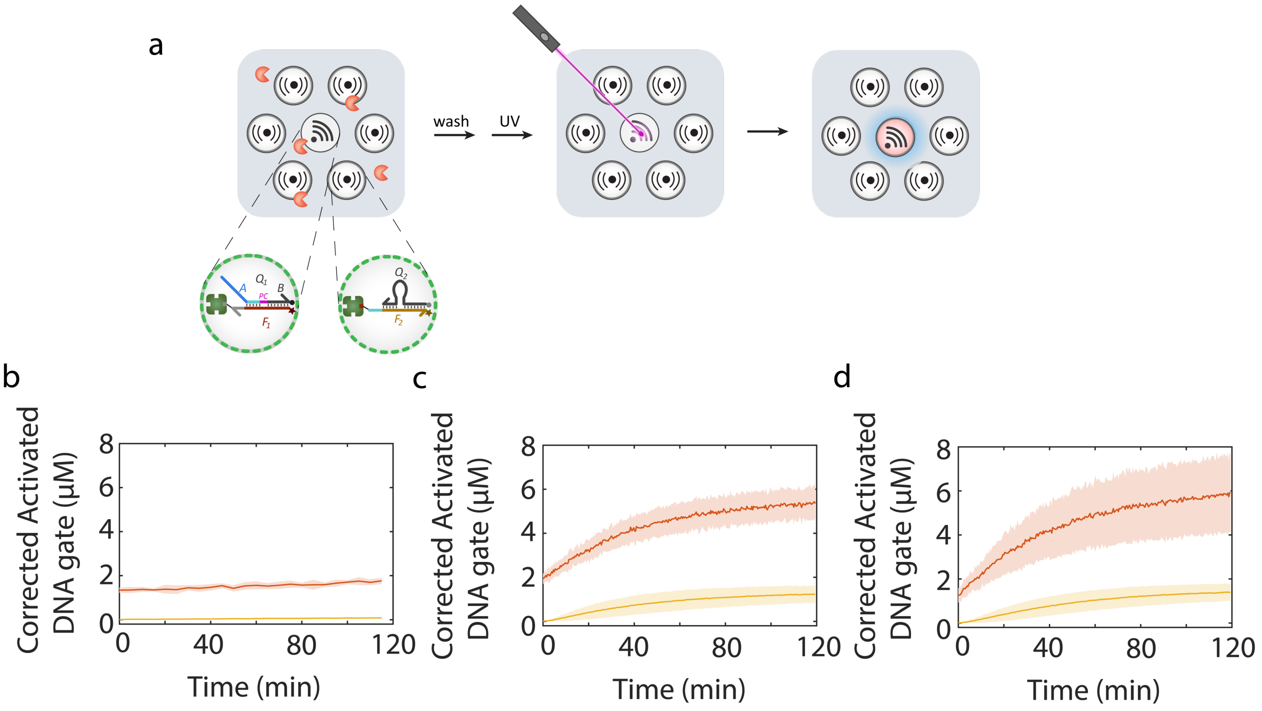

**Supplementary Figure 25.** The effect of exonuclease on 5' to 3' ssDNA and dsDNA stability in protocells. (a) Light-triggered signal release and receiver activation after incubation in the presence of exonuclease. There is a free 5'-end on one side of the sender DNA gate complex, while all other terminal bases on sender and receiver DNA gates are hybridized. (b) Time-traces of sender (red line) and receiver (yellow line) activated DNA gates when incubated in the presence of 0.05 Unit/μL exonuclease I for 2 hours. The exonuclease was then removed by gently flowing in buffer through the trapping chamber. (c) Signal release from the sender upon UV irradiation and subsequent receiver activation is shown. (d) Control experiment in absence of exonuclease I. The amount of streptavidin used to prepare sender and receiver protocells are 2 μM and 0.2 μM respectively. The fractions of the two populations were sender = 0.1, receiver = 0.9.

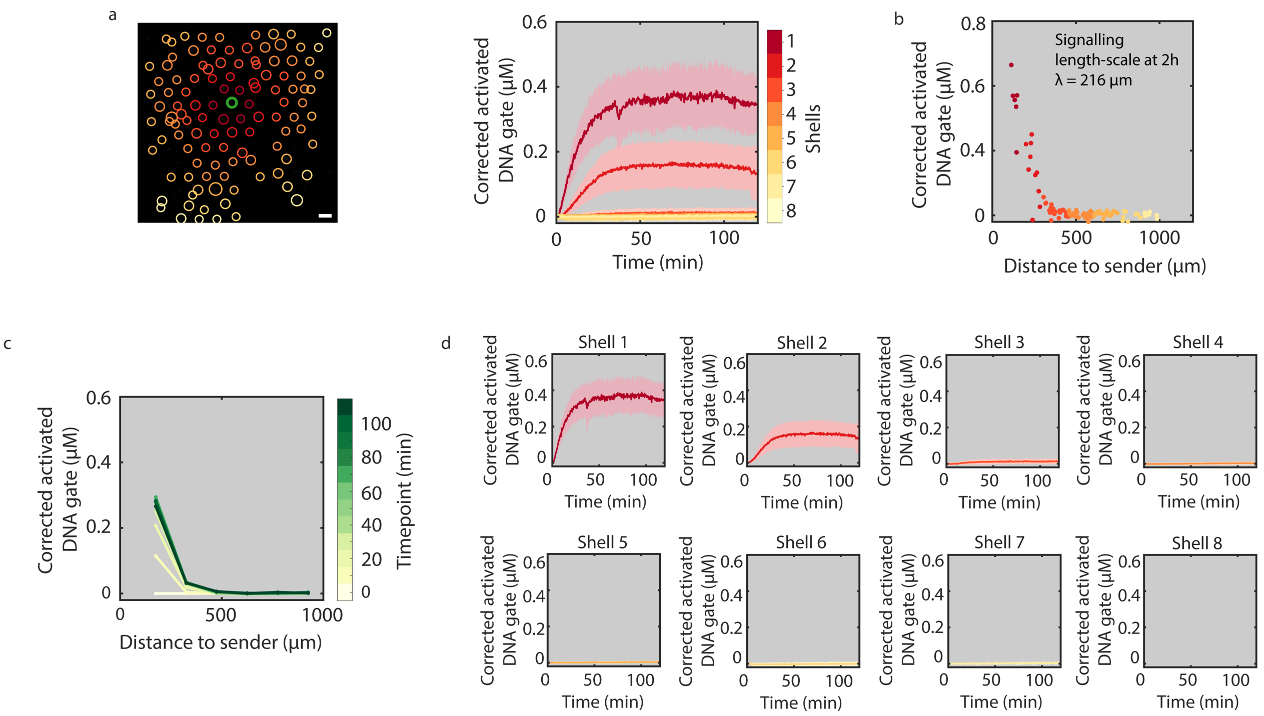

**Supplementary Figure 26.** Activation of receiver protocells in the presence of high concentration (0.1 Unit/µL) of exonuclease (data set 1/3).(a) Time-traces of receiver activation including the standard deviation (shaded regions). (b) Concentrations of activated DNA gates in individual receivers after 2 h of signal release as function of the distance to the sender protocell and the calculated signaling length-scale. (c) The concentration of activated gate complexes in receiver protocells is plotted for different times of the reaction as a function of distance to the sender. (d) Time-traces of mean receiver activity per shell including the standard deviation (shaded region). All conditions are summarized in Supplementary Table 2, entry 6. Scale bar 100 µm.

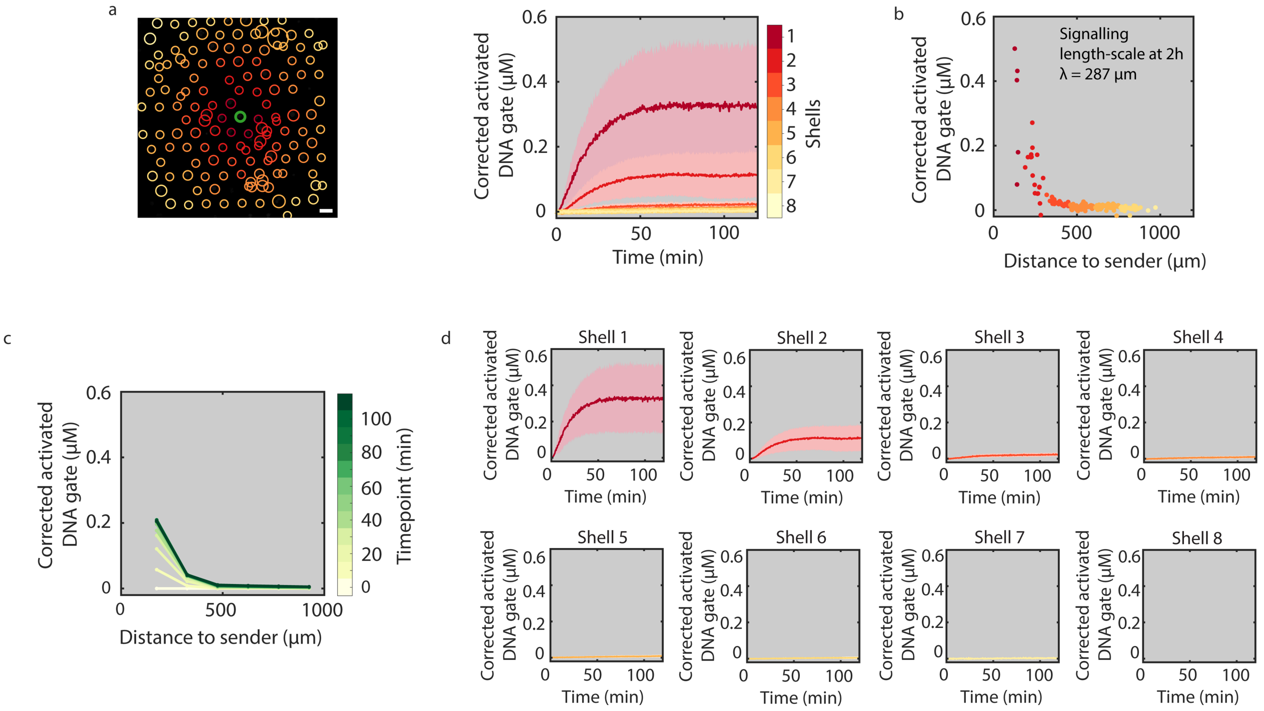

**Supplementary Figure 27.** Activation of receiver protocells corresponding to a high concentration (0.1 Unit/µL) of exonuclease (data set 2/3). (a) Time-traces of receiver activation including the standard deviation for each shell (shaded regions). (b) Concentrations of activated DNA gates in individual receivers after 2 h of signal release as function of the distance to the sender protocell and the calculated signaling length-scale. (c) The concentration of activated gate complexes in receiver protocells is plotted for different times of the reaction as a function of distance to the sender. (d) Time-traces of mean receiver activity per shell including the standard deviation (shaded region). All conditions are summarized in Supplementary Table 2, entry 6. Scale bar 100 µm.

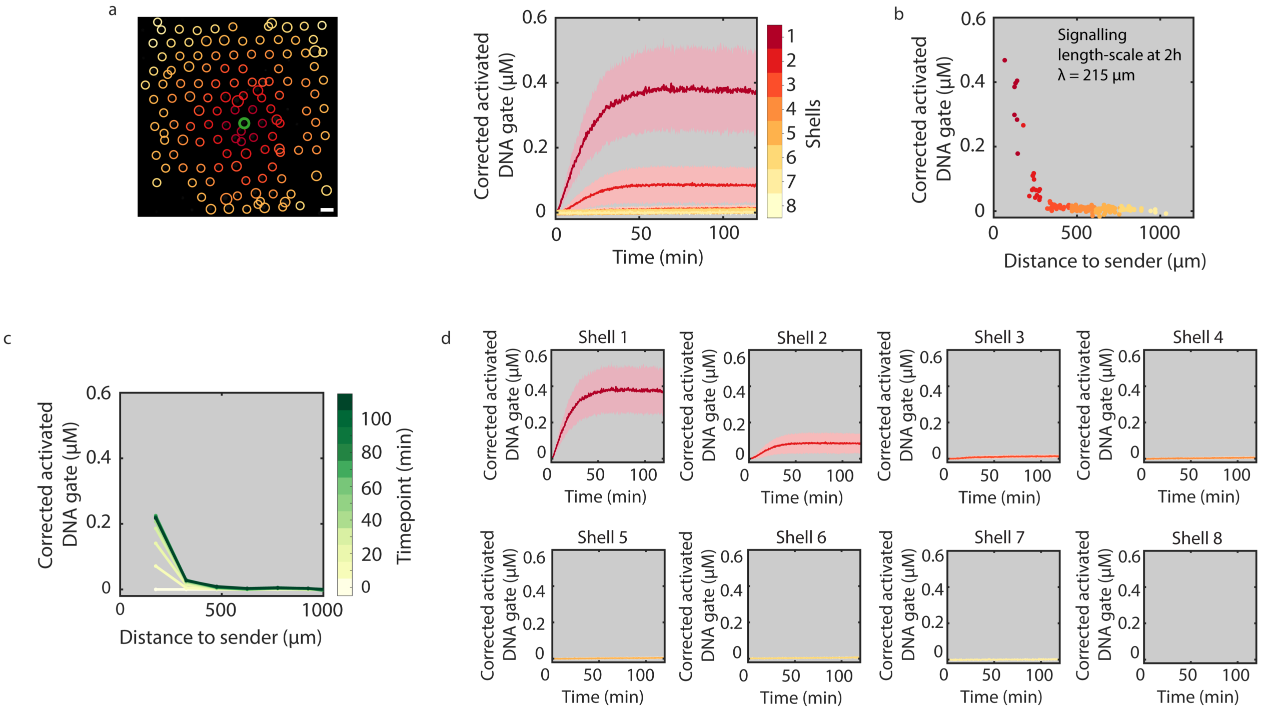

**Supplementary Figure 28.** Activation of receiver protocells corresponding to a high concentration (0.1 Unit/µL) of exonuclease (data set 3/3). (a) Time-traces of receiver activation including the standard deviation for each shell (shaded regions). (b) Concentrations of activated DNA gates in individual receivers after 2 h of signal release as function of the distance to the sender protocell and the calculated signaling length-scale. (c) The concentration of activated gate complexes in receiver protocells is plotted for different times of the reaction as a function of distance to the sender. (d) Time-traces of mean receiver activity per shell including the standard deviation (shaded region). All conditions are summarized in Supplementary Table 2, entry 6. Scale bar 100 µm.

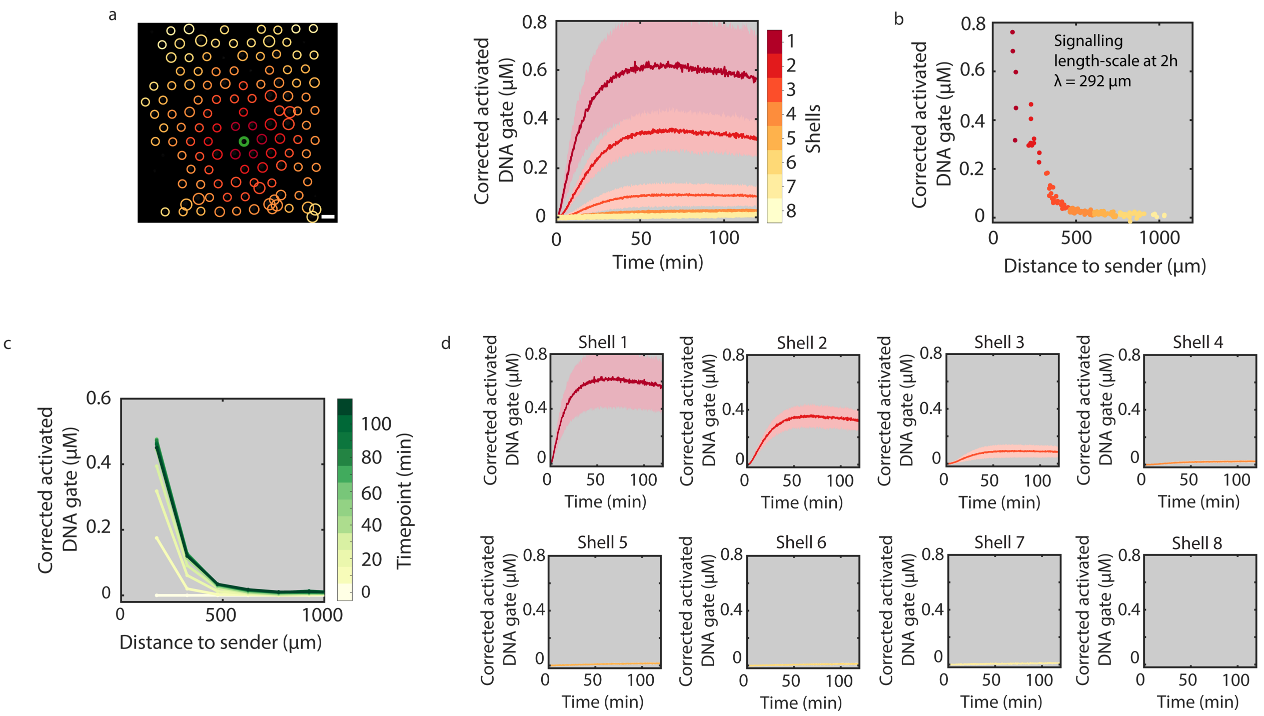

**Supplementary Figure 29.** Activation of receiver protocells corresponding to a low concentration (0.05 Unit/μL) of exonuclease (data set 1/3). (a) Time-traces of receiver activation including the standard deviation for each shell (shaded regions). (b) Concentrations of activated DNA gates in individual receivers after 2 h of signal release as function of the distance to the sender protocell and the calculated signaling length-scale. (c) The concentration of activated gate complexes in receiver protocells is plotted for different times of the reaction as a function of distance to the sender. (d) Time-traces of mean receiver activity per shell including the standard deviation (shaded region). All conditions are summarized in Supplementary Table 2, entry 7. Scale bar 100 μm.

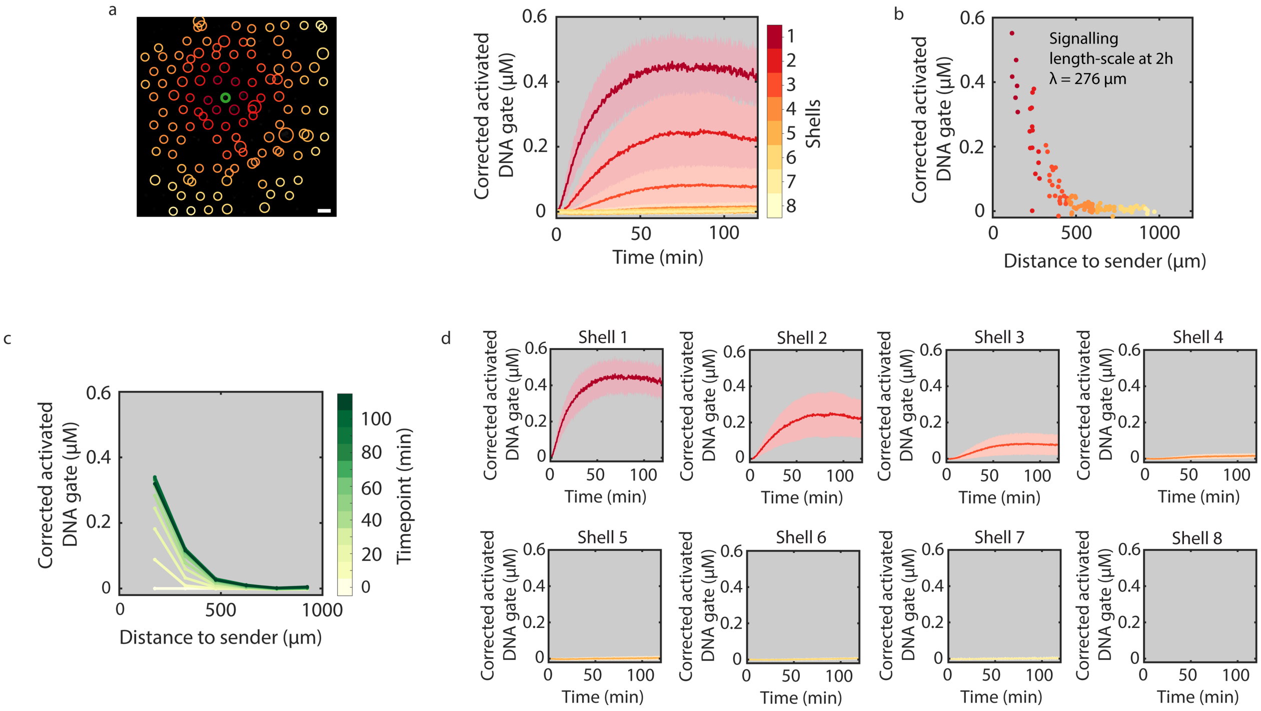

**Supplementary Figure 30.** Activation of receiver protocells corresponding to the low concentration (0.05 Unit/µL) of exonuclease (data set 2/3). (a) Time-traces of receiver activation including the standard deviation for each shell (shaded regions). (b) Concentrations of activated DNA gates in individual receivers after 2 h of signal release as function of the distance to the sender protocell and the calculated signaling length-scale. (c) The concentration of activated gate complexes in receiver protocells is plotted for different times of the reaction as a function of distance to the sender. (d) Time-traces of mean receiver activity per shell including the standard deviation (shaded region). All conditions are summarized in Supplementary Table 2, entry 7. Scale bar 100 µm.

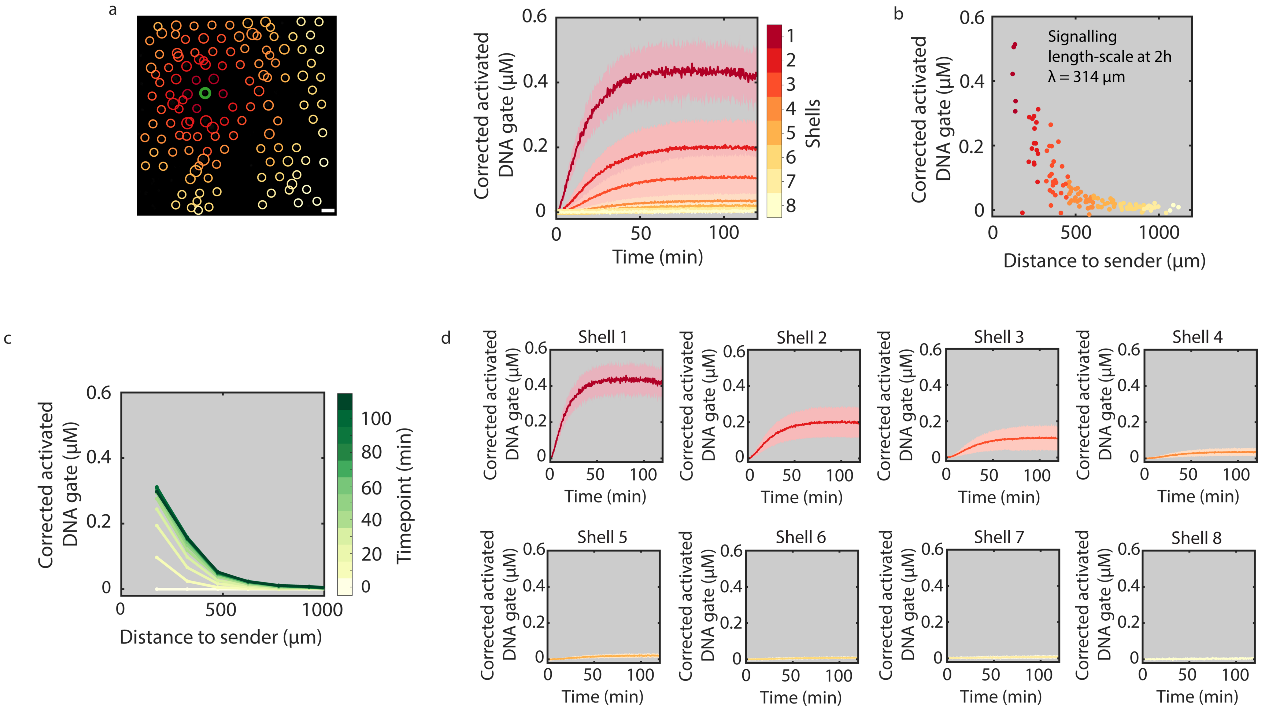

**Supplementary Figure 31.** Activation of receiver protocells corresponding to a low concentration (0.05 Unit/µL) of exonuclease (data set 3/3). (a) Time-traces of receiver activation including the standard deviation (shaded regions). (b) Concentrations of activated DNA gates in individual receivers after 2 h of signal release as function of the distance to the sender protocell and the calculated signaling length-scale. (c) The concentration of activated gate complexes in receiver protocells is plotted for different times of the reaction as a function of distance to the sender. (d) Time-traces of mean receiver activity per shell including the standard deviation (shaded region). All conditions are summarized in Supplementary Table 2, entry 7. Scale bar 100 µm.

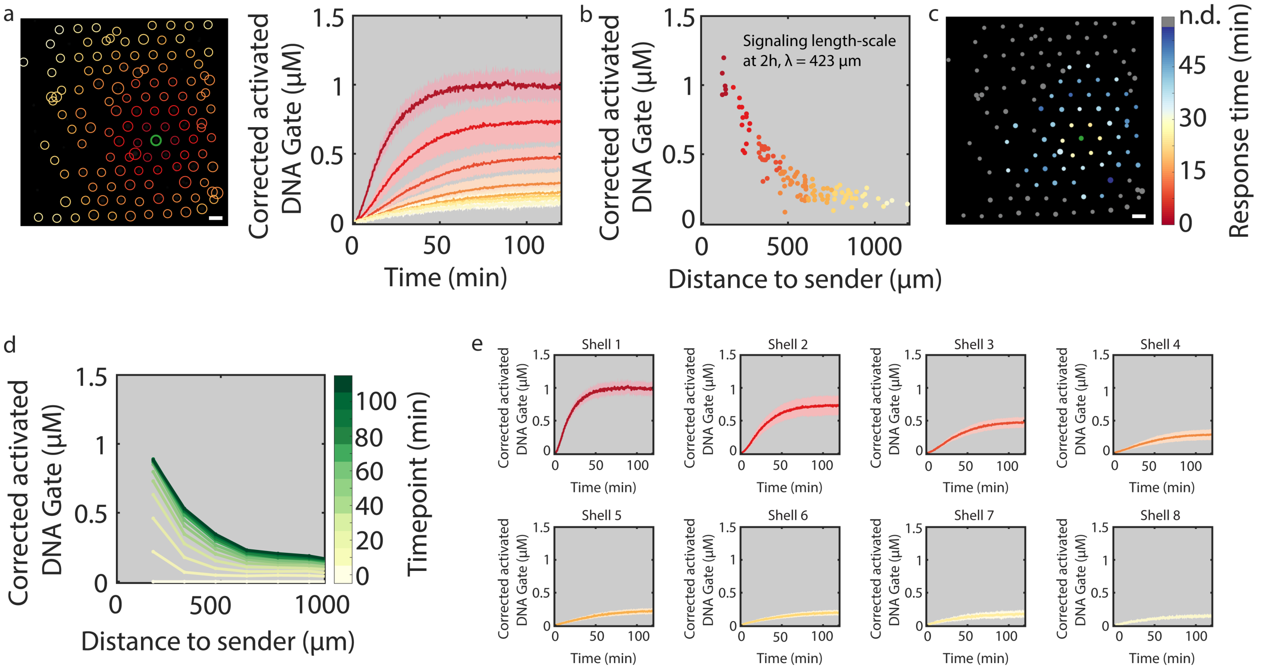

**Supplementary Figure 32.** Activation of transceiver protocells in the presence of 0.1 µM fuel (data set 1/3). (a) Time-traces of receiver activation including the standard deviation for each shell (shaded regions). (b) Concentrations of activated DNA gates in individual receivers after 2 h of signal release as function of the distance to the sender protocell and the calculated signaling length-scale. (c) Spatial barcode image of response time of transceiver protocells. The response time is defined as the time in which a transceiver reaches 50% of its final activated concentration. (red: short 'on' time; blue: long 'on' time). To remove background noise, any protocells with an absolute increase less than 20 RFU are excluded and labelled with gray (n.d.). The sender protocell is labeled with green. (d) The concentration of activated gate complexes in receiver protocells is plotted for different times of the reaction as a function of distance to the sender. (e) Time-traces of mean receiver activity per shell including the standard deviation (shaded region). All conditions are summarized in Supplementary Table 2, entry 8. Scale bar 100 µm.

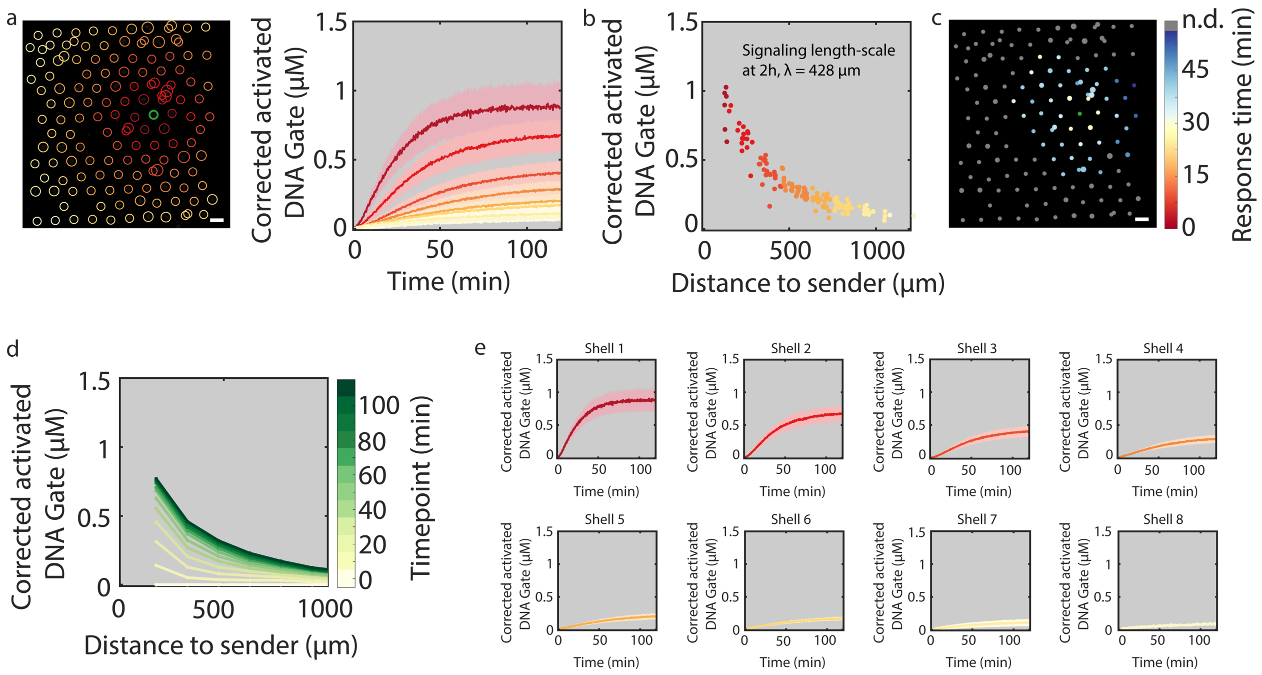

**Supplementary Figure 33.** Activation of transceiver protocells in the presence of 0.1 µM fuel (data set 2/3). (a) Time-traces of receiver activation including the standard deviation for each shell (shaded regions). (b) Concentrations of activated DNA gates in individual receivers after 2 h of signal release as function of the distance to the sender protocell and the calculated signaling length-scale. (c) Spatial barcode image of response time of transceiver protocells. The response time is defined as the time in which a transceiver reaches 50% of its final activated concentration. (red: short 'on' time; blue: long 'on' time). To remove background noise, any protocells with an absolute increase less than 20 RFU are excluded and labelled with gray (n.d.). The sender protocell is labeled with green. (d) The concentration of activated gate complexes in receiver protocells is plotted for different times of the reaction as a function of distance to the sender. (e) Time-traces of mean receiver activity per shell including the standard deviation (shaded region). All conditions are summarized in Supplementary Table 2, entry 8. Scale bar 100 µm.

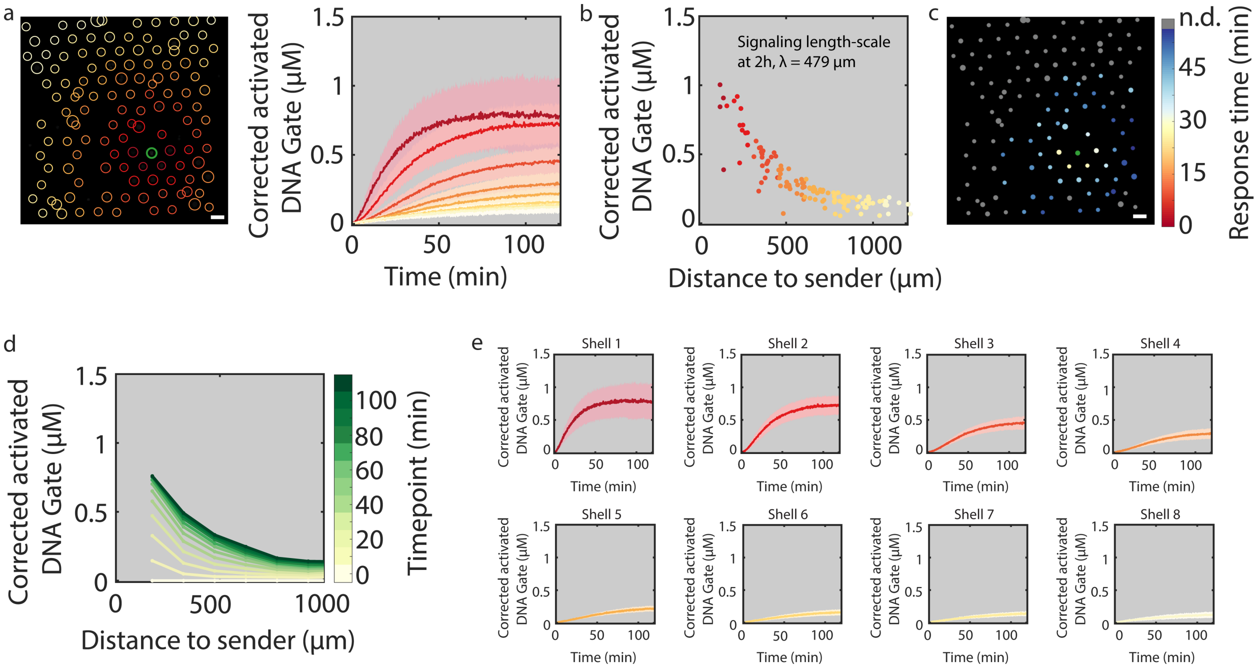

**Supplementary Figure 34.** Activation of transceiver protocells in the presence of 0.1 μM fuel (data set 3/3). (a) Time-traces of receiver activation including the standard deviation for each shell (the highlighted regions). (b) Concentrations of activated DNA gates in individual receivers after 2 h of signal release as function of the distance to the sender protocell and the calculated signaling length-scale. (c) Spatial barcode image of response time of transceiver protocells. The response time is defined as the time in which a transceiver reaches 50% of its final activated concentration. (red: short 'on' time; blue: long 'on' time). To remove background noise, any protocells with an absolute increase less than 20 RFU are excluded and labelled with gray (n.d.). The sender protocell is labeled with green. (d) The concentration of activated gate complexes in receiver protocells is plotted for different times of the reaction as a function of distance to the sender. (e) Time-traces of mean receiver activity per shell including the standard deviation (shaded region). All conditions are summarized in Supplementary Table 2, entry 8. Scale bar 100 μm.

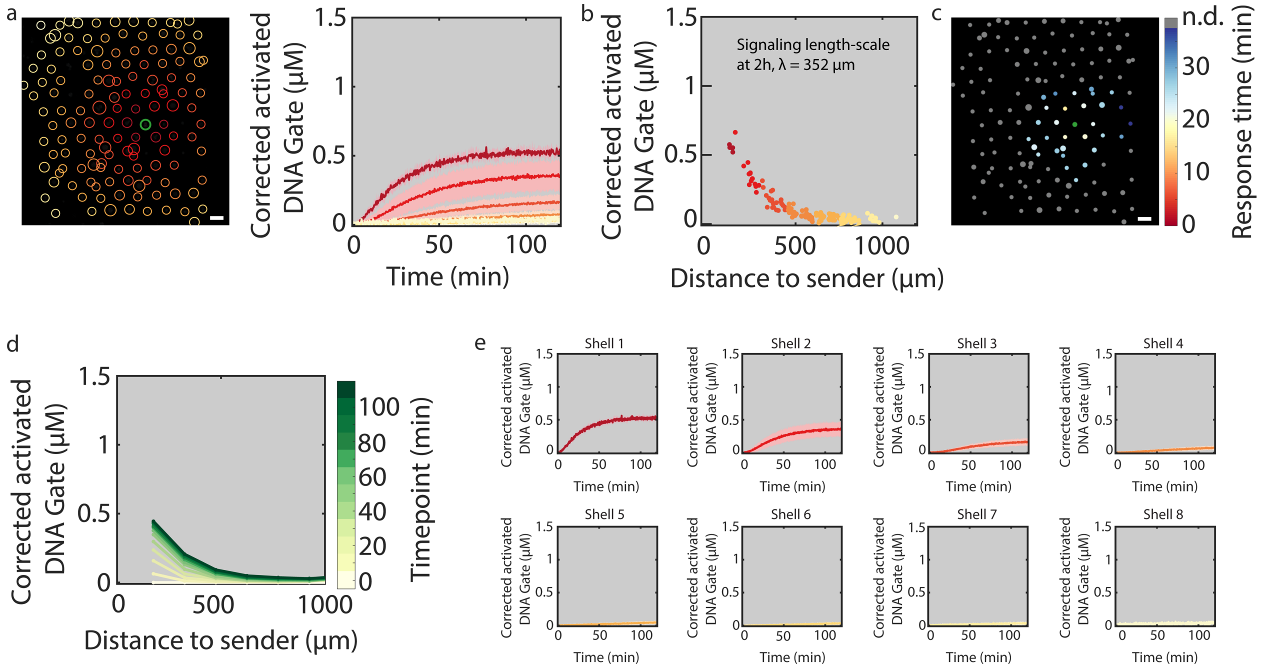

**Supplementary Figure 35.** Activation of transceiver protocells in the absence of fuel (data set 1/3). (a) Time-traces of receiver activation including the standard deviation for each shell (shaded regions). (b) Concentrations of activated DNA gates in individual receivers after 2 h of signal release as function of the distance to the sender protocell and the calculated signaling length-scale. (c) Spatial barcode image of response time of transceiver protocells. The response time is defined as the time in which a transceiver reaches 50% of its final activated concentration. (red: short ‘on’ time; blue: long ‘on’ time). To remove background noise, any protocells with an absolute increase less than 20 RFU are excluded and labeled with gray (n.d.). The sender protocell is labeled with green. (d) The concentration of activated gate complexes in receiver protocells is plotted for different times of the reaction as a function of distance to the sender. (e) Time-traces of mean receiver activity per shell including the standard deviation (shaded region). All conditions are summarized in Supplementary Table 2, entry 8. Scale bar 100 μm.

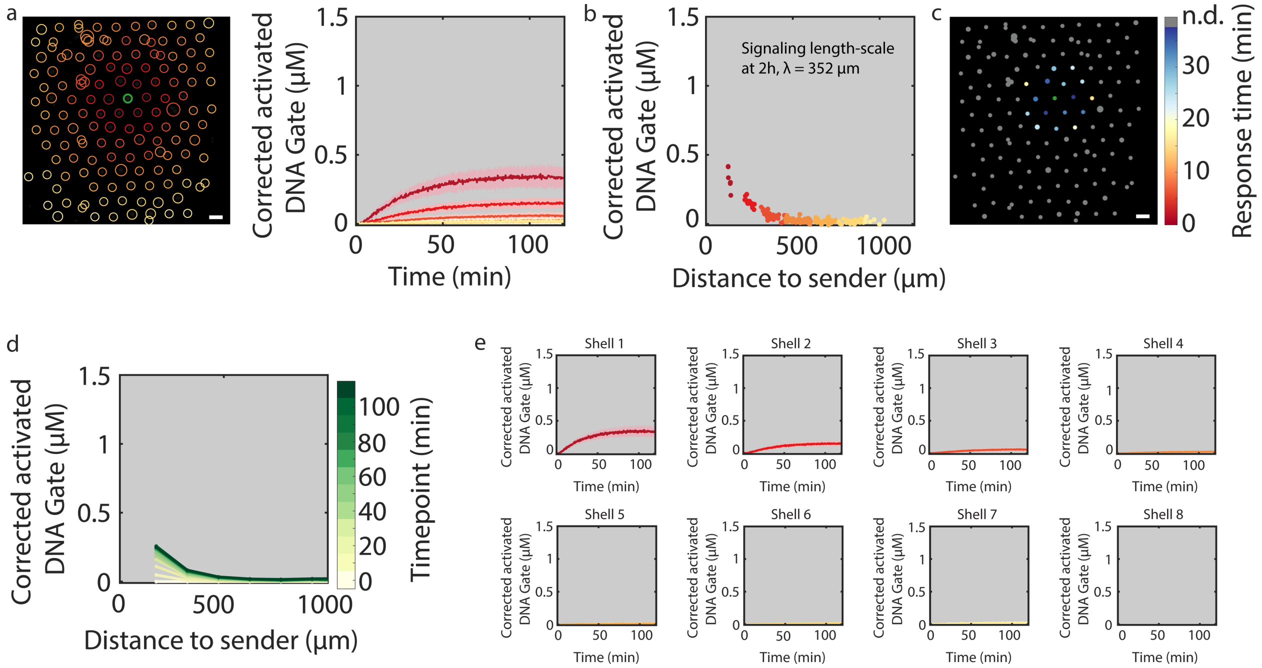

**Supplementary Figure 36.** Activation of transceiver protocells in the absence of fuel (data set 2/3). (a) Time-traces of receiver activation including the standard deviation for each shell (shaded regions). (b) Concentrations of activated DNA gates in individual receivers after 2 h of signal release as function of the distance to the sender protocell and the calculated signaling length-scale. (c) Spatial barcode image of response time of transceiver protocells. The response time is defined as the time in which a transceiver reaches 50% of its final activated concentration. (red: short ‘on’ time; blue: long ‘on’ time). To remove background noise, any protocells with an absolute increase less than 20 RFU are excluded and labeled with gray (n.d.). The sender protocell is labeled with green. (d) The concentration of activated gate complexes in receiver protocells is plotted for different times of the reaction as a function of distance to the sender. (e) Time-traces of mean receiver activity per shell including the standard deviation (shaded region). All conditions are summarized in Supplementary Table 2, entry 8. Scale bar 100 µm.

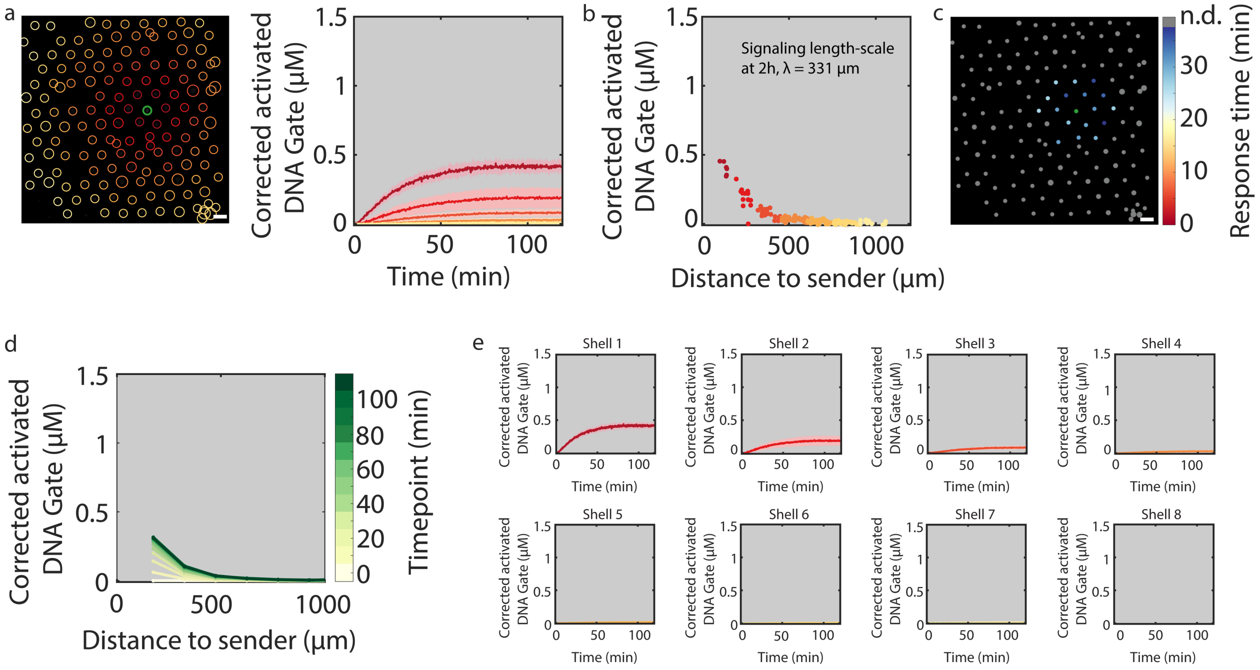

**Supplementary Figure 37.** Activation of transceiver protocells in the absence of fuel (dataset 3/3). (a) Time-traces of receiver activation including the standard deviation for each shell (shaded regions). (b) Concentrations of activated DNA gates in individual receivers after 2 h of signal release as function of the distance to the sender protocell and the calculated signaling length-scale. (c) Spatial barcode image of response time of transceiver protocells. The response time is defined as the time in which a transceiver reaches 50% of its final activated concentration. (red: short ‘on’ time; blue: long ‘on’ time). To remove background noise, any protocells with an absolute increase less than 20 RFU are excluded and labelled with gray (n.d.). The sender protocell is labeled with green. (d) The concentration of activated gate complexes in receiver protocells is plotted for different times of the reaction as a function of distance to the sender. (e) Time-traces of mean receiver activity per shell including the standard deviation (shaded region). All conditions are summarized in Supplementary Table 2, entry 8. Scale bar 100 μm.

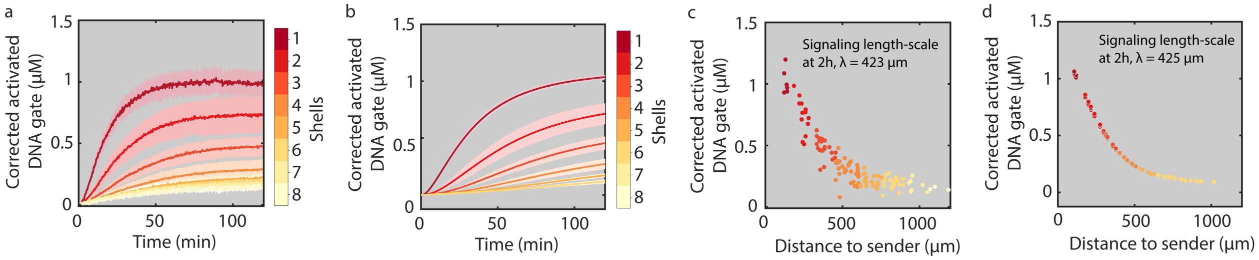

**Supplementary Figure 38.** Experimental and simulated data from 2D reaction-diffusion simulations corresponding to the sender-transceiver configuration. Experimental (a) and simulated (b) time traces of activated transceiver protocells corresponding to different shells including standard deviation. The standard deviation per shell in the model originates from the different distance of individual transceiver protocells toward sender protocell. Experimental (c) and simulated (d) concentrations of activated DNA gates in individual transceivers after 2 h of signal release as function of the distance to the sender protocell and the calculated signaling length-scales. The reactions and parameters used in the model are shown in Supplementary Methods and Table 5.

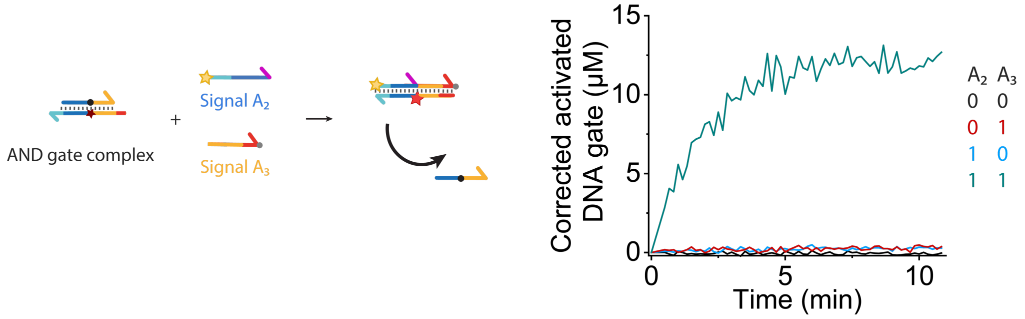

**Supplementary Figure 39.** Time traces of batch experiments with the AND logic circuit based on cooperative strand displacement[3]. The final concentrations of AND gate complex, signal $\mathbf{A_2}$ and signal $\mathbf{A_3}$ were 40 nM, 20 nM and 20 nM respectively. The color-coded traces correspond to the rows of the table on the right which indicates the specific signal configuration (presence (1) or absence (0) of the two input signals $\mathbf{A_2}$ and $\mathbf{A_3}$).

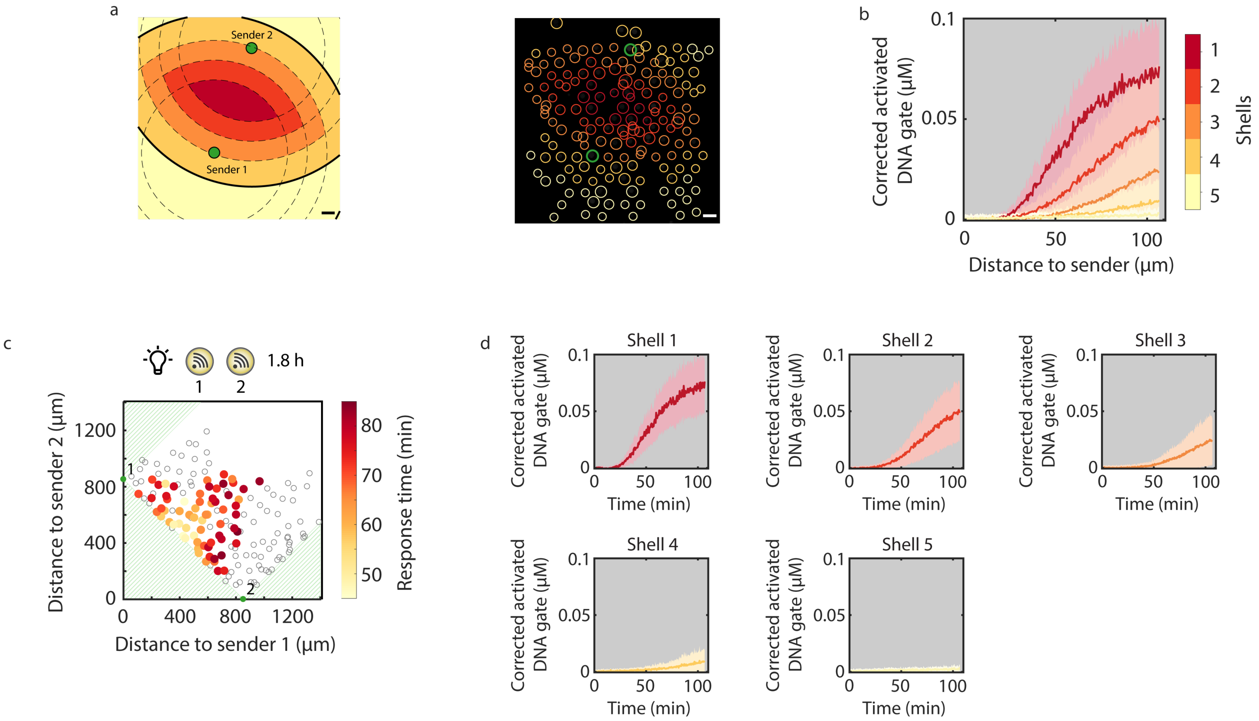

**Supplementary Figure 40.** Spatiotemporal activation of AND-gate receiver protocells upon simultaneous laser irradiation (405 nm, 1.5 h). (a) Illustration of the binning method to analyze spatiotemporal activation of AND-gate receiver protocells. Protocells are binned based on the maximum of the two distances to the senders, which yields bins with outer bounds that are the intersection of the equivalent bounds of single sender systems, as illustrated by the black lines. (b) Time-traces of AND-gate receiver activation including the standard deviation (shaded region). (c) Response time of individual receivers upon simultaneous laser irradiation of both senders for 1.8 h is plotted as a function of their distance to each sender. The response time is defined as the time it takes for an individual receiver to reach 50% of its final activated concentration. Two senders are marked in green. To remove background noise, any protocells with an absolute increase less than 1 RFU are excluded and labelled in hollow circles. (d) Time-traces of mean AND-gate receiver activity per shell including the standard deviation (shaded region). All conditions are summarized in Supplementary Table 2, entry 9. Scale bar 100 µm.

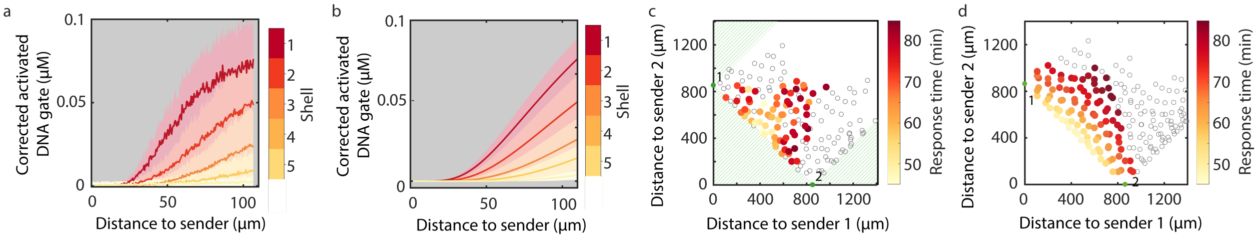

**Supplementary Figure 41.** Experimental and simulated response of the AND-gate system obtained by 2D reaction-diffusion simulations. The experimental (a) and simulated (b) time traces of activated receiver protocells in different shells including standard deviation. The standard deviation per shell in the model originates from the different distance of individual receiver protocells toward sender protocell. Experimental (c) and simulated (d) response time of activated DNA gates in individual receivers after 1.8 h of signal release as function of their distance to each sender protocell. The response time is defined as the time it takes for an individual receiver to reach 50% of its final activated concentration. To exclude the effect of background reactions in the experimental data, any protocells with an absolute increase less than 1 RFU are excluded and labeled as hollow circles. Similarly, a threshold is set in the simulation with a concentration of 0.02 μM, which is converted from the experimental RFU threshold. Two senders are marked in green. The reactions and parameters used in the model are shown in Supplementary Methods and Table. 6.

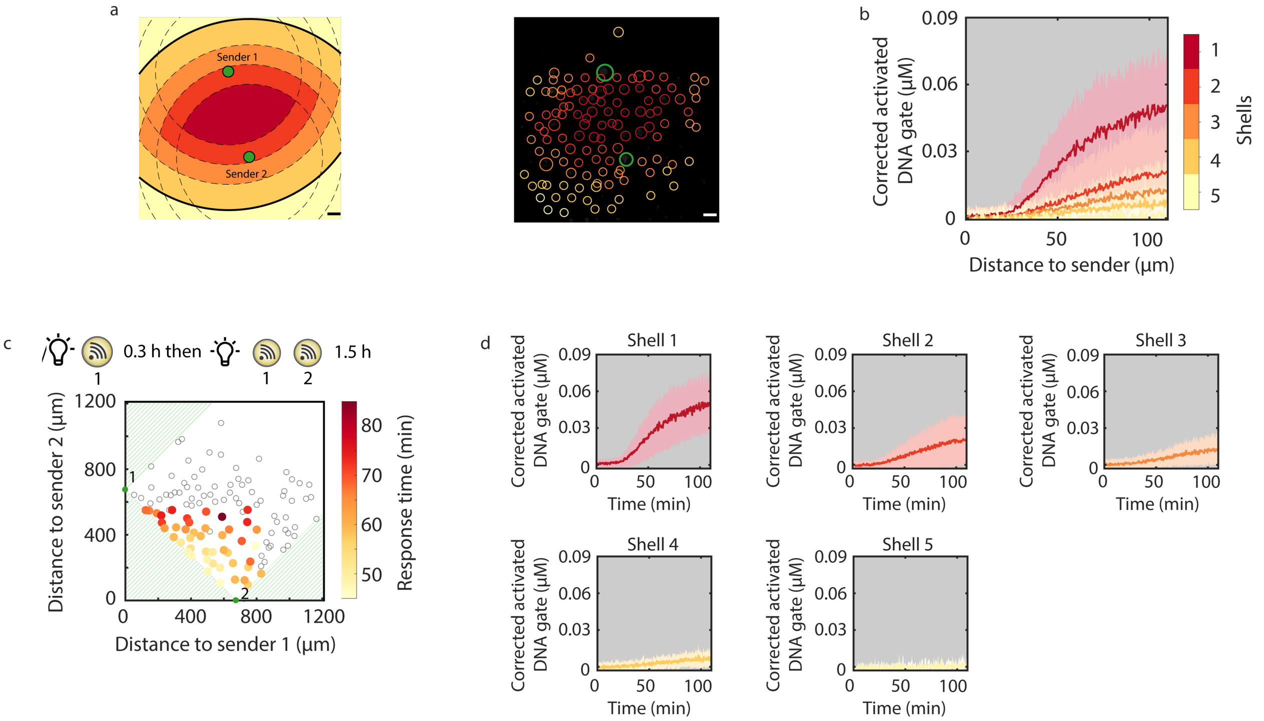

**Supplementary Figure 42.** Spatiotemporal activation of AND-gate receiver protocells upon sequential laser (405 nm) irradiation (Sender **1** is irradiated for 18 min, followed by irradiation of both senders for 1.5 h). (a) Illustration of the binning method to analyze spatiotemporal activation of AND-gate receiver protocells. Protocells are binned based on the maximum of the two distances to the senders, which yields bins with outer bounds that are the intersection of the equivalent bounds of single sender systems, as illustrated by the black lines. (b) Time-traces of AND-gate receiver activation including the standard deviation (shaded regions). (c) Response time of individual receivers upon sequential laser irradiation is plotted as a function of their distance to each sender. The response time is defined as the time it takes for an individual receiver to reach 50% of its final activated concentration. Two senders are marked in green. To remove background noise, any protocells with an absolute increase less than 1 RFU are excluded and labelled in hollow circles. (d) Time-traces of mean AND-gate receiver activity per shell including the standard deviation (shaded region). All conditions are summarized in Supplementary Table 2, entry 9. Scale bar 100 µm.

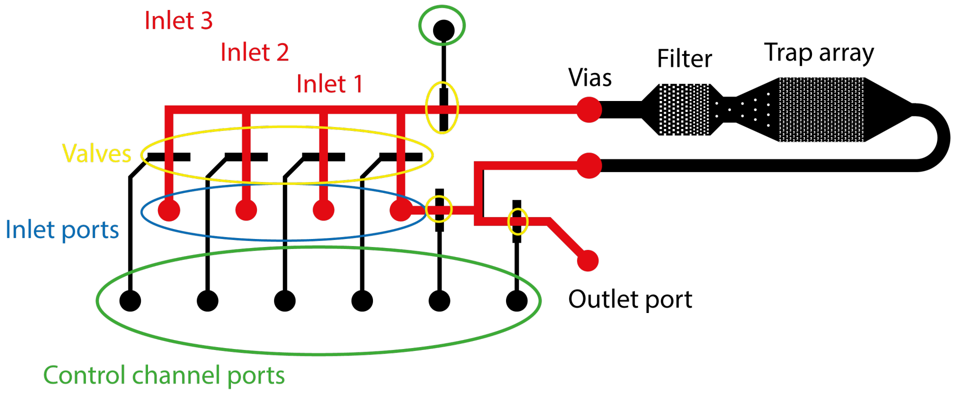

**Supplementary Figure 43.** Design of the microfluidic device[5]. CAD drawing of the microfluidic device. Channels of the bottom layer (bonded to the glass slide) are in black. The rounded top layer channels are in red. The flow channels cross from the top layer to the bottom layer as only the rounded channels of the top layer can be closed using the pneumatic valves, while the trapping array requires the rectangular channels of the bottom layer.

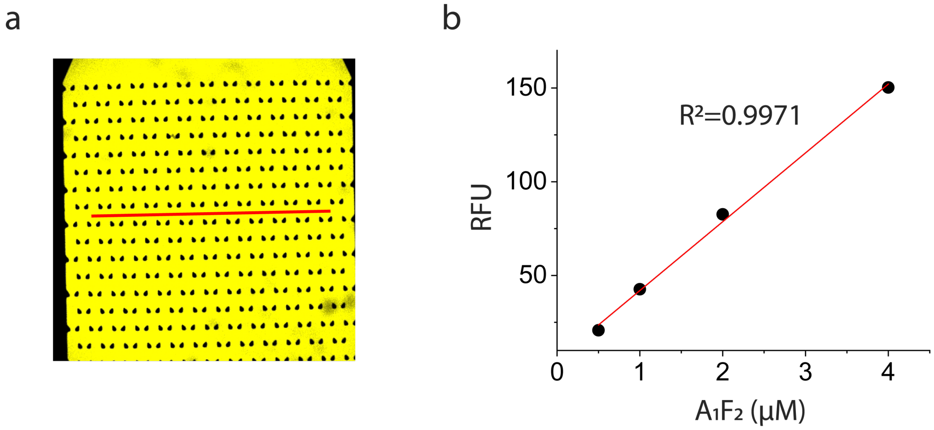

**Supplementary Figure 44.** Verification of the linearity of the concentration estimation method. (a) A confocal micrograph of the microfluidic trapping device filled with active form of the gate complex **$F_2A_1$** (4 μM). The corresponding RFU was obtained by measuring the average RFU value across a straight horizontal line (red) through the device. (b) The linearity of the conversion method was verified by performing the measurement shown in (a) over a range of concentrations (500 nM, 1 μM, 2 μM and 4 μM) of **$F_2A_1$**, fitting a straight line through the data and calculating the coefficient of determination ($R^2$).

**Supplementary Table 1.** Calculated free energies of the DNA gate complex used in sender protocell.

| Gate complex | Calculated ΔG° (kcal/mol) |
|---|---|
| Before laser irradiation | |
| **$F_1$-$Q_1$** | -21.43 |
| After laser irradiation | |
| **$F_1$-A** | -12.94 |
| **$F_1$-B** | -11.49 |

The free energies were predicted by the Nucleid Acid Package (NUPACK)[4]. The buffer conditions were set to 0.05 M $Na^+$ and 0.115 M $Mg^{2+}$ and a temperature of 25°C was used.

**Supplementary Table 2.** Summarization of experimental conditions.

| Entry | Main text Fig number | Supple-mentary Fig number | Streptavidin Sender (μM) | Binding capacity (μM) | Streptavidin receiver (μM) | Consumption capacity (μM) | Receiver permeability* | Receiver density† | Exonuclease‡ |
|---|---|---|---|---|---|---|---|---|---|
| Sender-receiver system | | | | | | | | | |
| 1 | 1<br>2b (high)<br>2d (high) | 4, 7-8 | 10 | 19.3 | 4 | 2 | high | high | no |
| 2 | 2b(medium)<br>2c (high)<br>2e (control) | 9-11 | 10 | 19.3 | 2 | 0.82 | high | high | no |
| 3 | 2b (low) | 12-14 | 10 | 19.3 | 1 | 0.25 | high | high | no |
| 4 | 2c (low) | 16-18 | 10 | 19.3 | 4 | 0.82 | low | high | no |
| 5 | 2d (low) | 20-22 | 10 | 19.3 | 4 | 2 | high | low | no |
| 6 | 2e (high) | 25-27 | 10 | 19.3 | 2 | 0.82 | high | high | high |
| 7 | 2e (low) | 28-30 | 10 | 19.3 | 2 | 0.82 | high | high | low |
| Sender-transceiver system | | | | | | | | | |
| 8 | 3 | 31-36 | 10 | 10 | 2 | 1 | high | high | no |
| Boolean AND gate | | | | | | | | | |
| 9 | 4 | 39, 41 | 30 | 40 | 2 | 0.82 | high | high | no |

*.Receiver permeability: High 202.8 μm $min^{-1}$ and low 2.16 μm $min^{-1}$

†.Receiver density: High 90 traps per $mm^2$ and low 70 traps per $mm^2$

‡.Concentration of Exonuclease: High 0.1 unit/μL and low 0.05 unit/μL

**Supplementary Table 3**. Summarization of DNA sequences.

| sender | | Length bases | 5' modification | 3' modification |
|---|---|---|---|---|
| Fig. 1, 2 and 3, Supplementary Figure 3, 4, 5, 8-15, 17-19, 21-23, 26-37 | | | | |
| $Q_1$ | CGA ACG AAC GAC CAT GAT AGA CTA ATG CAC TAC TAC /PC/ TAA CTA GC | 44 | | Iowa Black |
| $F_1$ | GCT AGT TAT TGT AGT AGT TTT TTT TT | 26 | Cy5 | Biotin-TEG |
| Supplementary Figure 2 | | | | |
| $Q_7$ | CGA ACG AAC GAC CAT GAT AGA CTA ATG CAC TAC TAC /PC/ TAA CTA GC | 44 | Cy3 | Iowa Black |
| $F_1$ | GCT AGT TAT TGT AGT AGT TTT TTT TT | 26 | Cy5 | Biotin-TEG |
| Fig. 4 and Supplementary Figure 40, 42 | | | | |
| $Q_4$ | CAC ACA TCT ATA CAA CCA CTT ACT T /PC/ TA ACT ACC | 33 | Cy3 | |
| $F_4$ | GGT AGT TAT TAA GTA AGT TTT TTT TT | 26 | | Biotin-TEG |
| $Q_5$ | TAA CTA CC /PC/ G CCA TCA GAA CTT AAC CTA ACT CCT | 33 | Cy3 | Iowa Black |
| $F_5$ | TGA TGG CTT GGT AGT TAT TTT TTT T | 25 | | Biotin-TEG |
| Supplementary Figure 39 | | | | |
| $A_2$ | CAC ACA TCT ATA CAA CCA CTT ACT T | 25 | | |
| $A_3$ | GCC ATC AGA ACT TAA CCT AAC TCC T | 25 | | |
| receiver | | Length bases | 5' modification | 3' modification |
| Fig. 1, 2 and Supplementary Figure 2, 5, 8-15, 17-19, 21-23, 26-31 | | | | |
| $Q_2$ | CGA ACG AAC GAC CAT GCG TGA AAC ATA GAC TAA TGC | 36 | Iowa Black | phosphate |
| $F_2$ | GTA GTA GTG CAT TAG TCT ATC ATG GTC GTT CGT TCG | 36 | Biotin-TEG | Alexa546 |
| Fig. 3 and Supplementary Figure 32-37 | | | | |
| $Q_3$ | GGA ACT GTC GAA CGA ACG TGA AAC CGA CCA TGA TAG | 36 | Iowa Black | phosphate |
| $F_3$ | GCA TTA GTC TAT CAT GGT CGT TCG TTC GAC AGT TCC | 36 | Biotin-TEG | Alexa546 |
| Fuel | GGA ACT GTC GAA CGA ACG ACC ATG ATA G | 28 | | phosphate |
| Fig. 4 and Supplementary Figure 39, 40, 42 | | | | |
| $Q_6$ | CTA TAC AAC CAC TTA CTT /BHQ/ GC CAT CAG AAC TTA ACC | 35 | | |
| $F_6$ | AGG AGT TAG GTT AAG TTC TGA TGG C /Quasar670/ AAG TAA GTG GTT GTA TAG ATG TGT GTT TTT TTT | 58 | | Biotin-TEG |

**Supplementary Table 4**. Simulation parameters for sender-receiver system.

| | |
|---|---|
| Release rate ($k_{rel}$) | $2.7 \times 10^{-2}$ $min^{-1}$ |
| Permeability (P) | 200 μm $min^{-1}$ |
| Diffusion coefficient of DNA in buffer (d) | $2.2 \times 10^{3}$ $μm^{2}$ $min^{-1}$ |
| DSD reaction ($k_{dsd}$) | $2 \times 10^{-3}$ $nM^{-1}$ $min^{-1}$ |
| Protocell size in radius (rprot) | 20 μm |
| Release signal from sender | 20 μM |
| Consumption capacity of receiver | 2 μM |

**Supplementary Table 5**. Simulation parameters for sender-transceiver system.

| | |
|---|---|
| Release rate ($k_{rel}$) | $2.7 \times 10^{-2}$ $min^{-1}$ |
| Permeability (P) | 200 μm $min^{-1}$ |
| Diffusion coefficient of DNA in buffer (d) | $2.2 \times 10^{3}$ $μm^{2}$ $min^{-1}$ |
| DSD reaction ($k_{dsd}$, $k_{fuel}$) | $2 \times 10^{-3}$ $nM^{-1}$ $min^{-1}$ |
| Leak reaction ($k_{leak}$) | $6 \times 10^{-6}$ $nM^{-1}$ $min^{-1}$ |
| Protocell size in radius (rprot) | 20 μm |
| Release signal from sender | 10 μM |
| Consumption capacity of transceiver | 1 μM |
| Good fuel | 0.08 μM |
| Bad Fuel | 0.02 μM |

**Supplementary Table 6**. Simulation parameters for Boolean AND gate system.

| | |
|---|---|
| Release rate ($k_{rel}$) | $2.7 \times 10^{-2}$ $min^{-1}$ |
| Permeability (P) | 200 μm $min^{-1}$ |
| Diffusion coefficient of DNA in buffer (d) | $2.2 \times 10^{3}$ $μm^{2}$ $min^{-1}$ |
| Forward DSD reaction ($k_f$) | $2 \times 10^{-3}$ $nM^{-1}$ $min^{-1}$ |
| Backward DSD reaction ($k_b$) | 0.6 $min^{-1}$ |
| Protocell size in radius (rprot) | 20 μm |
| Release signal from senders | 40 μM |
| Consumption capacity of transceiver | 1 μm |

**Supplementary Movie (separate file).**

**Movie 1**. Photo-cleavage of gate complex inside a sender protocell (Fig. 1c). Time lapse confocal microscopy recording the photo-cleavage of gate complex $\mathbf{F_1Q_1}$ inside a sender protocell upon laser irradiation (405 nm, 2 h). The left channel shows the FITC fluorescence (green) and the right channel shows the Cy5 fluorescence (red). The cleavage of PC linker results in dissociation of quencher-labeled fragment **B** indicated by an increase in Cy5 fluorescence. Sender protocell was prepared using 10 μM streptavidin.

**Movie 2**. Photo-cleavage of gate complex inside a sender protocell and activation of receiver protocells (Fig. 1d). Time lapse confocal microscopy recording the photo-cleavage of gate complex $\mathbf{F_1Q_1}$ inside a sender protocell upon laser irradiation (405 nm, 2 h) and the activation of gate complex $\mathbf{F_2Q_2}$ localized in receiver protocells. The left channel shows the FITC fluorescence (green) and the right channel combines the Cy5 (red) and Alexa546 (yellow) fluorescence. The increase in Cy5 fluorescence confirms the cleavage of PC linker on $\mathbf{F_1Q_1}$ in sender protocell and Alexa546 fluorescence associated with the activation in receiver protocells. Sender and receiver protocells were prepared using 10 μM and 4 μM streptavidin respectively.
